# The human brain vasculature shows a distinct expression pattern of SARS-CoV-2 entry factors

**DOI:** 10.1101/2020.10.10.334664

**Authors:** Moheb Ghobrial, Jason Charish, Shigeki Takada, Taufik Valiante, Philippe P. Monnier, Ivan Radovanovic, Gary D. Bader, Thomas Wälchli

**Author notes:** These authors contributed equally (Philippe P. Monnier, Ivan Radovanovic, Gary D. Bader). Corresponding author: Thomas Wälchli Group of CNS Angiogenesis and Group of Brain Vasculature and Neurovascular link Perivascular Niche Division of Neurosurgery Division of Neurosurgery, Neuroscience Center Zurich, and Division Krembil Neuroscience Institute of Neurosurgery University and University Hospital Zurich Toronto Western Hospital, University of Toronto Frauenklinikstrasse 10 60 Leonard Avenue CH-8057 Zürich / CH-8091 Zürich, Switzerland Toronto, Ontario, M5T 2S8.

## Abstract

A large number of hospitalized COVID-19 patients show neurological symptoms such as ischemic- and hemorrhagic stroke as well as encephalitis, and SARS-CoV-2 can directly infect endothelial cells leading to endotheliitis across multiple vascular beds. These findings suggest an involvement of the brain- and peripheral vasculature in COVID-19, but the underlying molecular mechanisms remain obscure. To understand the potential mechanisms underlying SARS-CoV-2 tropism for brain vasculature, we constructed a molecular atlas of the expression patterns of SARS-CoV-2 viral entry-associated genes (receptors and proteases) and SARS-CoV-2 interaction partners in human (and mouse) adult and fetal brain as well as in multiple non-CNS tissues in single-cell RNA-sequencing data across various datasets. We observed a distinct expression pattern of the cathepsins B (CTSB) and -L (CTSL) - which are able to substitute for the ACE2 co-receptor TMPRSS2 - in the human vasculature with CTSB being mainly expressed in the brain vasculature and CTSL predominantly in the peripheral vasculature, and these observations were confirmed at the protein level in the Human Protein Atlas and using immunofluorescence stainings. This expression pattern of SARS-CoV-2 viral-entry associated proteases and SARS-CoV-2 interaction partners was also present in endothelial cells and microglia in the fetal brain, suggesting a developmentally established SARS-CoV-2 entry machinery in the human vasculature. At both the adult and fetal stages, we detected a distinct pattern of SARS-CoV-2 entry associated genes’ transcripts in brain vascular endothelial cells and microglia, providing a potential explanation for an inflammatory response in the brain endothelium upon SARS-CoV-2 infection. Moreover, CTSB was co-expressed in adult and fetal brain endothelial cells with genes and pathways involved in innate immunity and inflammation, angiogenesis, blood-brain-barrier permeability, vascular metabolism, and coagulation, providing a potential explanation for the role of brain endothelial cells in clinically observed (neuro)vascular symptoms in COVID-19 patients. Our study serves as a publicly available single-cell atlas of SARS-CoV-2 related entry factors and interaction partners in human and mouse brain endothelial- and perivascular cells, which can be employed for future studies in clinical samples of COVID-19 patients.

## INTRODUCTION

The coronavirus disease 2019 (COVID-19) is caused by severe acute respiratory syndrome coronavirus 2 (SARS-CoV-2)^1^, which was first detected in Wuhan, China in December 2019^2^. Since, the virus has spread worldwide causing a global pandemic with important medical, social, and economic consequences^3,4^. An increasing number of reports describe neurological symptoms in COVID-19 patients, as the SARS-CoV-2 pandemic continues to spread around the world^5–32^. Mao and colleagues recently reported that over a third of hospitalized patients affected by COVID-19 showed neurological symptoms such as acute cerebrovascular events including ischemic and hemorrhagic strokes, impaired consciousness, and muscle injury, and that those neurological manifestations were even more common in patients with severe infection as classified according to their respiratory status^33^. Others reported COVID-19 patients with clinical presentations of large-vessel occlusions resulting in ischemic stroke in young patients^32,34^, acute hemorrhagic necrotizing encephalopathy^35^, encephalitis^29,36^, and headaches and dizziness^37^. These cerebrovascular manifestations may be due to impairment of the distinct features of the brain vasculature, namely the blood-brain-barrier and the neurovascular unit, a functional entity composed of endothelial cells and perivascular cells^38–41^. Moreover, neurological/encephalitic manifestations similar to those described in adults such as white matter injury (bilateral gliosis of the deep white periventricular- and subcortical matter), lethargy, hypertonia, hyperexcitability, and high-pitched crying were also reported in human neonates born to SARS-CoV-2 infected mothers^42,43^. The fetuses/neonates were infected via transplacental transmission of SARS-CoV-2 leading to placental inflammation and neonatal viremia, and these neurological symptoms can be associated to cerebral vasculitis^42,43^. These observations together with the reported “vascular symptoms” like Kawasaki syndrome are especially interesting given that the brain- and peripheral vasculature are established during fetal and postnatal developmental periods^42–48^.

Recent evidence supports that endothelial cells are at the center stage of the disease as the SARS-CoV-2 virus can cause endotheliitis^49^, which is suggested to be causative for multi-organ failure observed in some COVID-19 patients^32^, and can be targeted *in vitro* and in mice *in vivo*^50,51^. For instance, human recombinant soluble ACE2 (hrsACE2) inhibits infection of human blood vessel organoids and human kidney organoids *in vitro*, suggesting that targeting SARS-CoV-2-blood vessel interaction may prevent deleterious effects of the virus on the neurovascular unit and the BBB^50^.

SARS-CoV-2 (as well as SARS-CoV-1) enters cells via binding of the spike (S) protein on the cell surface to its entry receptor angiotensin converting enzyme 2 (ACE2)^52^. Subsequently, the S protein is primed by the SARS-CoV-2 entry-associated proteases TMPRSS2, CTSB (the gene encoding for human Cathepsin B), and CTSL (the gene encoding for human Cathepsin L)^50,53,54^. CTSB and CTSL are SARS-CoV-2 entry-associated proteases and their activity are suggested to be able to substitute for TMPRSS2 protease activity indicating a potential alternative mechanisms in protein S protease priming^53,55–58^. The proteolytic separation of the S1 and S2 subunits, termed priming, provides the SARS-CoV S protein with the structural flexibility required for the membrane fusion reaction. Whereas CTSB cleaves the SARS-CoV-2 spike protein, it does not process the spike protein in SARS-CoV-1^59^. On the other hand, CTSL appears to process the spike protein of both viruses with, however, a higher efficacy for SARS-CoV-1 than SARS-CoV-2^59^. Other cathepsins important for SARS-CoV viral entry include CTSA, CTSC, CTSD, CTSE, CTSF, CTSG, CTSH, CTSK, CTSO, CTSS, CTSV, CTSW, CTSZ, whereas other viral receptors/receptor-associated enzymes include ANPEP (HCoV-229E), DPP4 (MERS-CoV), ST6GAL1 (Influenza), ST3GAL4 (Influenza) (see https://www.covid19cellatlas.org/).

The ACE2 receptor is expressed in different cell types in several peripheral organs, including the lung, heart, kidney, liver, spleen, colon, and intestine^60,61^. Interestingly, in the brain, ACE2 is predominantly expressed in vascular endothelial- and smooth muscle cells of blood vessels, as evidenced by immunohistochemistry^60,61^. Recently, 332 high-confidence SARS-CoV-2- human protein-protein interactions (PPIs, proteins that are physically associated with SARS-CoV-2 proteins) were identified using affinity-purification mass spectrometry (AP-MS), thereby defining a set of potentially druggable SARS-CoV-2 interaction partners^62^. Despite these indications, the expression patterns of the ACE2 receptor, the entry-associated proteases, and the SARS-CoV-2 interaction partners in endothelial- and perivascular cells of the human and mouse brain vasculature have not been investigated in detail so far.

## RESULTS

### The brain and the brain vasculature/vascular endothelium show high expression of the SARS-CoV-2 protease Cathepsin B (CTSB) but not of the Transmembrane protease, serine 2 (TMPRSS2)

ACE2 is expressed in vascular endothelial- and smooth muscle cells of the brain, as previously shown at the protein level using immunohistochemistry^61^, but the pattern of expression of ACE2 and SARS-CoV-2 entry associated genes (https://www.covid19cellatlas.org/) in the human brain and peripheral organs remains incompletely understood. To address the expression patterns of the SARS-CoV-2 entry receptor, SARS-CoV-2 entry-associated proteases, SARS-CoV-2 interaction partners, and other genes involved in SARS-CoV-2 pathogenesis such as cathepsins and viral receptors/receptor-associated enzymes in the human and mouse brain vasculature at cellular resolution, and because the human (brain) vasculature is established during fetal/embryonic and postnatal development, we analyzed various human fetal and adult single-cell RNA sequencing (scRNA-seq) datasets from healthy donors generated in the Human Cell Landscape compendium and other resources^63^. To address SARS-CoV-2 entry-associated gene expression patterns in the human brain and in peripheral organs at the single-cell level, we referred to a recently published molecular atlas of the human cell landscape utilizing adult and fetal human samples covering all major human organs and a total of 42 human tissue types (we analyzed the 42 out of 60 human tissues which included endothelial cells) and 471,942 single cells (283,355 adult-cells from 26 tissues/organs and 188,587 fetal cells from 16 tissues/organs)^63^.

We visualized the data using t-distributed stochastic neighbor embedding (t-SNE) and computed differential gene expression analyses for each specific organ in adult (26 organs/tissues, ca. 283’355 cells) and fetal (16 organs/tissues, ca. 188’587 cells) human organs/tissues (Figure 1a,b).

**Figure 1.**
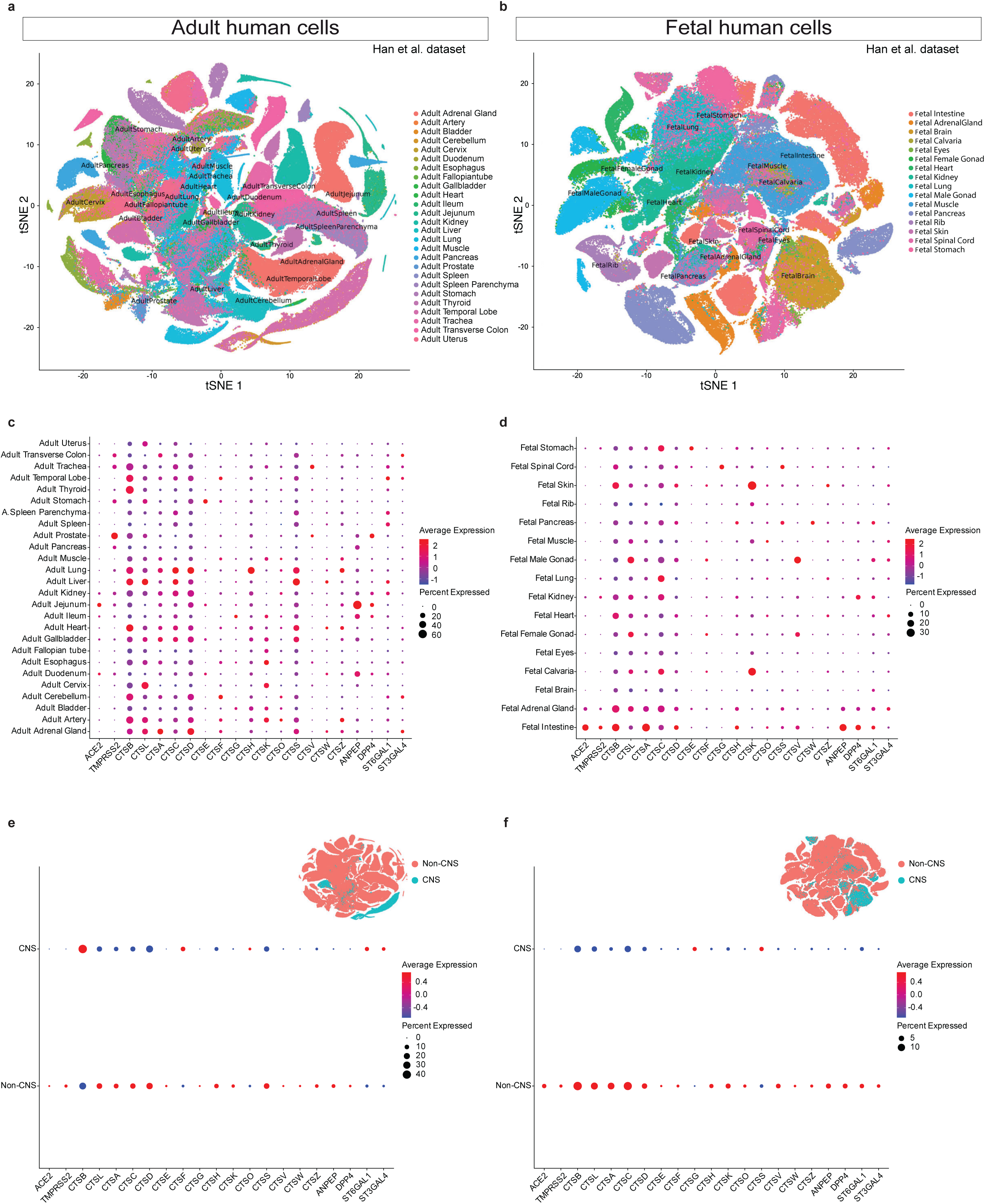
Expression of SARS-Cov-2 entry associated genes in human adult and fetal CNS and non-CNS tissues. **a**,**b**, t-SNE plot of 283’355 and 188’587 single cells of adult (**a**) and fetal (**b**) tissues respectively from the human cell landscape scRNA-seq dataset^63^. In the *t*-SNE plots, tissues are labelled by different colors, and every tissue represents an individual cluster; 26 adult and 16 fetal tissues were analyzed. **c,d**, RNA expression of the SARS-CoV-2 entry receptor ACE2, and of entry-associated proteases: TMPRSS2, CTSB, CTSL, other cathepsins: CTSA, CTSC, CTSD, CTSF, CTSG, CTSH, CTSK, CTSO, CTSS, CTSV, CTSW, CTSZ,and other viral receptors/receptor-associated enzymes: ANPEP, DPP4, ST6GAL1, ST3GAL4 in adult (**c**) and fetal (**d**) cells from the human cell atlas scRNA-seq dataset across different tissues. **e**,**f**, RNA expression of SARS-CoV-2 entry receptor, entry-associated proteases, other cathepsins and other viral receptors/receptor-associated enzymes in CNS vs non-CNS cells of adult (**e**) and fetal (**f**) tissues from the human cell atlas. The adult and fetal datasets were obtained from existing publically available sources and tissue clustering and nomenclature were retained based on the author annotation^63^. Expression values were normalized, log transformed and scaled. For results in the dot plots, the dot size represents the proportion of cells within the respective tissue type expressing the gene, and the dot color represents the average gene expression level within the particular tissue type.

In the adult, the expression of ACE2 was generally low in all tissues, in agreement with the previously reported low expression level of ACE2 in scRNA seq datasets^64^, demonstrating the highest expression in the jejunum, duodenum, kidney, and ileum (Figure 1c). TMPRSS2 expression displayed a somewhat broader distribution than ACE2 (as reported in Sungnak et al., *Nat Med*, 2020^64^) with the highest expression in the prostate, followed by trachea, colon, stomach, pancreas, kidney, jejunum, duodenum, and lung (Figure 1c). In the temporal lobe and the cerebellum, both ACE2 and TMPRSS2 demonstrated low expression (Figure 1c). As mentioned above, CTSB and CTSL can substitute TMPRSS2 protease activity. Therefore, to address potential alternative mechanisms for protein S protease priming and thus SARS-CoV-2 entry into cells, we hereafter mainly focus on CTSB and CTSL. CTSB and CTSL were highly expressed in various organs, and CTSB showed the highest expression in the thyroid, heart, liver, and temporal lobe, followed by the lung, artery, and cerebellum (Figure 1c). CTSL, in contrast, showed low expression in the temporal lobe and cerebellum (Figure 1c), and CTSB showed higher expression than CTSL in both the temporal lobe and the cerebellum (Figure 1c). In the fetus, the expression of both ACE2 and TMPRSS2 was low in most tissues, displaying, however highest expression values in the intestine and adrenal gland (Figure 1d); ACE2 and TMPRSS2 showed very similar expression patterns across all organs, as revealed by dot plots (Figure 1d). Similar to the adult, CTSB and CTSL were variably but overall highly expressed in all tested organs, and CTSB showed higher expression than CTSL in both the spinal cord and brain (Figure 1d).

To examine differential expression patterns of the brain/CNS and the periphery (non-CNS), we compared the brain/CNS (pool of temporal lobe and cerebellum in the adult, and pool of brain and spinal cord in the fetus) to a pool of all peripheral organs (Figure 1e,f). Interestingly, ACE2 and TMPRSS2 showed a higher expression in the periphery in both the adult and the fetus (Figure 1e,f). In the adult, CTSB was higher expressed in the brain/CNS, both in intensity and percent of expression, whereas CTSL showed higher expression in the periphery (Figure 1e). In the fetus, both CTSB and CTSL showed a higher expression in the periphery (Figure 1f).

CTSB also displayed a higher expression than CTSL in the adult and fetal mouse brain/CNS as compared to the periphery, thereby confirming the observations made in humans (Extended Data Figure 3e-f, 3k-l). Next, to address the cellular expression patterns of SARS-CoV-2 entry-associated genes, we analyzed all cell types of all organs in both the adult and fetus (Extended Data Figure 1a-x and Extended Data Figure 2a-p). In the adult and fetal brain, CTSB showed a higher expression than CTSL in all cell types (including the endothelium) with the exception of smooth muscle cells, astrocytes, and epithelial cells in the adult (Extended Data Figure 1d). In the fetus, CTSB showed a higher expression in endothelial cells, macrophages, and microglia and CTSL was higher expressed in stromal cells and fibroblasts whereas CTSB and CTSL showed a similar expression in all other cell types (Extended Data Figure 2b).

Together, these data indicate that in the adult, CTSB shows a higher expression pattern in the CNS as compared to the periphery, whereas CTSL seems to display an inversed expression pattern with a predominant expression in the periphery.

The brain vasculature is established during fetal and postnatal brain development and its endothelial cells display very specific characteristics distinguishing it from vascular beds of peripheral organs (Figure 2a), as we and others have described previously^38,39,41^. First, to examine SARS-CoV-2 entry-associated gene expression patterns in all (CNS- and periphery) endothelial cells at the fetal and adult stage, we computationally extracted adult and fetal 10 endothelial cells (Figure 2a). Interestingly, the expression patterns were very similar between the developing fetal and mature adult stage (with a generally higher expression in the adult), indicating a developmentally established endothelial expression pattern of SARS-CoV-2 entry-associated genes (Figure 2b). Next, to address potential differences between CNS- and non-CNS endothelial cells, we compared the expression patterns of SARS-CoV-2 receptor, - proteases, -and other entry-associated genes in endothelial cells of the human brain and of peripheral organs (Figure 2c,d). The expression of ACE2 was generally low in endothelial cells of all tissues/organs (Figure 2e,f), again in agreement with the previously reported low expression level of ACE2 in scRNA seq datasets^64^. In the adult, ACE2 was, highest in the duodenum, ileum, transverse colon, gallbladder, and kidney (Figure 2e, Extended Data Figure 1a-x), as previously described^54,65,66^, whereas the fetal intestine, heart, adrenal gland and kidney showed the highest ACE2 expression (Figure 2f, Extended Data Figure 2a-p) Similarly, TMPRSS2 expression in endothelial cells was generally low in all adult and fetal tissues displaying – similar to the situation observed when looking at all cells (see above) - the highest expression in the adult prostate, pancreas, and duodenum (Figure 2e, Extended Data Figure 1a-x), and in the fetal intestine, kidney, and lung, but was expressed in non-endothelial cells across multiple tissues, (with a broader distribution than ACE2), as recently shown (Figure 2f, Extended Data Figure 2a-p). At the adult stage, CTSB was highly expressed in endothelial cells of the liver, gallbladder, and adrenal gland, followed by the endothelial cells of the temporal lobe, heart and the cerebellum (Figure 2e), whereas at the fetal stage, the brain endothelial cells showed the highest CTSB expression of all organs/tissues, followed by the endothelial cells of the skin, heart, spinal cord, and others (Figure 2f). In human endothelial cells of the adult temporal lobe and the adult cerebellum as well as of the fetal brain, CTSB was the highest expressed SARS-CoV-2 entry-associated protease, (Figure 2e,f) whereas CTSB and CTSC showed a very similar expression level in 11 the endothelial cells of the fetal spinal cord (Figure 2f), further suggesting the relative importance of CTSB in the developing and adult CNS vasculature. Other cathepsins that showed higher expression in the adult and fetal brain endothelium included CTSA, CTSD, and as well as CTSC, CTSF, and (Figure 2e,f).

**Figure 2.**
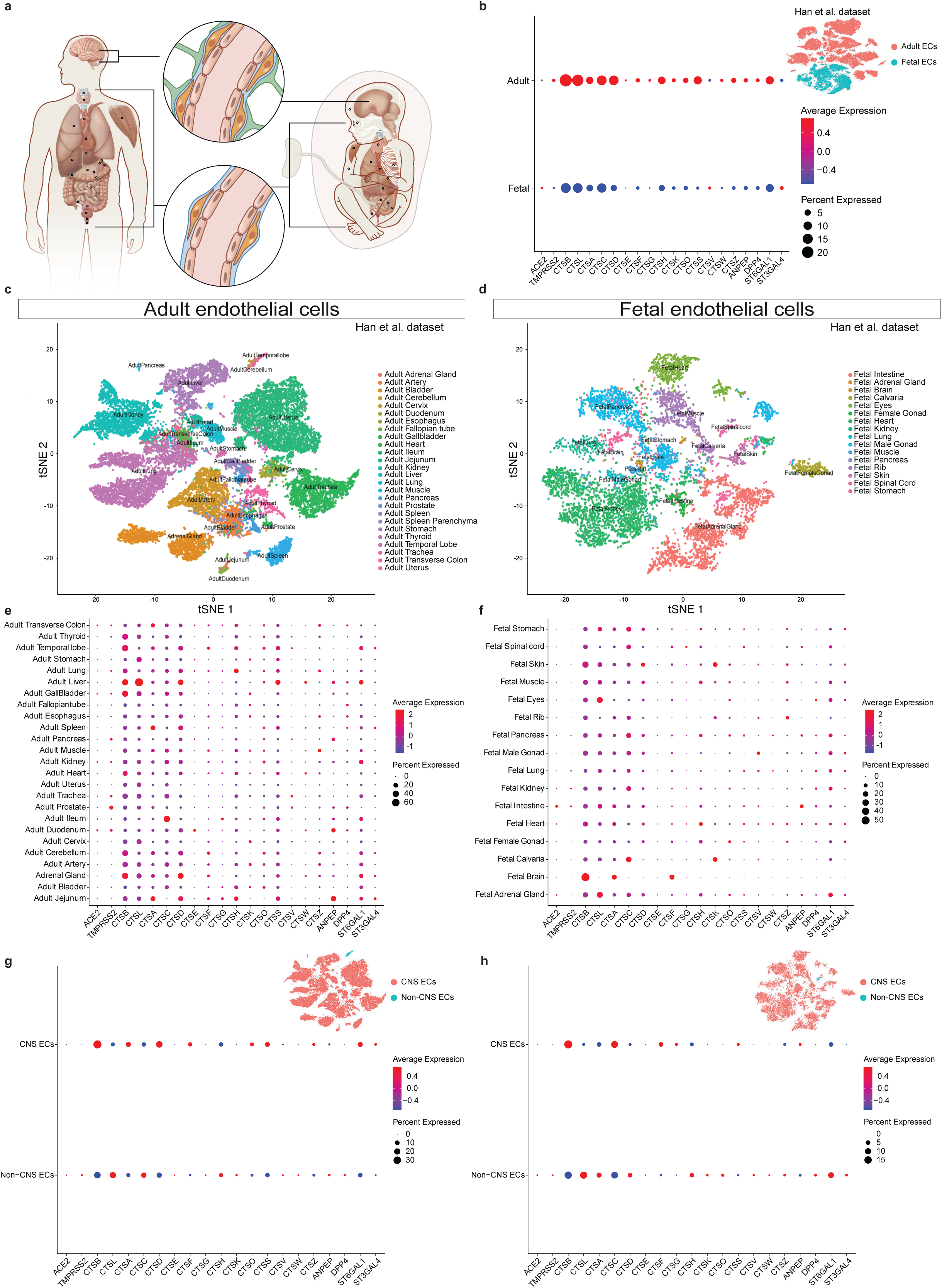
Expression of SARS-Cov-2 entry associated genes in human adult and fetal CNS and non-CNS endothelial cells. **a**, Schematic illustration depicting major tissues from which endothelial cells were computationally extracted and analysed referring to single cell RNA-seq datasets from adult (left) and fetal (right) developmental stages. **b**, Dotplot comparing expression of SARS-CoV-2 entry receptor, entry-associated proteases, other cathepsins, and other viral receptors/receptor-associated enzymes in adult and fetal endothelial cells that were computationally extracted from tissues illustrated in (**a**). **c**,**d**, t-SNE plot of 22’993 and 9’022 endothelial cells of adult (**c**) and fetal (**d**) tissues respectively from the human cell landscape scRNA-seq dataset^63^. In the *t*-SNE plot, tissues are labelled by different colors; and every tissue represent an individual cluster; endothelial cells were extracted from 26 adult and 16 fetal tissues. **e**,**f**, RNA expression of SARS-CoV-2 entry receptor, entry-associated proteases (TMPRSS2, CTSB, CTSL), other cathepsins, and other viral receptors/receptor-associated enzymes in adult (**e**) and fetal (**f**) endothelial cells from the human cell landscape scRNA-seq dataset across different tissues^63^ . **g**,**h**, RNA expression of SARS-CoV-2 entry receptor, entry-associated proteases (TMPRSS2, CTSB, CTSL), other cathepsins and other viral receptors/receptor-associated enzymes in CNS and non-CNS endothelial cells of adult (**g**) and fetal (**h**) tissues from the human cell atlas.

When comparing the pooled CNS/brain (temporal lobe and cerebellum) against all peripheral endothelial cells, these observations were confirmed (Figure 2g,h). Moreover, we observed that in both the adult and fetus, CTSB showed a higher expression in the CNS as compared to the periphery, whereas CTSL showed the opposite, namely a higher expression in the periphery as compared to the CNS (Figures 2g,h), and these observations were confirmed in most of the analyzed datasets (Extended Data Figure 1a-x, Extended Data Figure 2a-p). Interestingly, CTSB was the highest expressed SARS-CoV-2 entry associated gene in adult and fetal brain endothelial whereas CTSL showed the highest expression of all SARS-CoV-2 entry-associated genes in adult and fetal peripheral endothelial (Figure 2g,h). Together, these data suggest that the distinct expression pattern of SARS-CoV-2 proteases and associated enzymes in the CNS is developmentally established. .

### Brain endothelial cells and microglia display high expression of CTSB but not of the Transmembrane protease, serine 2 (TMPRSS2)

The brain vasculature is comprised of endothelial cells and perivascular cells of the neurovascular unit including neurons, pericytes, astrocytes, and immune cells such as microglia and macrophages^38,39,67–69^ (Figure 3a). To examine the expression patterns of SARS-CoV-2 entry factor transcripts in the neurovascular unit and thereby address potential entry mechanisms of SARS-CoV-2, we next addressed brain endothelial- and perivascular cells (Figure 3a). First, looking at all brain cells (endothelial- and perivascular cells pooled 12 together), we observed that CTSB showed the highest transcript expression of SARS-CoV-2 entry associated genes in both the adult and fetal brain, and that the SARS-CoV-2 entry associated genes were more highly expressed in the adult than in the fetal stage while displaying certain similarity (Figure 3b). Next, we compared the SARS-CoV-2 entry-associated gene expression patterns between endothelial- and perivascular cells at the adult and fetal stages (Figure 3c,d). Of note, ACE2 and TMPRSS2 showed low expression levels throughout all adult and fetal datasets, consistent with their low expression patterns described above (Figure 3e,f, and Extended Data Figure 5a-j).

**Figure 3.**
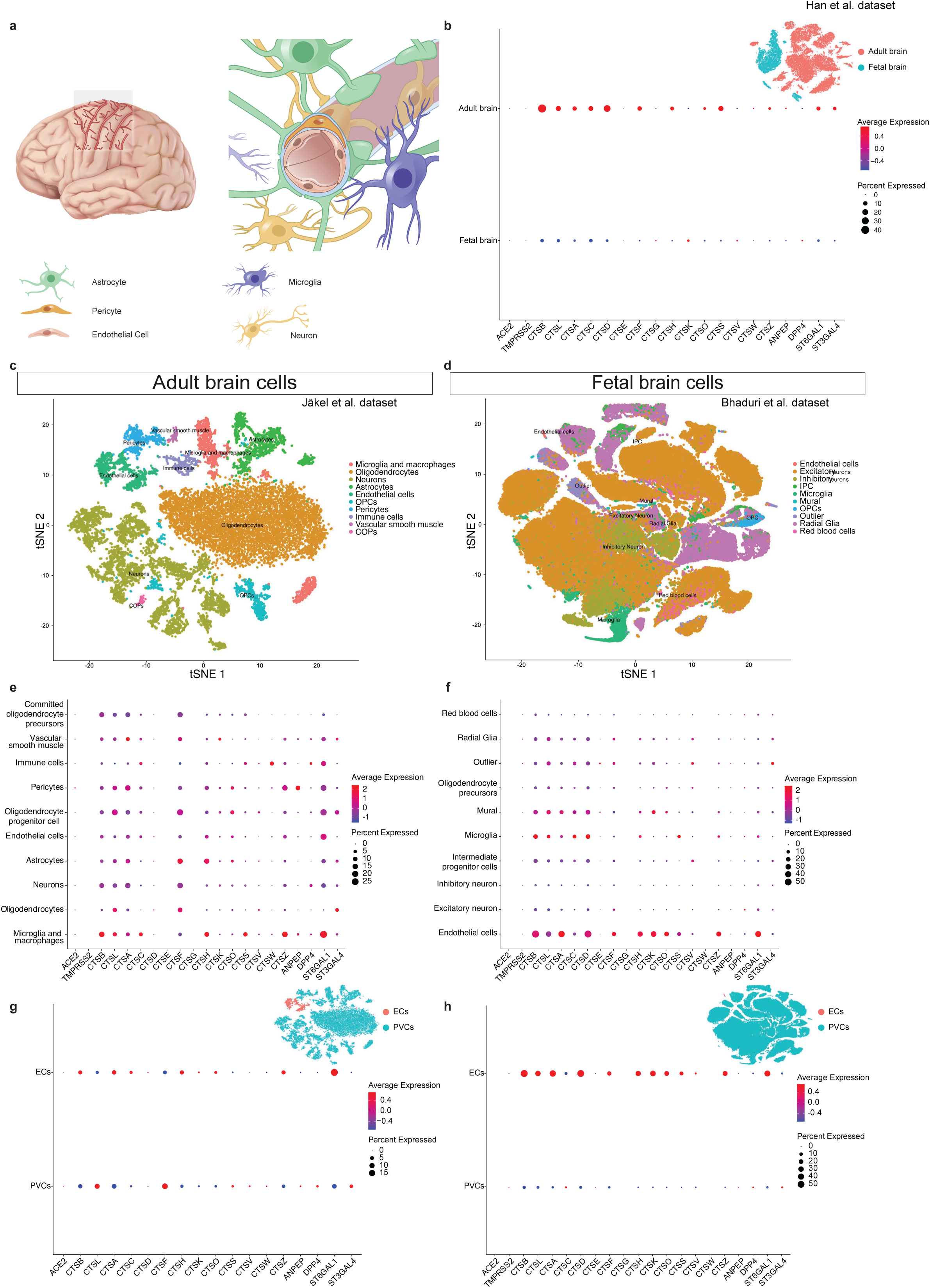
Expression of SARS-Cov-2 entry associated genes in human adult and fetal brain endothelial-and perivascular cells. **a** Scheme illustrating brain blood vessels running within cortical sulci (left). The neurovascular unit (NVU) is composed of endothelial cells and of perivascular cell types such as pericytes, astrocytes, neurons, and microglia. **b**, Dotplot comparing expression of SARS-CoV-2 entry receptor, entry-associated proteases (TMPRSS2, CTSB, CTSL), other cathepsins, and other viral receptors/receptor-associated enzymes in adult and fetal brain cells of the human cell landscape scRNA-seq dataset^63^. **c**,**d**, t-SNE plot of single cells of adult (**c**) and fetal (**d**) brain tissue from the Jäkel et al.^70^ and Bhaduri et al.^110^ datasets respectively. In the *t*-SNE plot, cell types are labelled by different colors. **e**,**f**, RNA expression of SARS-CoV-2 entry receptor, entry-associated proteases, other cathepsins and other viral receptors/receptor-associated enzymes in adult (**e**) and fetal (**f**) brain cells from Jäkel et al.^70^ and Bhaduri et al.^110^ datasets respectively scRNA-seq datasets. **g,h**, RNA expression of SARS-CoV-2 entry receptor, entry-associated proteases, other cathepsins and other viral receptors/receptor-associated enzymes in endothelial cells vs perivascular cells of adult (**g**) and fetal (**h**) brain cells from the Jäkel et al.^70^ and Bhaduri et al.^110^ datasets respectively.

In the adult, CTSB was highly expressed in endothelial cells and in microglia and macrophages, and it was amongst the highest expressed SARS-CoV-2 entry-associated genes across different datasets (Figure 3e-f and Extended Data Figure 5a-j). Namely, CTSB was either the highest expressed SARS-CoV-2 entry-associated genes (in 2 out of 6 datasets, namely Jäkel et al.^70^, Welch et al.^71^) or figured in the top five highest expressed SARS-CoV-2 entry-associated genes (in 6 out of 6 datasets, Jäkel et al.^70^, Welch et al.^71^, Han et al.^63^, Lake et al.^72^, Habib et al.^73^) (Figure 3g, Extended Data Figure 4a,c,e,g,i and Extended Data Figure 5 a,c,e,g,i).

To address whether this endothelial- and perivascular cell expression pattern was developmentally established, we referred to the human fetal scRNA seq datasets. Endothelial cells and microglia again showed a similar expression pattern of SARS-CoV-2 entry associated genes displaying high CTSB expression, similar to the adult situation (Figure 3f and Extended Data Figure 5b,d,f,h,j). CTSB (showed – to the exception of one out of six datasets (Polioudakis et al. dataset, in which endothelial cells showed a (Figure 3h, Extended Data Figure 4b,d,f,h,j and Extended Data Figure 5b,d,f,h,j)) – consistently highest expression in endothelial cells and microglia (Figure 3h, Extended Data Figure 4b,d,f,h,j and Extended Data Figure 5b,d,f,h,j), and these two cells types also both expressed CTSL, (Figure 3h, 13 Extended Data Figure 4b,d,f,h,j and Extended Data Figure 5b,d,f,h,j). Notably, SARS-CoV-2 entry associated genes showed a similar pattern of expression between the fetal and adult endothelial cells and microglia further supporting the aforementioned developmentally established expression pattern of SARS-CoV-2 entry associated genes.

Looking at the datasets with pooled perivascular cells we observed that within endothelial cells, CTSB consistently showed the highest expression of all SARS-CoV-2 entry-associated genes across all adult and fetal datasets examined (Figure 3g,h and Extended Data Figure 4a-j). Comparison of endothelial cells against a pool of all perivascular cells revealed that CTSB (showed a higher expression pattern in endothelial- as compared to perivascular cells in two out of six adult datasets (Jäkel et al.^70^, Welch et al.^71^) whereas CTSB was higher in the perivascular cells in the four other datasets (Han et al.^63^, Lake et al.^72^, Habib et al.^73^, Velmeshev et al.^74^) due to the high expression in microglia and macrophages (Figure 3g, Extended Data Figure 4a,c,e,g,i and Extended Data Figure 5a,c,e,g,i). CTSB displayed a higher endothelial- than perivascular expression in five out of six fetal datasets, with an inverse pattern in one sole dataset (La Manno et al.^75^) again due to the high microglial and macrophage CTSB expression (Figure 3h, Extended Data Figure 4b,d,f,h,j and Extended Data Figure 5b,d,f,h,j). Next, we wondered whether these observed CTSB/CTSL- and SARS-CoV-2 entry associated factor expression patterns were retained in *in vitro* brain organoid models which were recently employed to study SARS-CoV-2 invasiveness to the brain^76^. In a recent single cell RNA-seq datasets of brain organoids^77^, CTSB showed the highest expression in cells undergoing epithelial-to-mesenchymal transition (EMT), followed by endothelial-like and endothelial-like progenitor cells and neuronal-like progenitor cells (Extended Data Figure 6a-c). When comparing the expression patterns in endothelial-like cells to the pool of perivascular cells, CTSB was is higher in endothelial-like cells and CTSL was higher in 14 perivascular cells (Extended Data Figure 6c). These expression patterns were similar to the ones in the adult and fetal human brain datasets, indicating that brain organoids might constitute a model system to further explore the brain vascular-related observations made here Together, these data indicate that endothelial cells and perivascular cells (mainly microglia) of the fetal and adult CNS neurovascular unit express CTSB but not TMPRSS2, suggesting a potential alternative of how SARS-CoV-2 might enter the brain endothelium and neurovascular unit. Importantly, additional validation experiments need to be carried out to validate these findings at the protein level.

### CTSB is highest expressed in veins and capillaries in fetal and adult brain vasculature and correlates with multiple pathways involved in COVID-19 pathogenesis

To further characterize the SARS-CoV-2 entry factor transcript expression patterns in brain endothelial cells, we examined the different brain endothelial cell clusters according to their arterio-venous specification, as previously described in mouse brain^78–80^ (Figure 4a). We found that CTSB was expressed in multiple brain endothelial cell types including veins, capillaries, endothelial cells 1 (EC1), antigen presenting- and vein (antigen presenting) endothelial cells (Figure 4b). In all adult and fetal brain endothelial cell clusters, CTSB was amongst the highest expressed SARS-CoV-2 entry-associated transcripts (Figure 4a,b and Extended Data Figure 7a,b). In the adult brain vasculature, the veins cluster showed the highest average expression of CTSB, followed by EC1, capillaries, antigen-presenting veins, and antigen-presenting endothelial cells (Figure 4a,b), whereas CTSB showed the highest expression in fetal veins > capillaries > arteries > EC1 (Extended Data Figure 7a,b).

**Figure 4.**
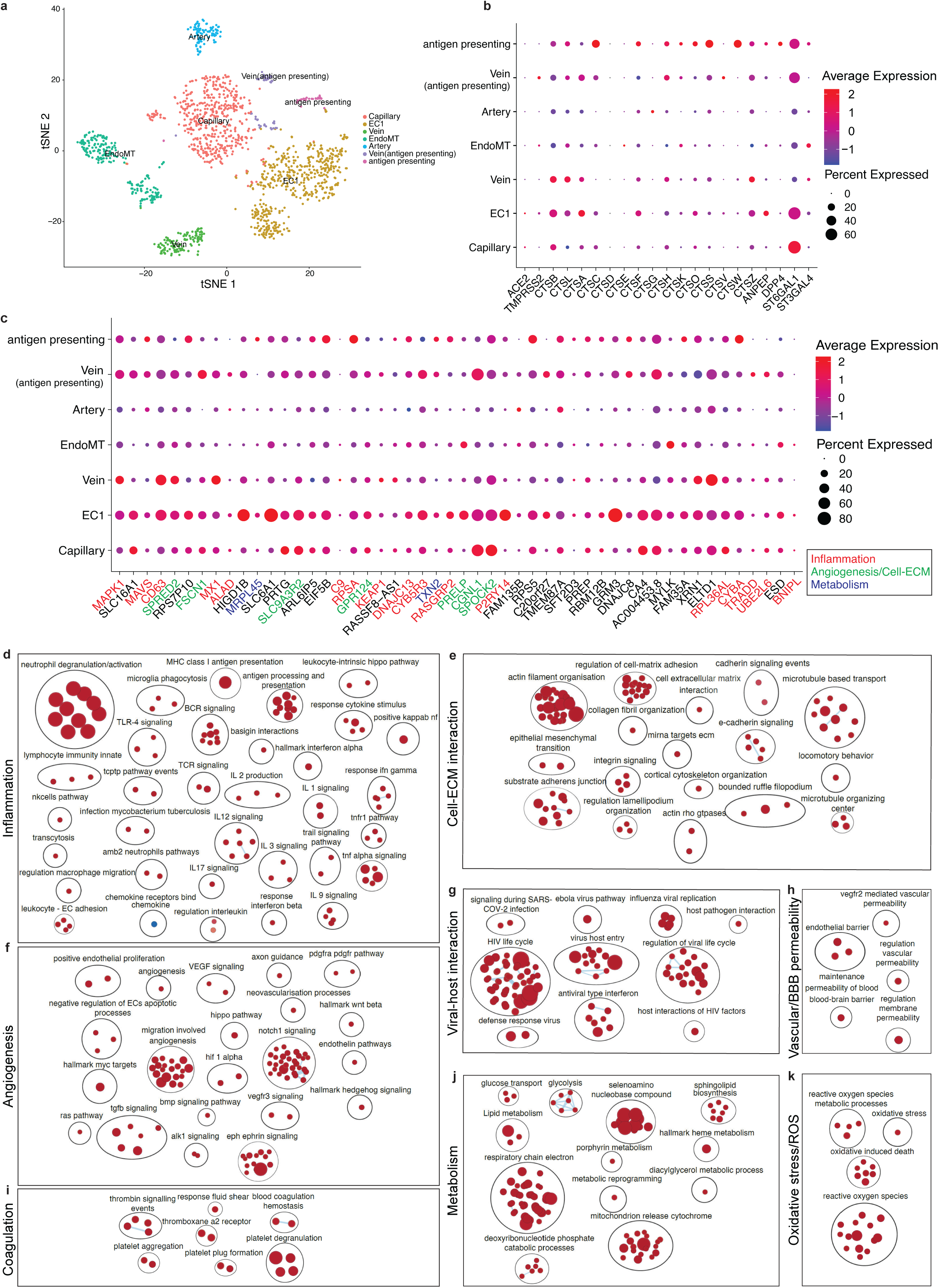
Expression of SARS-Cov-2 entry associated genes in human adult brain endothelial cells. **a,** t-SNE plot of 1’548 endothelial cells of adult brain tissue computationally extracted from the Velmeshev^74^ et al. dataset. In the *t*-SNE plot, endothelial cell sub-types are labelled by different colors, and every endothelial cell subtype represents an individual cluster. **b,** RNA expression of the SARS-CoV-2 entry receptor ACE2, entry-associated proteases (TMPRSS2, CTSB, CTSL), other cathepsins and other viral receptors/receptor-associated enzymes in adult brain endothelial cell sub-types from Velmeshev et al. dataset^74^. **c,** Brain endothelial expression of the top 50 genes correlated with CTSB expression based on Spearman’s correlation analysis performed on endothelial cells within the Velmeshev et al. dataset^74^. The colored gene names represent genes that are immune-associated/inflammatory response related (red), angiogenesis/cell-ECM interaction related (green) and metabolism related (blue). For gene expression results in the dot plots, the dot size represents the proportion of cells within the respective cell type expressing the gene and the color represents the average gene expression level within the particular endothelial cell sub-type. **d-k,** Enrichment map visualizing the GSEA pathway analysis results performed on the ranked list of the CTSB correlating genes, the top significantly enriched pathways included inflammation (**d**), angiogenesis (**f**), coagulation (**i**), cell-extracellular matrix interaction (**e**), viral-host interaction (**g**), vascular metabolism (**j**), blood-brain-barrier permeability (**h**), and reactive oxygen species (ROS) (**k**) in adult brain endothelial cells of the Velmeshev et al. dataset^74^.

Notably, recent studies using single-cell transcriptomics in human and mouse tissues revealed endothelial clusters in a variety of tissues including the human bladder, and kidney (and others) as well as the mouse and human lung expressing MCH class II genes such as HLA-DPA1 and HLA-DRA involved in MHC class II- mediated antigen processing, loading and presentation^80^.

These observations of professional antigen-presenting signatures of endothelial cells suggested potential immune functions of endothelial cells, and CTSB might be presented at the cell-surface upon SARS-CoV-2-infection of brain endothelial cells.

These results suggest that CTSB expression in different endothelial clusters including antigen-presenting endothelial cells is developmentally established and might be reactivated in pathological conditions, for instance upon endothelial cell infection with SARS-CoV-2.

### CTSB correlates with genes and pathways involved in inflammation, angiogenesis, coagulation, and blood-brain-barrier permeability in brain endothelial cells

Next, we wondered which genes were associated with CTSB in brain endothelial cells. We performed Spearman’s correlation analysis of CTSB in brain endothelial cells within the datasets. The observed correlation coefficients of the top 50 CTSB-correlated genes were relatively low, which might be due to dropout effects and technical noise in the scRNA seq datasets, and the top 50 CTSB correlated genes showed a slight predominance for the capillary-venous side, consistent with that of CTSB (Figure 4c). Notably, we observed various genes associated with immune functions/inflammation including innate and antiviral immune functions in the rank list, including MAPK1, MAVS, CD63, MX1, ALAD, C9, RPSA, KEAP1, DNAJC12, CYB5R3, and others (Figure 4c, red and Supplementary Table 1). Expression of most of these inflammatory genes is highest in venous endothelial cells, and for some in vein antigen-presenting endothelial cells, antigen-presenting endothelial cells, and EC1 (Figure 4c), indicating immunity functions for these endothelial cell clusters. The relatively high expression of inflammatory-related genes in antigen-presenting (venous) endothelial cells is plausible whereas the relative preference for the venous side remains to be further explored.

Moreover, genes involved in angiogenesis and cell-extracellular matrix interactions including SPRED2, FSCN1, SLC9A3R2, GPR124, and others, and were also abundantly present in the rank list and were highest in capillaries, veins, and EC1 (Figure 4c, green and Supplementary Table 1). We also observed several genes associated with endothelial glucose- and fatty acid metabolism, namely MRPL45 and TXN2, which were highest in EC1 and veins (Figure 4c, blue and Supplementary Table 1).

We observed a similar expression pattern of the top 50 CTSB-correlated genes in the fetal brain vascular endothelial cells (Extended Data Figure 7c and Supplementary Table 2) with, however, higher correlation coefficients (<0.39). The top 50 CTSB-correlated genes were consistent with that of CTSB, with a predominant expression pattern in veins > capillaries > arteries > EC1s (Extended Data Figure 7c).

Similar to the adult situation, genes associated with immune functions/inflammation including ESAM, IFITM3, PECAM1, CD81, TSC22D3, CSRP2, and others (Extended Data Figure 7c, red and Supplementary Table 2) were high in the rank list and showed highest expression in veins.

Genes involved in angiogenesis and cell-extracellular matrix interactions including SPARC, RHOA, TMSB10, VIM, CLDN5, and others, as well as genes associated with endothelial glucose- and fatty acid metabolism, namely SEPW1 and POMP were also abundantly present and were all highest in veins, followed by capillaries, arteries, and EC1 (Extended Data Figure 7c, green and Supplementary Table 2).

We next performed pathway analysis using GSEA and Cytoscape on the CTSB-correlated genes. Notably, the top regulated pathways included inflammation, angiogenesis, coagulation, cell-extracellular matrix interaction, viral-host interaction, vascular metabolism, blood-brain-barrier permeability, and reactive oxygen species (ROS) in both the adult and the fetal brain endothelium (Figure 4d-k and Extended Data Figure 7d-k). Notably, all these pathways and mechanisms have been linked to clinical observations of COVID-19 patients with neurological- and vascular symptoms^30,49,81,82^.

Together, these data reveal that CTSB is highly expressed in various endothelial cell clusters of the fetal and adult human brain and that pathways downstream of CTSB might provide a suggestive explanation of some of the neurovascular symptoms observed in COVID-19 patients.

### The human brain vascular endothelium displays high expression of intracellular SARS-CoV-2 interaction partners

Upon entry of the SARS-CoV-2 virus into the host cell, the virus interacts with multiple intracellular proteins^62^. Therefore, we next addressed the expression of the recently described 332 SARS-CoV-2 interaction partners (= protein-protein interactions (PPIs) = proteins that are physically associated with SARS-CoV-2 proteins) in the different adult and fetal organs (Extended Data Figure 8a-d). In the adult, SARS-CoV-2 interactions partners were highly expressed in trachea, cerebellum, artery, kidney, and temporal lobe (Extended Data Figure 8e), and the pooled analysis showed a higher expression in CNS versus non-CNS organs (Extended Data Figure 8f). In the fetus, SARS-CoV-2 interactions partners were highly expressed in fetal male gonad, adrenal gland, brain conjointly with heart, intestine and kidney (Extended Data Figure 8g). In the pooled analysis, however, due to the relatively low expression in the fetal spinal cord the expression was higher in non-CNS- as compared to CNS organs (Extended Data Figure 12h).

Next, we focused on the endothelial cells in the different adult and fetal organs (Extended Data Figure 8i-l). In the adult, endothelial cells of the temporal lobe and the cerebellum were among the organs/tissue displaying high expression of SARS-CoV-2 interaction partners, together with endothelial cells from artery, cervix, trachea, uterus, kidney, muscle and spleen (Extended Data Figure 8m). The relatively high expression in CNS versus non CNS endothelial cells was further revealed in the pooled analysis (Extended Data Figure 8n). At the fetal stage, brain endothelial cells showed the highest expression of SARS-CoV-2 interaction partners among all organs/tissues, followed by endothelial cells from male gonad, adrenal gland, pancreas, heart, and muscle (Extended Data Figure 8o). Again, due to the relatively low expression in fetal spinal cord endothelial cells (Extended Data Figure 8o), however, the pooled analysis revealed a higher expression in non-CNS- as compared to CNS endothelial cells (Extended Data Figure 8p).

In the adult neurovascular unit, endothelial cells showed the second or third highest expression of SARS-CoV-2 interaction partners after neurons in five out of six datasets; and in one dataset showed a very low expression (Jäkel et al.^70^) (Extended Data Figure 9a-l), whereas in the fetal perivascular niche, endothelial cells displayed the highest expression of SARS-CoV-2 interaction partners (either as stand-alone or tied with neurons, radial glia, and pericytes; and in one dataset the fourth highest after neurons, radial glia, pericytes (La Manno et al.^75^) in five out of six datasets (Extended Data Figure 9m-x). In the pooled analysis, endothelial cells showed a higher expression of SARS-CoV-2 interaction partners as compared to the pool of all perivascular cells in two out of six adult datasets, and in five out of six fetal datasets, with neurons displaying most often the highest expression (Extended Data Figure 9a-x). Together, these data indicate a developmentally established predisposition of brain endothelial cells to respond to a cellular infection with SARS-CoV-2, as reflected by the expression of SARS-CoV-2 interaction partners.

When addressing the different brain endothelial cell clusters according to their arterio-venous specification, we found that in both adult and fetal brain endothelial cell clusters, SARS-CoV-2 interaction partners showed a tendency of higher expression in the veins and capillaries as compared to the arteries (Extended Data Figure 9y-z). In the adult endothelial cells, the cluster endothelial cell 1 EC1 showed the highest expression of SARS-CoV-2 interaction partners, followed by antigen-presenting veins, veins, capillaries, EndoMT, and arteries (Extended Data Figure 9y), whereas SARS-CoV-2 interaction partners showed the highest expression in fetal capillaries > arteries > veins > EC1 (Extended Data Figure 9z).

With CTSB being a SARS-CoV-2 entry associated gene and SARS-CoV-2 interaction partners being intracellular virus-interaction partners post viral cellular entry, we next wondered whether there was a correlation between the top CTSB correlated pathways and the SARS-CoV-2 interaction partners. Notably, the top CTSB-correlated pathways including inflammation, angiogenesis, coagulation, cell-extracellular matrix interaction, vascular metabolism, blood-brain-barrier permeability, and reactive oxygen species (ROS) all displayed a high correlation with SARS-CoV-2 interaction partners in both the adult and fetal brain endothelium (Extended Data Figure 10a-zvi, Extended Data Figure 11a-zvi). Thus, all these pathways that were recently linked to COVID-19 patients with neurological- and vascular symptoms^30,49,81,82^ showed a strong correlation with the intracellular SARS-CoV-2 interaction partners, therefore suggesting a link between CTSB-mediated cellular entry and intracellular signaling post viral entry into the host cell.

### CTSB - but not TMPRSS2 - protein is expressed in the human brain vasculature whereas CTSL– but not TMPRSS2 - protein is expressed in the peripheral vasculature

Finally, to examine the observed mRNA expression patterns of CTSB, CTSL, and TMPRSS2 at the protein level, we took advantage of the publicly available Human Protein Atlas dataset (https://www.proteinatlas.org/).

TMPRSS2 did not show protein expression in the brain, but was expressed in the prostate, jejunum, kidney, and others, again similar to the observed mRNA expression patterns (Figure 5a). Interestingly, and in line with the observations made in the scRNA-seq data, immunohistochemical analysis revealed that TMPRSS2 was neither expressed in the brain endothelium (Figure 5b-d) nor in endothelial cells of peripheral organs (Figure 5e and Extended Data Figure 13a-t), suggesting alternative entry mechanisms employed by the virus to enter vascular endothelial cells in the CNS and the periphery, as mentioned above.

**Figure 5.**
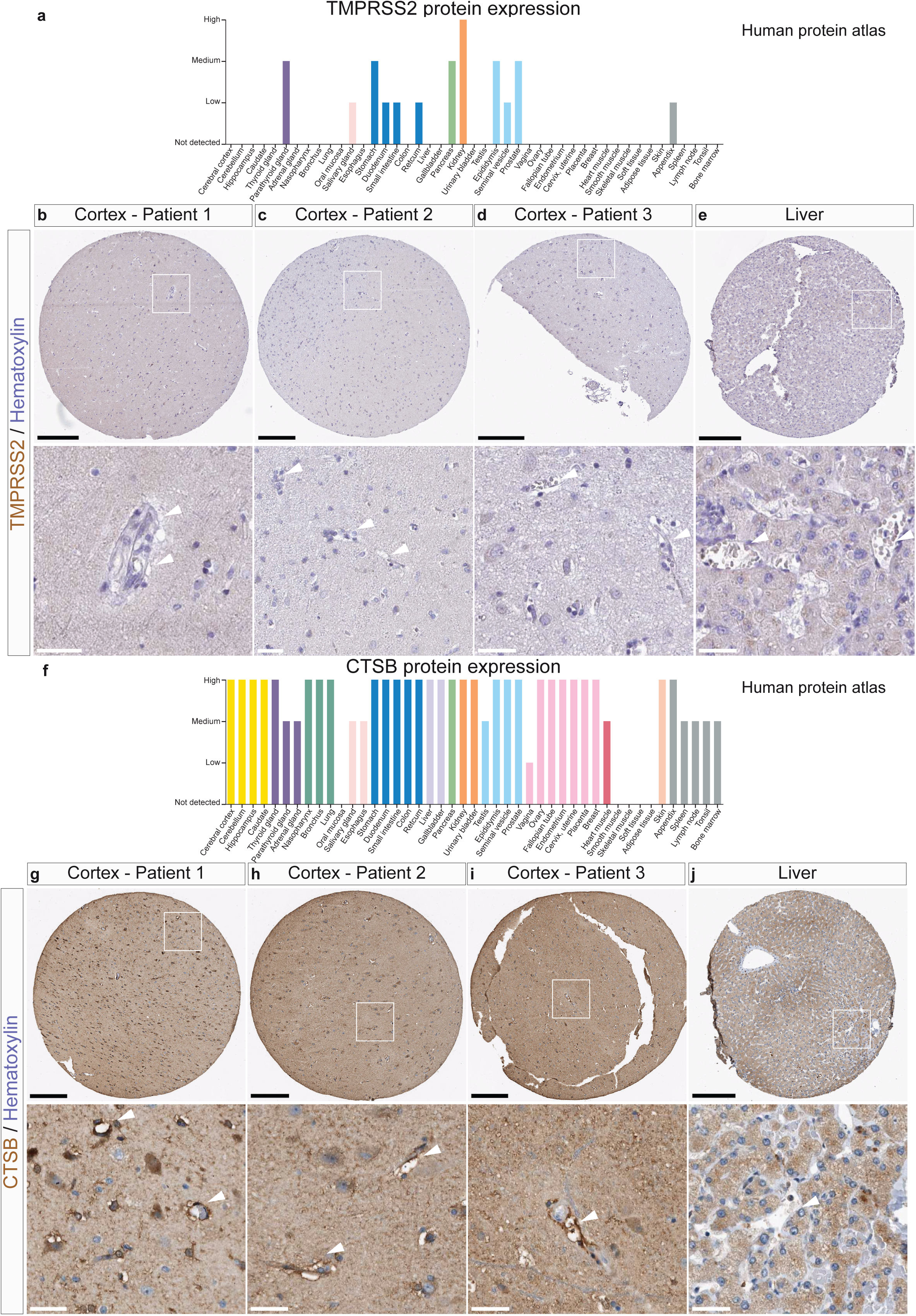
Protein expression of TMPRSS2, CTSB in situ in CNS and non-CNS tissues. **a,** Protein expression scores for TMPRSS2 in human tissues and organs in the human protein atlas revealing no expression in various brain regions and heterogeneous expression scores across various organs. **b**-**e,** Immunohistochemistry (IHC) images for the protein expression of TMPRSS2 in the brain and in the liver indicating that TMPRSS2 is not expressed in blood vessels (arrowheads). **f,** Protein expression scores for CTSB in human tissues and organs in the human protein atlas showing a high expression in various brain regions as well as in most peripheral organs. **g**-**j,** IHC images for the protein expression of CTSB in the brain and in the liver indicating that CTSB is expressed in blood vessels in the brain (arrowheads) but not in the liver (arrowheads). Scale bars: 200 μm for core images, 40 μm for zoomed in images. Arrowheads indicate vascular structures, while arrows indicate positive IHC signals in non-vessel structures.

CTSB showed a high protein expression in various brain regions (cerebral cortex, cerebellum, hippocampus, and caudate), and was highly expressed in numerous other organs known to be affected by systemic SARS-CoV-2, such as the lung, gastrointestinal tract, liver, kidney, and heart, among others (Figure 5f-j and Extended Data Figure 14a-l). These HPA protein expression data are consistent with the observed scRNA sequencing mRNA expression patterns. Immunohistochemical analysis using the validated antibody HPA018156^83–85^, revealed that CTSB was expressed in the brain, displaying high cytoplasmic/membranous expression in brain vascular endothelial cells and in the neuropil, while displaying medium expression in neuronal cells and astrocytes (Figure 5g-i). In the liver, the majority of liver endothelial cells was CTSB negative, with very few blood vessel endothelial cells showing cytoplasmic/membranous CTSB expression (with medium expression levels in hepatocytes and weak expression in bile duct cells, using the same antibody) (Figure 5j). CTSB was also mostly absent in endothelial cells (but positive in non-vascular / non-endothelial structures / cells further validating the antibody) across various peripheral organs including the lung, heart, kidney, colon (Extended Data Figure 14a-l).

In contrast to CTSB, CTSL did not show protein expression in the brain, but was expressed in numerous other organs known to be affected by systemic SARS-CoV-2, such as the prostate, lung, gastrointestinal tract, liver, kidney, jejunum, and colon, among others (Extended Data Figure 12a-e and Extended Data Figure 15a-m). These HPA protein expression data were again consistent with the observed scRNA sequencing mRNA expression patterns. Immunohistochemical analysis using the validated antibody CAB000459^83–85^ revealed that CTSL was expressed in endothelial cells of various organs but not in the brain vasculature, displaying high cytoplasmic/membranous expression in peripheral vascular endothelial cells across various peripheral organs including the liver, placenta, lung, spleen, colon, kidney, pancreas, and heart (Extended Data Figure 15a-m).

To further validate these findings, we performed double immunostainings for an endothelial cell marker (CD31) and CTSB, CTSL, and TMPRSS2 on human temporal lobe specimens from our neurosurgical operating theatre. CTSB showed high expression in CD31^+^ endothelial cells as well as in CD31^-^ cells of the neuropil and intermediate expression in NeuN^+^ neurons, GFAP^+^ astrocytes, IB1^+^ microglia (Figure 6a-zii), in agreement with the observations made in the HPA and the scRNA seq datasets described above. TMPRSS2 and CTSL either were absent in brain endothelial- and perivascular cells or showed very low expression levels (Figure 6a-l), again confirming the scRNA-seq- and human protein atlas data.

**Figure 6.**
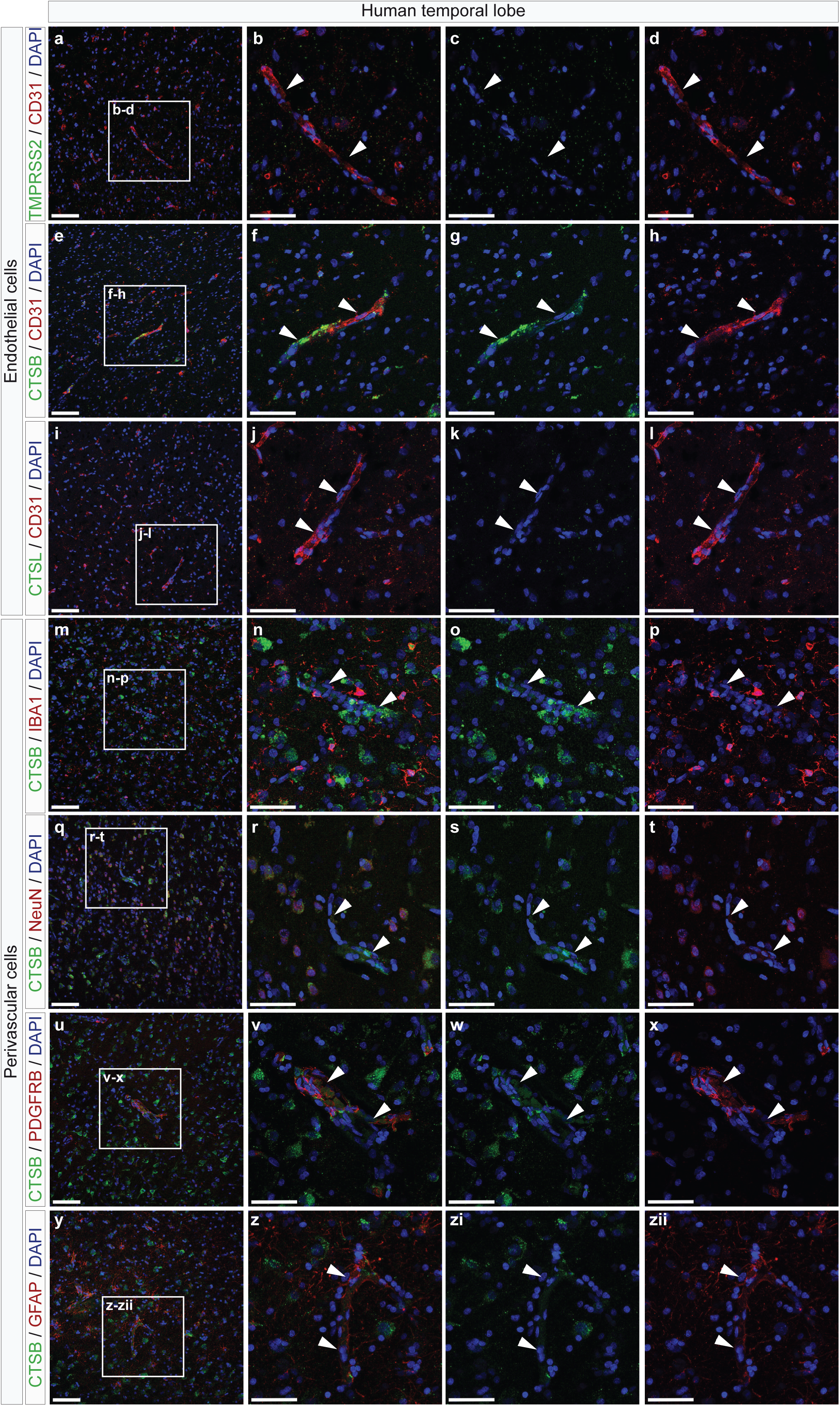
Immunofluorescence staining of TMPRSS2, CTSB and CTSL expression in endothelial cells and perivascular niche of human adult brain vasculature. Tissue sections (40 µm) of human adult brain (derived from temporal lobectomies) were stained for TMPRSS2, CTSB, CTSL (green), the vascular endothelial cell markers CD31 (pan-endothelial marker, red), the microglial marker IBA1 (red), the neuronal marker NeuN (red), the pericyte marker PDGFRB (red), the astrocytic marker GFAP (red), and DAPI nuclear counterstaining (blue). **a-l** Expression of TMPRSS2, CTSB, and CTSL in blood vessel endothelial cells in the adult human brain. **a-d** TMPRSS2 (green) is not expressed in CD31^+^ blood vessel endothelial cells of the human adult brain (**a**). The boxed area in **a** is enlarged in **b-d**, showing absence of TMPRSS2 expression (green) in CD31^+^ blood vessel endothelial cells (red) in the adult human brain. **e-h** CTSB (green) is highly expressed in CD31^+^ blood vessel endothelial cells (red) in the human adult brain (**e**). The boxed area in **e** is enlarged in **f-h**, showing CTSB expression (green) in CD31^+^ blood vessel endothelial cells (red) in the adult human brain. **i-l** CTSL (green) is not expressed in CD31^+^ blood vessel endothelial cells of the human adult brain (**i**). The boxed area in **a** is enlarged in **j-l**, showing absence of CTSL expression (green) in CD31^+^ blood vessel endothelial cells (red) in the adult human brain. **m-zii** Expression of CTSB in perivascular cells of the neurovascular in the adult human brain. **m-p** CTSB (green) is highly expressed in IBA1^+^ microglia (red) in the adult human brain (j). The boxed area in **m** is enlarged in **n-p**, showing CTSB expression (green) in IBA1^+^ microglia (red) in the adult human brain. **m-p** CTSB (green) is highly expressed in IBA1^+^ microglia (red) in the adult human brain (j). The boxed area in **m** is enlarged in **n-p**, showing CTSB expression (green) in IBA1^+^ microglia (red) in the adult human brain. **q-t** CTSB (green) is expressed in NeuN^+^ neurons (red) in the adult human brain (**q**). The boxed area in **q** is enlarged in **r-t**, showing CTSB expression (green) in NeuN^+^ neurons (red) in the adult human brain. **u-x** CTSB (green) is expressed in PDGFRB^+^ pericytes (red) in the adult human brain (**u**). The boxed area in **u** is enlarged in **v-x**, showing CTSB expression (green) in PDGFRB^+^ pericytes (red) in the adult human brain. **y-zii** CTSB (green) is expressed in GFAP^+^ astrocytes (red) in the adult human brain (**y**). The boxed area in **y** is enlarged in **z-zii**, showing CTSB expression (green) in GFAP^+^ astrocytes (red) in the adult human brain. Scale bars: 100 μm in **a,e,i,m,q,u,y** (overviews); and 50 µm in **b**-**d, f-h,j-l,n-p, r-t, v-x, z-zii**, (zooms). Arrowheads indicate vascular structures.

Together, these data indicate that whereas TMPRSS2 is absent in brain- and peripheral vascular endothelial cells, CTSB is expressed in the brain vasculature and CTSL in the peripheral vasculature at both the mRNA and protein levels, suggesting potential alternative mechanism for SARS-CoV-2 cellular entry into the brain and peripheral vasculature.

## DISCUSSION

In this study, analysing multiple scRNA-seq datasets as well as referring to the Human Protein Atlas and performing immunofluorescent stainings, we found that the human brain vasculature expresses a distinct profile of SARS-CoV-2 entry associated genes. Most interestingly, the viral entry-associated protease CTSB but not TMPRSS2 is highly expressed in brain vascular endothelial cells whereas CTSL but not TMPRSS2 is highly expressed in vascular endothelial cells of peripheral organs. These observations suggest a potential mechanism of SARS-CoV-2 viral entry into brain- and peripheral endothelial cells that might underlie the neurovascular- and vascular symptoms observed in some COVID-19 patients (Extended Data Figure 16).

The expression patterns we found can provide insight about the susceptibility of the human vasculature to SARS-CoV-2 infections and could – in a next step – be followed by similar studies on currently still limited clinical samples from COVID-19 patients. Towards this end we examined gene expression of SARS-CoV-2 entry-associated genes and interaction partners in multiple scRNA-seq datasets from different tissues, including the brain, spinal cord and various peripheral tissues. We are aware of the limitations of these analyses, as for instance sparse cell types might be lacking or underrepresented/under-detected due to their low abundance, technical limitations related to isolation protocols, or bioinformatic analyses including technical/computational dropout effects. Thus, the specificity is high (meaning positive results are highly reliable) whereas the sensitivity is limited and thus negative results should be interpreted with care.

SARS-CoV-2 shows systemic effects affecting multiple organ systems including the brain, liver, kidney, heart, and others, and increasing evidence suggests that blood vessel endothelial cells exert crucial roles in the underlying pathogenesis^30,49,81,82^. Whereas cellular entry of SARS-CoV-1 exclusively depends on CTSL and TMPRSS1^50,53^, cellular entry of SARS-CoV-2 can occur either via TMPRSS2 or CTSB/CTSL^53,59^ (Extended Data Figure 16). Thus, the widespread expression of CTSB in the brain (vasculature) and the expression of CTSL in multiple other SARS-CoV-2 affected organs/tissues suggest that CTSB and CTSL might be involved in alternative entry mechanisms and transmission routes that could be responsible for the neurovascular- and vascular phenotypes observed in COVID-19 patients^30,32^. Furthermore, our findings that CTSB is – in addition to the brain endothelium – also highly expressed in microglia (and that CTSL shows high expression in peripheral endothelial cells and macrophages) suggests a potential mechanism involving these two cell types e.g. via an endothelial-to-microglia crosstalk^86,87^, that could explain endotheliitis/endothelialitis in peripheral organs^49,81^ and in the brain^30^. It has been suggested that the SARS-CoV-2 affected endothelial cell is characterized by increased vascular leakage, pro-coagulative and pro-inflammatory states in the lung^82^, and that a combination of those might account for the 30% of COVID-19 patients with severe lung damage in part owed to an overreacting inflammatory response^82^ and multi-organ failure^82^. Interestingly, our endothelial cluster analysis revealed a relative preference of CTSB expression for the venous endothelial side while the correlation analysis showed a high expression of inflammatory-related genes in antigen-presenting (venous) endothelial cells, suggesting a preference of SARS-CoV-2 entry factors and downstream pathways for the venous endothelium. In light of the findings that SARS-CoV-1 caused venous vasculitis in small veins of the brain and lung^88^, and that SARS-CoV-2 causes severe endothelialitis particularly in venous vascular bed^89^ – consistent with our findings – it will be exciting to further investigate the potentially different susceptibility of endothelial cells along the arteriovenous specification.

Moreover, CTSB is known to exert pivotal roles in cancer including brain tumors and brain inflammation/inflammatory brain disease via Interleukin-1β (IL1-β) and tumor necrosis factor-α (TNF-α)^90^, and is capable of crossing the blood-brain-barrier^91^. Thus, we speculate 24 that the co-expression of CTSB in microglia and endothelial cells could account for vascular leakage and opening of the blood-brain-barrier with subsequent increased leukocyte migration across the BBB and infection of the brain vascular endothelium^30,82^ which might explain neurological symptoms such as stroke, epilepsy, necrotizing hemorrhages, and encephalopathy^30,32^. The results from our correlation- and pathway analyses showing that CTSB is tightly linked to genes and pathways involved in viral entry, inflammation, angiogenesis, coagulation, vascular metabolism, and blood-brain-barrier leakage, further supports these speculations. For instance, we observed clusters of antigen (MHC class II)- presenting brain endothelial cells expressing CTSB, possibly explaining cellular crosstalk (for instance with microglia) required for inflammatory responses^82^; as endothelial cells cannot act as antigen-presenting cells on their own. Along these lines, the other cathepsins – in addition to CTSB - showing higher expression in the adult- (CTSA, CTSD) and fetal (CTSC, CTSF) brain endothelium were all linked to inflammatory processes. For instance, CTSB is known to interact with CTSA and CTSD^52,53,92–94^ and to be involved in brain inflammation and other immune functions such as brain inflammation^90^ and immune response^95^. Moreover, the viral receptor/receptor-associated enzyme ST6GAL1 encoding for the human Beta-galactoside alpha-2,6-sialyltransferase 1^96^, is a protein that is involved in the generation of the cell-surface carbohydrate determinants and differentiation antigens HB-6, CDw75, and CD76^97^ and that is found in mouse high endothelial cells of mesenteric lymph node and Peyer’s patches, where its suggested function is to be involved in the B cell homing to Peyer’s patches^98^. Interestingly, our correlation analysis indeed showed a high correlation of CTSB with inflammatory pathways, angiogenesis, and viral-host-interaction, therefore further suggesting a direct link of CTSB with inflammatory, angiogenic, and coagulative responses in the brain endothelium, pathways that were all recently shown to exert pivotal roles in SARS-CoV-2 pathogenesis and COVID-19 patients in peripheral- and CNS organs^30,81,82^. Moreover, 25 inflammatory-, angiogenic-, coagulative-, cell-ECM interaction, vascular/BBB permeability-, metabolism, and oxidative stress pathways all correlated with both the SARS-CoV-2 entry-associated gene CTSB as well with the SARS-CoV-2 intracellular interaction partners, suggesting a link between SARS-CoV-2 entry mechanisms and intracellular signaling. Thus, the role of CTSB and the other SARS-CoV-2 entry-associated genes in inflammatory, angiogenic, and the other aforementioned responses within the brain endothelium deserves further investigation.

For instance, we observed a high correlation of with endothelial glucose- and fatty acid metabolism, known to be key for viral replication and propagation^82,99^, indicating that endothelial cells represent an attractive metabolic target for SARS-CoV-2 infection.

To that regard, it was recently suggested that risk factors for COVID-19 such as old age, obesity, hypertension, and diabetes mellitus are all characterized by pre-existing vascular dysfunction with altered vascular endothelial metabolism^99^. Thus, whether certain endothelial clusters in specific organs and certain pathological conditions display a metabolic signature that is more prone to SARS-CoV-2 infection remains to be explored.

Interestingly, CTSB and CTSL have been shown to substitute TMPRSS2 for viral entry of the Ebola virus, which affects the vasculature in the brain and in peripheral organs^100^, but their role in SARS-CoV-2 remains unknown. Whereas the expression of CTSB in the brain and in multiple peripheral tissues was previously reported^101–105,106,107^, the high expression of CTSB in the brain vasculature endothelium and of CTSL in the peripheral vascular endothelium at both the mRNA and protein levels was not reported to our knowledge, nor was the absence of TMPRSS2. Our own experiments revealed CTSB expression in endothelial cells, neurons, astrocytes, pericytes and microglia within the human brain neurovascular unit and might indeed provide the basis of a potential explanation for some of the putative SARS-CoV-2- mediated effects on the human blood-brain-barrier^30^ as well as some of the observed neurovascular symptoms in COVID-19 patients^30,81^. We thus propose a working model in which SARS-CoV-2 (and to a much lower extent SARS-CoV-1) can infect brain endothelial cells via ACE2 and mainly CTSB and peripheral endothelial cells via ACE2 and mainly CTSL (Extended Data Figure 16). However, regarding the periphery, further validation is needed addressing the CTSL expression patterns within the perivascular niche in various peripheral vascular beds taking into account their tissue-specific properties^108^. We are also well aware of the limitations of our study needing further confirmation using functional assays *in vivo* and *in vitro*.

Taken together, our findings may have important implications for understanding SARS-CoV-2 cellular entry and viral transmissibility in the brain, the brain vasculature, and in peripheral vascular beds. Targeting CTSB (and CTSL) using already approved drugs/known inhibitors (for instance E-64d, ammonium chloride)^53^ could result in inhibiting angiogenesis, vascular metabolism, vascular leakage, and the inflammatory response, resulting in vascular normalization^53,82^. As CTSB is expressed in the brain endothelium, and as SARS-CoV-2 invades the brain (putatively via CTSB) and might affect blood-brain-barrier integrity^30,109^, our discoveries might have important translational implications for both neurovascular- and vascular symptoms observed in COVID-19 patients.

In summary, our work further illustrates the opportunities emerging from integrative analyses of publicly available datasets including scRNA sequencing and the Human Protein Atlas.

## ACKNOWLEDGMENTS

The author(s) disclosed receipt of the following financial support for the research, authorship, and/or publication of this article: T.W. was supported by the OPO Foundation, the Swiss Cancer Research foundation (KFS-3880-02-2016-R, KFS-4758-02-2019-R), the Stiftung zur Krebsbekämpfung, the Kurt und Senta Herrmann Foundation, Forschungskredit of the University of Zurich, the Zurich Cancer League, the Theodor und Ida Herzog Egli Foundation, the Novartis Foundation for Medical-Biological Research, and the HOPE Foundation. T.V was supported by the Natural sciences and engineering research council. I.R was supported by the Canadian Institutes of Health Research.

## AUTHOR CONTRIBUTIONS

T.W. had the idea for the study, designed the experiments, wrote the manuscript, and made the figures. T.W. and M.G. analyzed the data. J.C. and S.T. did the immunofluorescent stainings. T.W. supervised all the research. T.W. wrote the manuscript, M.G. helped editing the manuscript and figures. T.V., P.M., I.R., and G.B. gave critical inputs to the manuscript. All authors read and approved the final manuscript.

## METHODS

### Bioinformatics analysis

Datasets were retrieved from published datasets of multiple human and mouse tissues of the human and mouse cell atlas^63,111^. Adult brain datasets were retrieved from publically available sources including (Jäkel et al.^70^, Welch et al.^71^, Han et al.^63^, Lake et al.^72^, Habib et al.^73^, and Velmeshev et al.^74^ datasets), while fetal brain datasets were obtained from sources including (Bhaduri et al.^110^, Aldinger et al.^112^, Han et al.^63^, Nowakowski et al.^113^, La Manno et al.^75^, Polioudakis et al.^114^). The human cerebral organoids dataset was retrieved from Cakir et al.^77^ dataset. The standard Seurat (version 3.1.5) clustering procedure was followed, raw expression values were normalized and log transformed. The original author annotation and cell clustering was retained based on the original studies when available. For integration of the fetal brain endothelial cell datasets, Harmony/Seurat-wrappers package (SeuratWrappers version 0.1.0) was used to correct batch effects in the principal component space and the corrected principal components were used for computing nearest-neighbor graphs^115^. To annotate the endothelial cell clusters of the adult and fetal datasets, the cell identity classification and label transfer was done using the standard Seurat workflow using the Vanlandewijck et al. as the reference dataset^78^. Illustration of the results was generated using Seurat (v.3.1.5). For correlation analysis with CTSB, we performed Spearman’s correlation using the R Stats package (version 4.0.2) on the endothelial cells of the Velmeshev et al^74^ adult brain dataset and the integrated fetal brain datasets (Aldinger et al.^112^, Han et al.^63^, La Manno et al.^75^, Polioudakis et al.^114^, Nowakowski et al.^113^, and Bhaduri et al.^110^ datasets). The correlation coefficients for all genes are included as Supplementary table 1 and 2. The top 50 genes in each dataset were characterized based on GeneCards information and gene ontology classes from the Gene Ontology database.

Pathway analysis was performed using the Gene Set Enrichment Analysis (GSEA) software from the Broad Institute (software.broadinstitute.org/GSEA)(version 4.0.1). A permutation-based p-value is computed and corrected for multiple testing to produce a permutation based Benjamini – Hochberg correction false-discovery rate (FDR) q-value that ranges from 0 (highly significant) to 1 (not significant). The resulting pathways are ranked using NES, and FDR q-value, p-values are reported in the GSEA output reports. Human_GOBP_AllPathways_no_GO_iea_May_01_2020_symbol from [http://baderlab.org/GeneSets] was used to identify enriched pathways in GSEA analysis. Highly related pathways were grouped into a theme and labeled by AutoAnnotate (version 1.3) in Cytoscape (Version 3.7.0) and EnrichmentMap (version 3.3) was used to plot the pathways.

### Human adult brain tissue

Freshly resected tissue samples of the neocortical part of temporal lobes of anonymous, pharmacoresistant epilepsy patients were obtained from the Division of Neurosurgery, Toronto Western Hospital, University Health Network, University of Toronto. Tissues were fixed at 4°C in 4% paraformaldehyde (PFA) for 12 hours, and placed into 30% sucrose in PBS solution overnight. The tissues were then embedded in Optimum Cutting Temperature compound (Tissue-Tek O.C.T. Compound, #4583) and stored in a −80°C freezer at Toronto Western Hospital, University Health Network, Toronto. All the procedures performed with the use of samples obtained from patients were approved by the Institutional Research Ethics Review Board of the University Health Network (Approval Number: 13-6009).

### Immunofluorescence staining of TMPRSS2, CTSB, and CTSL in endothelial- and perivascular cells of the human adult brain vasculature

Fixed, Cryo-embedded, normal human temporal lobe of adult brain slices were cut in 40-µm thick sections, using a cryotome (Leica Cryostat 1720 Digital Cryotome), and submitted for single and double staining with the following antibodies (Abs): pig pAb anti-CD31 (1:400; Synaptic Systems #351004), rabbit mAb anti-CTSB (1:2000, Cellsignal #31718T), rabbit pAb anti-CTSL (1:1000, Ptglab, #10938-1-AP), rabbit mAb anti-TMPRSS2 (1:1000, Abcam, #ab109131), goat pAB anti-Ib1 (1:400, Novusbio, #NB100-1028SS), mouse mAB anti-GFAP (1:400, Sigma-Aldrich, #G3893), mouse mAB anti-NeuN (1:200, EMD Millipore, #MAB377), goat pAB anti-PDGFRβ (1:100, R&D Systems, #AF385). Briefly, the sections were: 1) permeabilized with 0.3% Triton X-100 in PBS for 30 min at room temperature (RT); 2) incubated overnight at 4°C with primary Abs, CTSB/CD31, CTSL/CD31, TMPRSS2/CD31, CTSB/GFAP, CTSB/NeuN, CTSB/Ib1 and CTSB/PDGFRβ 3) incubated with the appropriate secondary Abs, donkey anti-mouse Alexa Fluor 488 (1:1000, Thermo Fisher Scientific, #A-21202), donkey anti-Guinea pig Alexa Fluor 488 (1:1000, Jackson Immunoresearch Labs, #706-545-148), donkey anti-rabbit Alexa Fluor 555 (1:1000, Thermo Fisher Scientific, #A-31572), donkey anti-goat Alexa Fluor 488 (1:1000, Thermo Fisher Scientific, #A-11055) for 90 min at RT; to quench the autofluorescence signal, tissue sections were treated with 0.1% Sudan Black B (Thermo Fisher Scientific, Cat: J62268) in 70% ethanol for 15 minutes at room temperature. 4) counterstained with the 4, 6-diamidino-2-phenylindole (DAPI) (diluted 1:20’000; BioLegend, #422801). Finally, the sections on glass slides (Fisherbrand, Superfrost Plus Microscope Slides, Fisher Scientific) were coverslipped with VWR micro cover glass (VWR International). Negative controls were prepared by omitting the primary antibodies and mismatching the secondary antibodies. Sections were examined under Zeiss laser scanning confocal microscope (LSM 880). Laser scanning confocal images were taken through the z-axis of the section, with 20x and 40x lenses. Z-stacks of optical planes (maximum intensity projections) and single optical planes were recorded and analysed by Zeiss Zen software.

## Extended DATA FIGURE LEGENDS

**Extended Data Figure 1.**
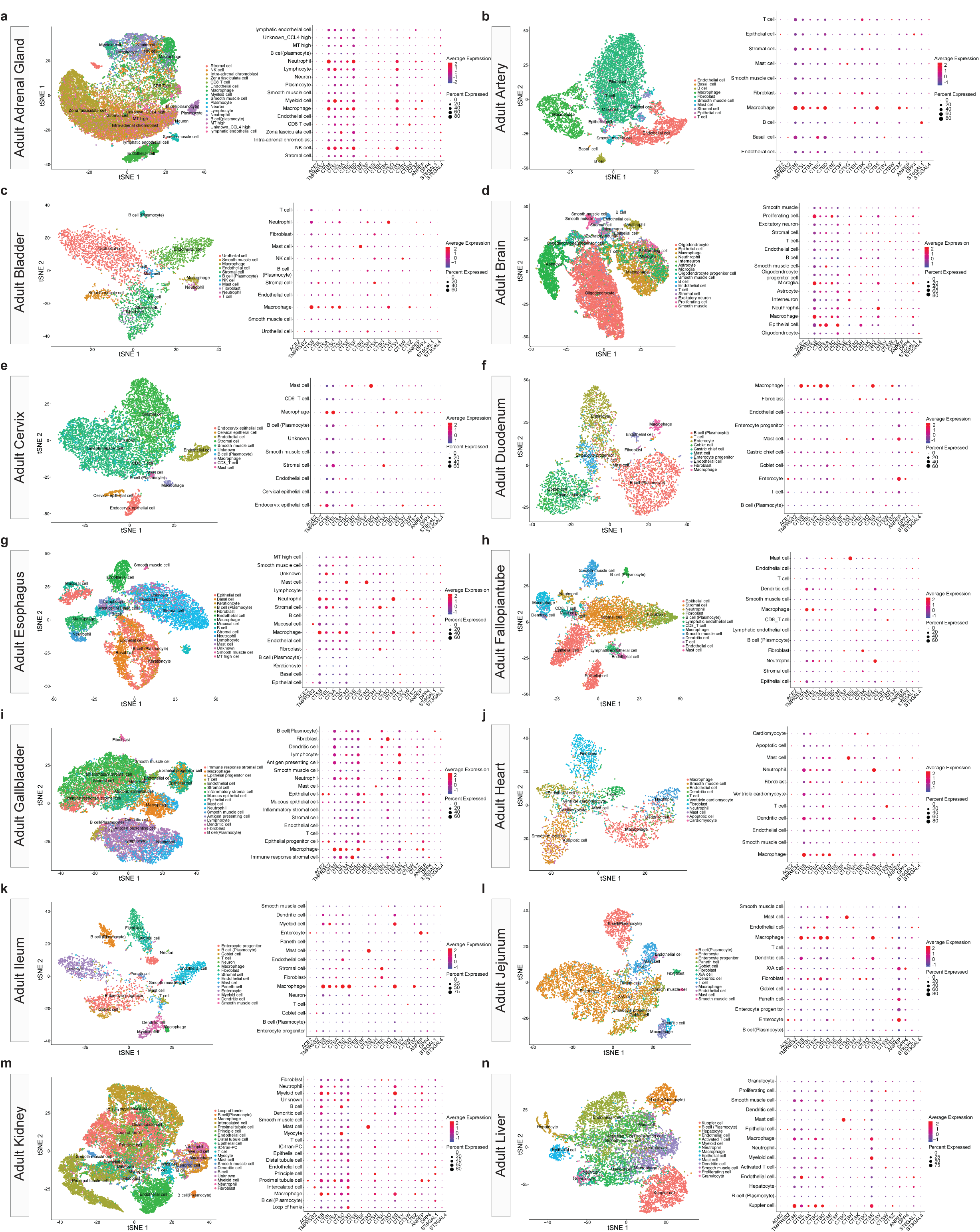
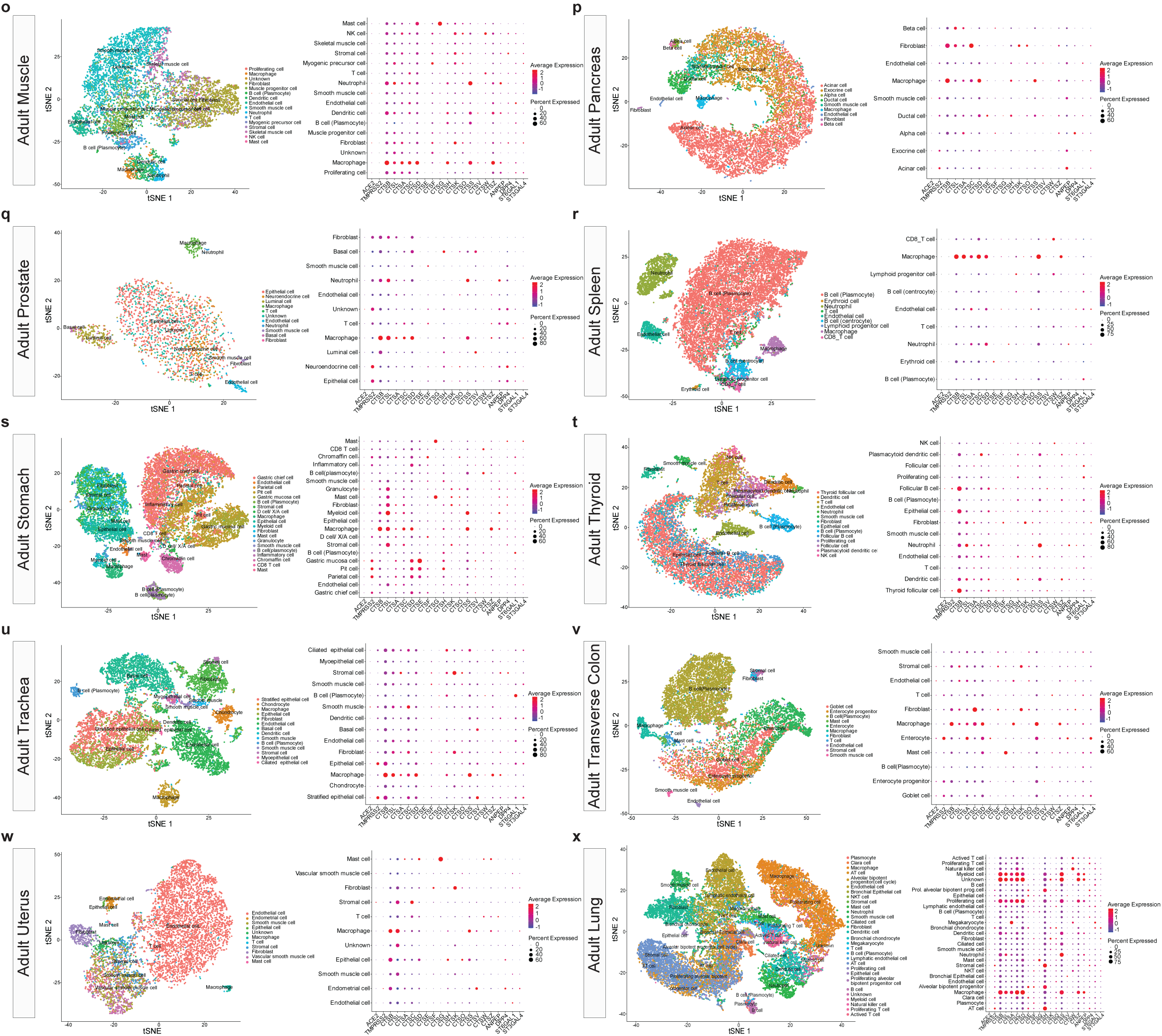
Expression of SARS-Cov-2 entry associated genes in human adult tissues. **a-x,** t-SNE plot of single cells of individual adult tissues from the human cell landscape scRNA-seq dataset^63^. In the *t*-SNE plots for the individual tissues, each cell type is labelled by a different color and every cell type represents an individual cluster; 26 adult tissues were analyzed, namely adrenal gland (**a**), artery (**b**), bladder (**c**), brain (**d**), cervix (**e**), duodenum (**f**), esophagus (**g**), fallopian tube.(**h**), gallbladder (**i**), heart (**j**), ileum (**k**), jejunum (**l**), Kidney (**m**), liver (**n**), muscle (**o**), pancreas (**q**), prostate (**r**), spleen (**s**), stomach (**t**), thyroid (**u**), trachea (**v**), transverse colon (**w**), uterus (**x**) and lung (**z**). Dotplots show RNA expression of SARS-CoV-2 entry receptor, entry-associated proteases, other cathepsins and other viral receptors/receptor-associated enzymes in the different cell types of each of the adult tissues analysed. The dot size represents the proportion of cells within the respective tissue type expressing the gene and the dot color represents the average gene expression level within the particular tissue type.

**Extended Data Figure 2.**
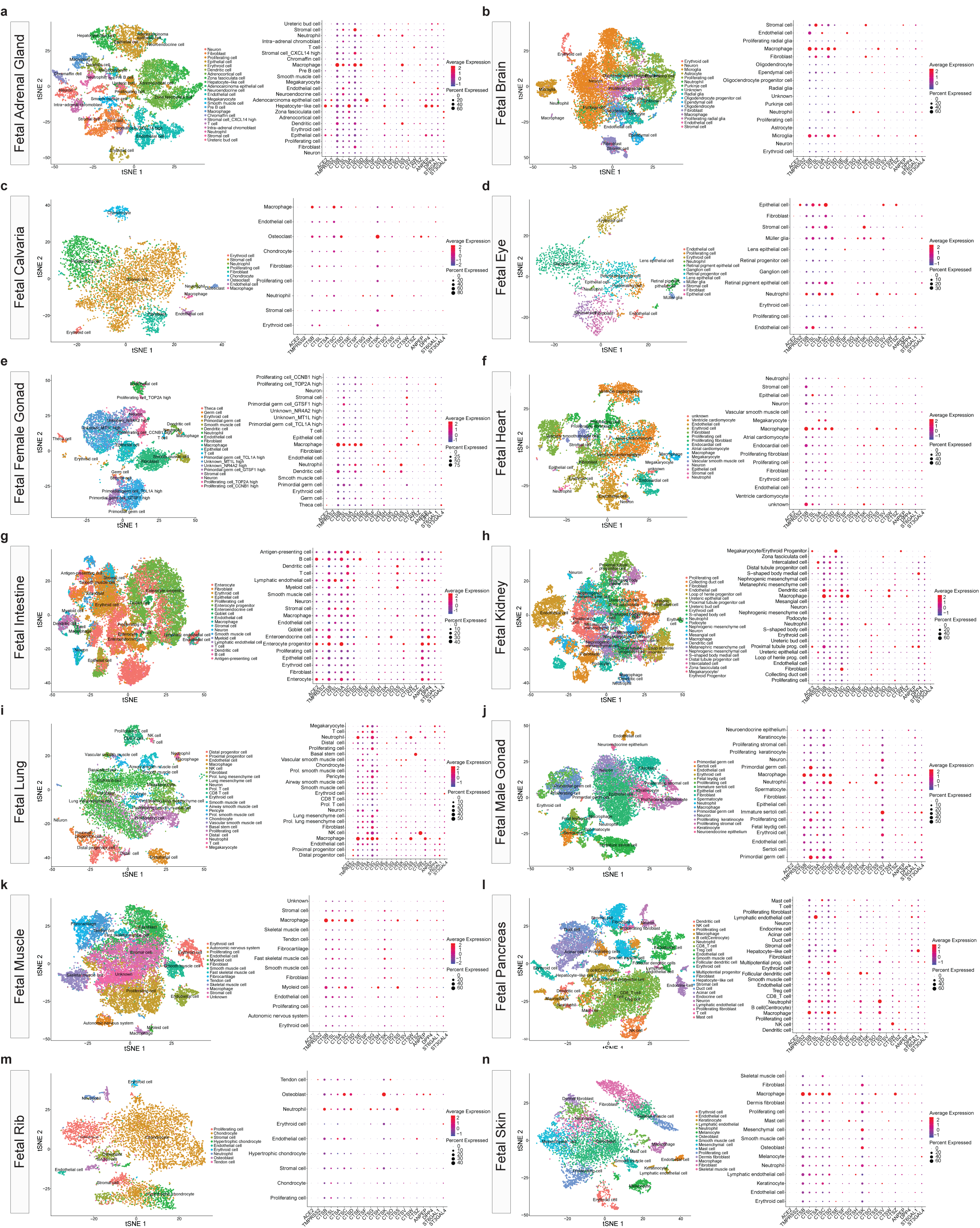
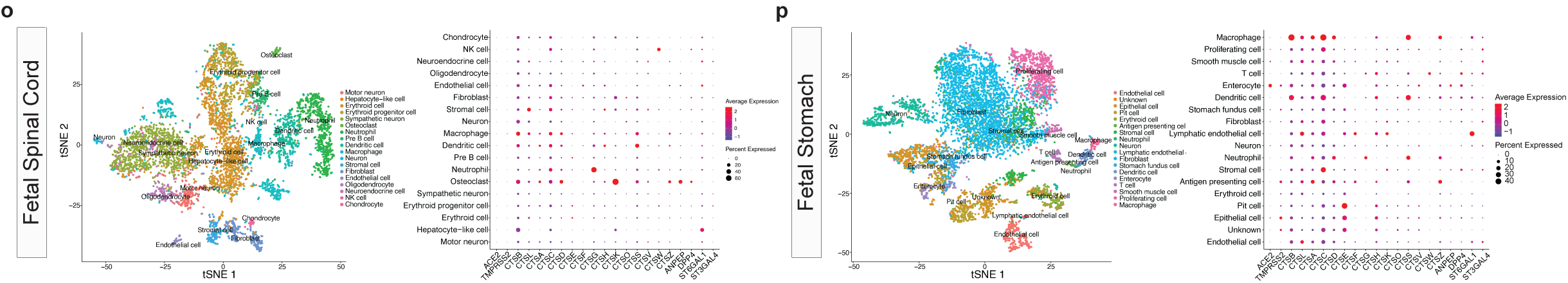
Expression of SARS-Cov-2 entry associated genes in human fetal tissues. **a-p**, t-SNE plot of single cells of individual fetal tissues from the human cell landscape scRNA-seq dataset^63^. In the *t*-SNE plots for the individual tissues, each cell type is labelled by a different color and every cell type represents an individual cluster; 16 fetal tissues were analyzed, namely adrenal gland (**a**), brain (**b**), calvaria (**c**), eye (**d**), female gonad (**e**), heart (**f**), intestine (**g**), kidney (**h**), lung (**i**), male gonad (**j**), muscle (**k**), pancreas (**l**), rib (**m**), skin (**n**), spinal cord (**o**) and stomach (**p**). Dotplots show RNA expression of SARS-CoV-2 entry receptor, entry-associated proteases, other cathepsins and other viral receptors/receptor-associated enzymes in the different cell types of each of the fetal tissues analysed. The dot size represents the proportion of cells within the respective tissue type expressing the gene and the dot color represents the average gene expression level within the particular tissue type.

**Extended Data Figure 3.**
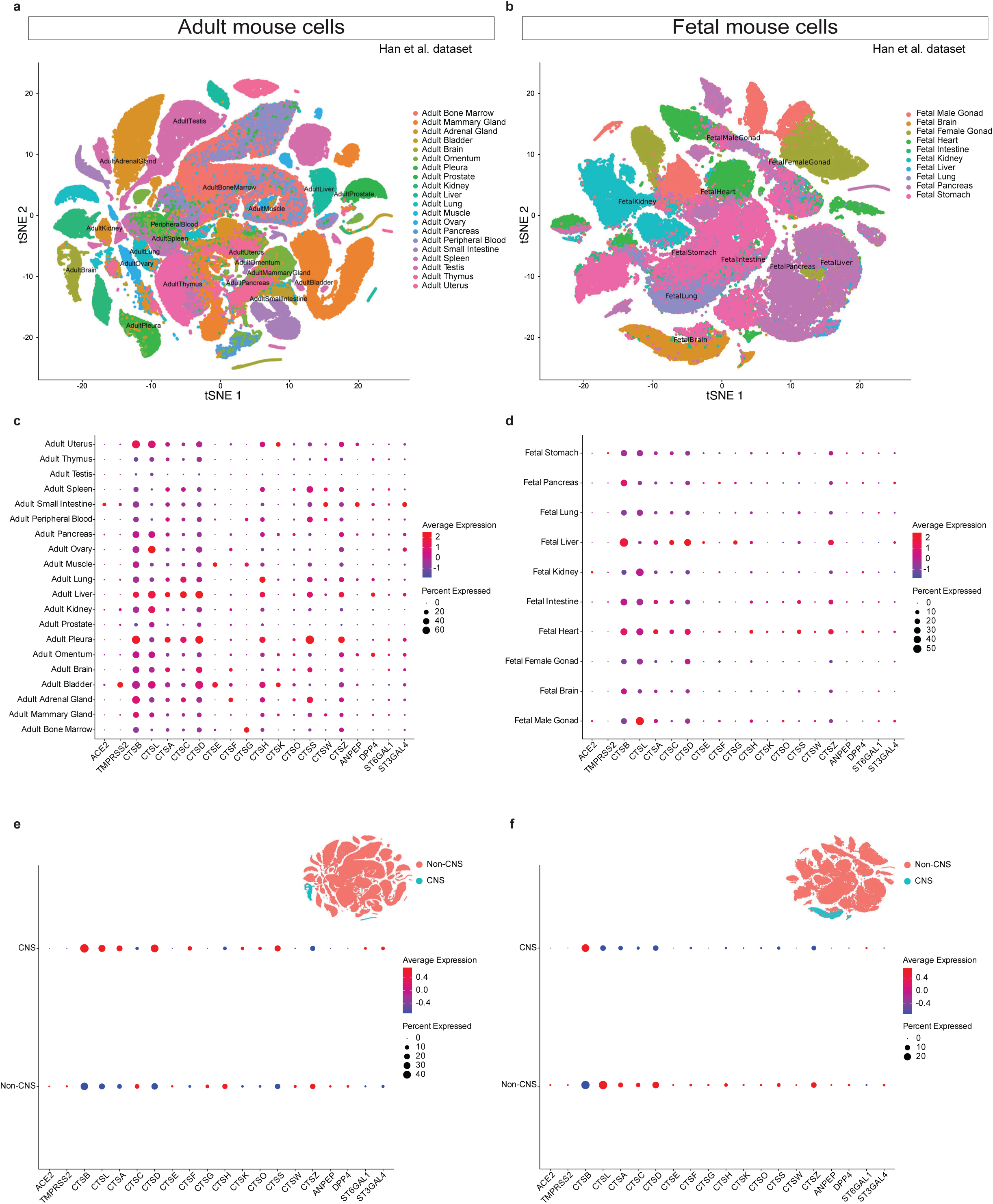
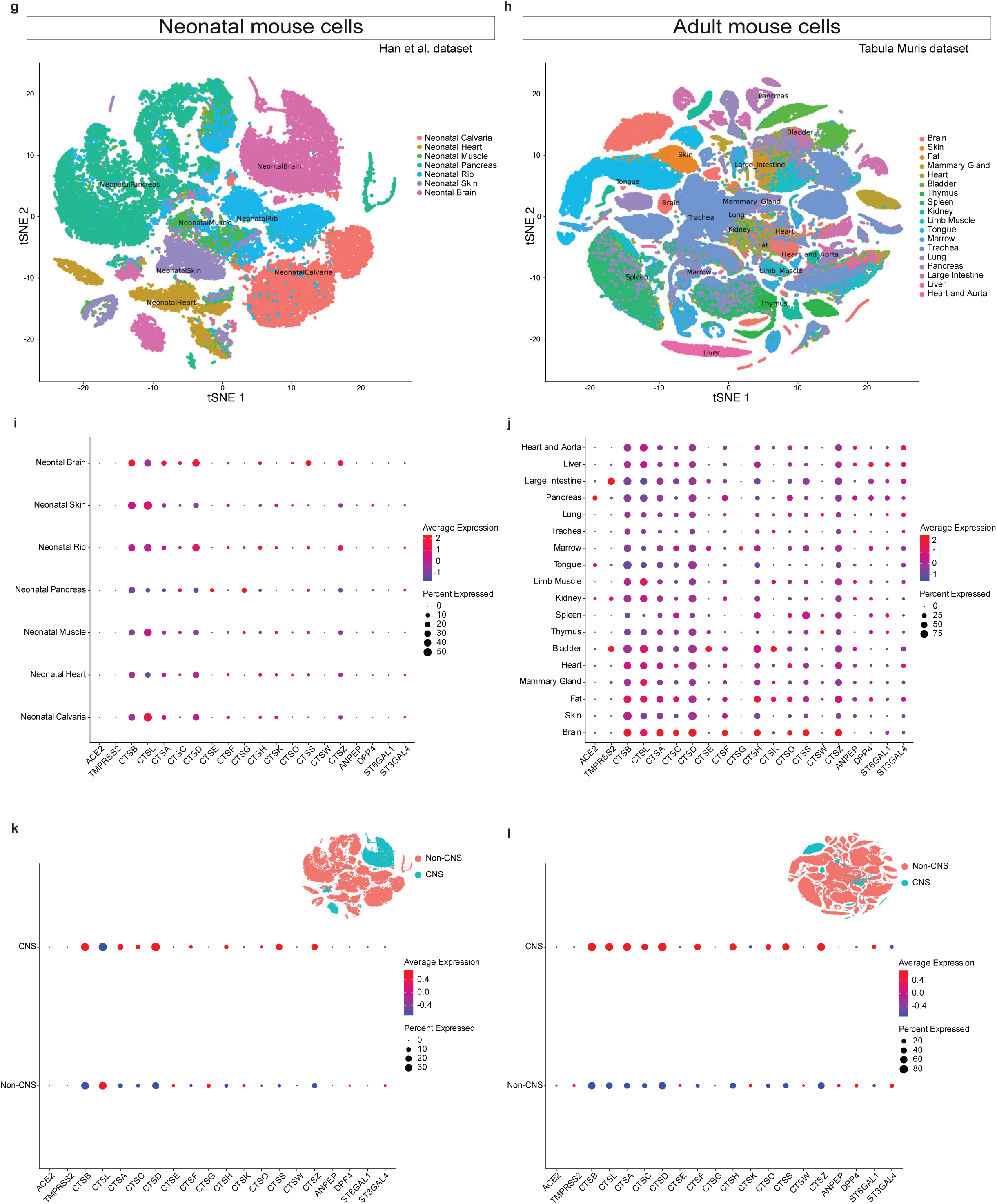
Expression of SARS-Cov-2 entry associated genes in mouse adult and fetal CNS and non-CNS tissues. **a,b,g,** t-SNE plot of 156’831, 81’240 and 59’637 single cells of adult (**a**), fetal (**b**), and neonatal (**g**) tissues respectively from the mouse cell atlas^111^. **h,** t-SNE plot of 123’878 single cells of adult (**h**) from the Tabula Muris dataset^116^. In the *t*-SNE plot, tissues are labelled by different colors, and every tissue represents an individual cluster; 20 adult, 10 fetal, and 7 neonatal tissues were analyzed from the mouse cell atlas dataset, while 18 adult mouse tissues were analyzed from the Tabula Muris dataset. **c,d,i,j,** RNA expression of the SARS-CoV-2 entry receptor ACE2, entry-associated proteases: (TMPRSS2, CTSB, CTSL), other cathepsins (CTSA, CTSC, CTSD, CTSF, CTSG, CTSH, CTSK, CTSO, CTSS, CTSV, CTSW, CTSZ), and other viral receptors/receptor-associated enzymes (ANPEP, DPP4, ST6GAL1, ST3GAL4) in adult, fetal, and neonatal mouse cells from the mouse cell atlas and Tabula Muris datasets across different tissues. **e,f**,**k,l,** RNA expression of SARS-CoV-2 entry receptor, entry-associated proteases, other cathepsins, and other viral receptors/receptor-associated enzymes in CNS vs non-CNS cells of adult, fetal, and neonatal mouse tissues. The datasets were obtained from existing publically available sources and tissue clustering and nomenclature were retained based on the author annotation^111,116^. Expression values were normalized, log transformed and scaled, for results in the dot plots, the dot size represents the proportion of cells within the respective tissue type expressing the gene and the dot color represents the average gene expression level within the particular tissue type.

**Extended Data Figure 4.**
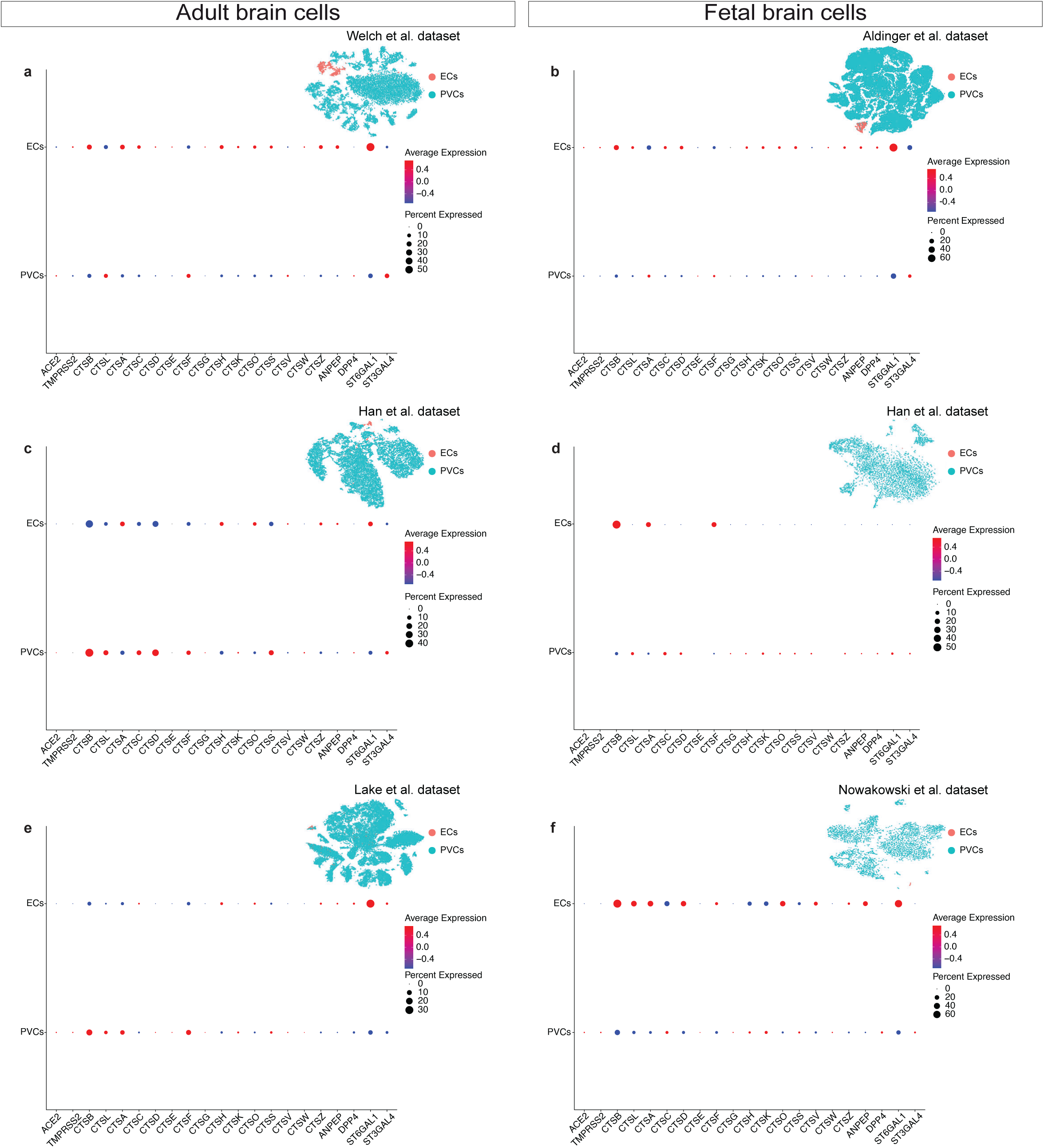
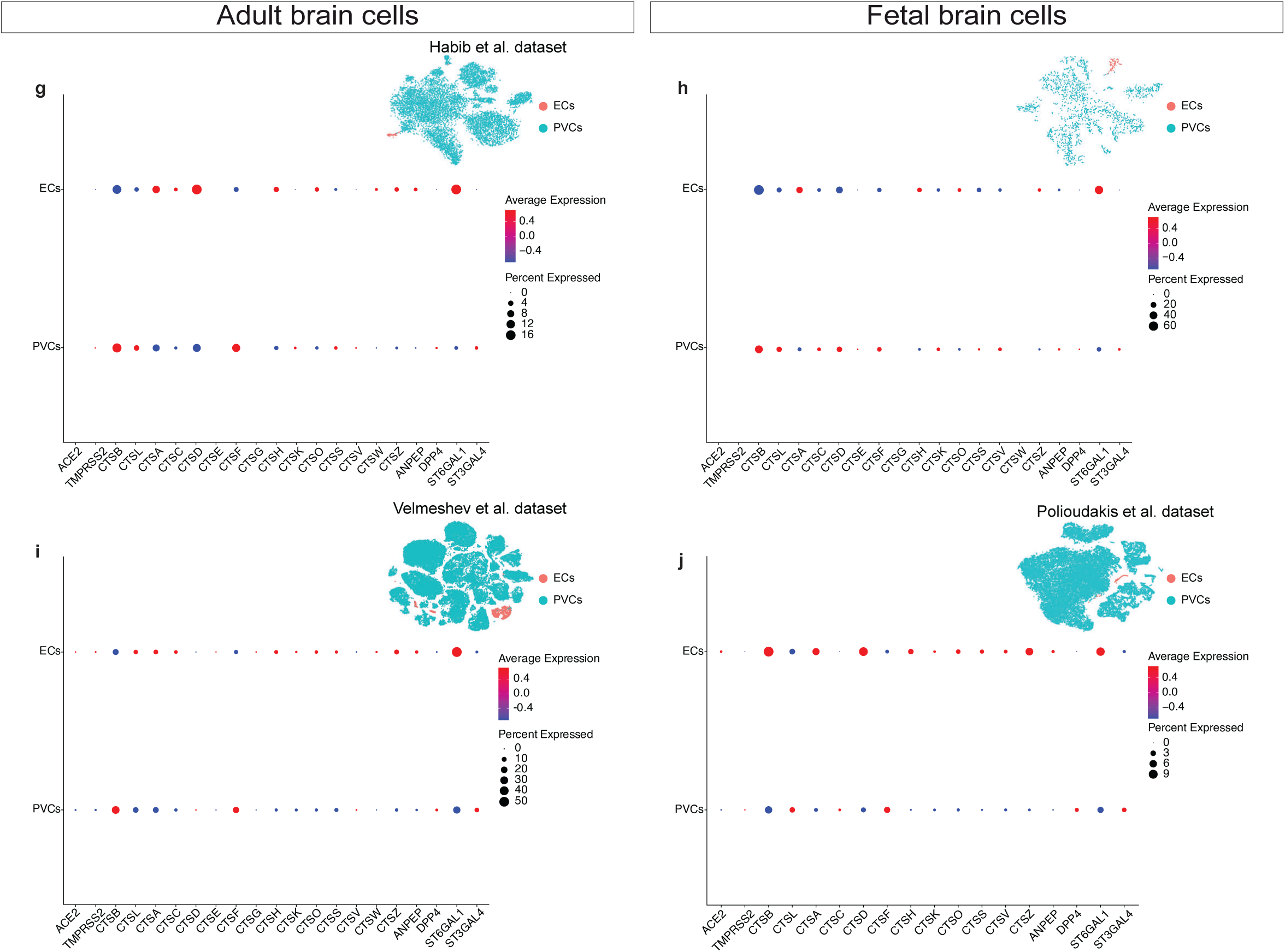
Expression of SARS-Cov-2 entry associated genes in human adult and fetal brain cells comparing endothelial cells to perivascular cells. **a-j,** RNA expression of SARS-CoV-2 entry receptor, entry-associated proteases, other cathepsins, and other viral receptors/receptor-associated enzymes in endothelial cells vs perivascular cells of human adult^63,71–74^ (**a,c,e,g,i**) and fetal^63,75,112–114^ (**b,d,f,h,j**) brain cells from various datasets indicated.

**Extended Data Figure 5.**
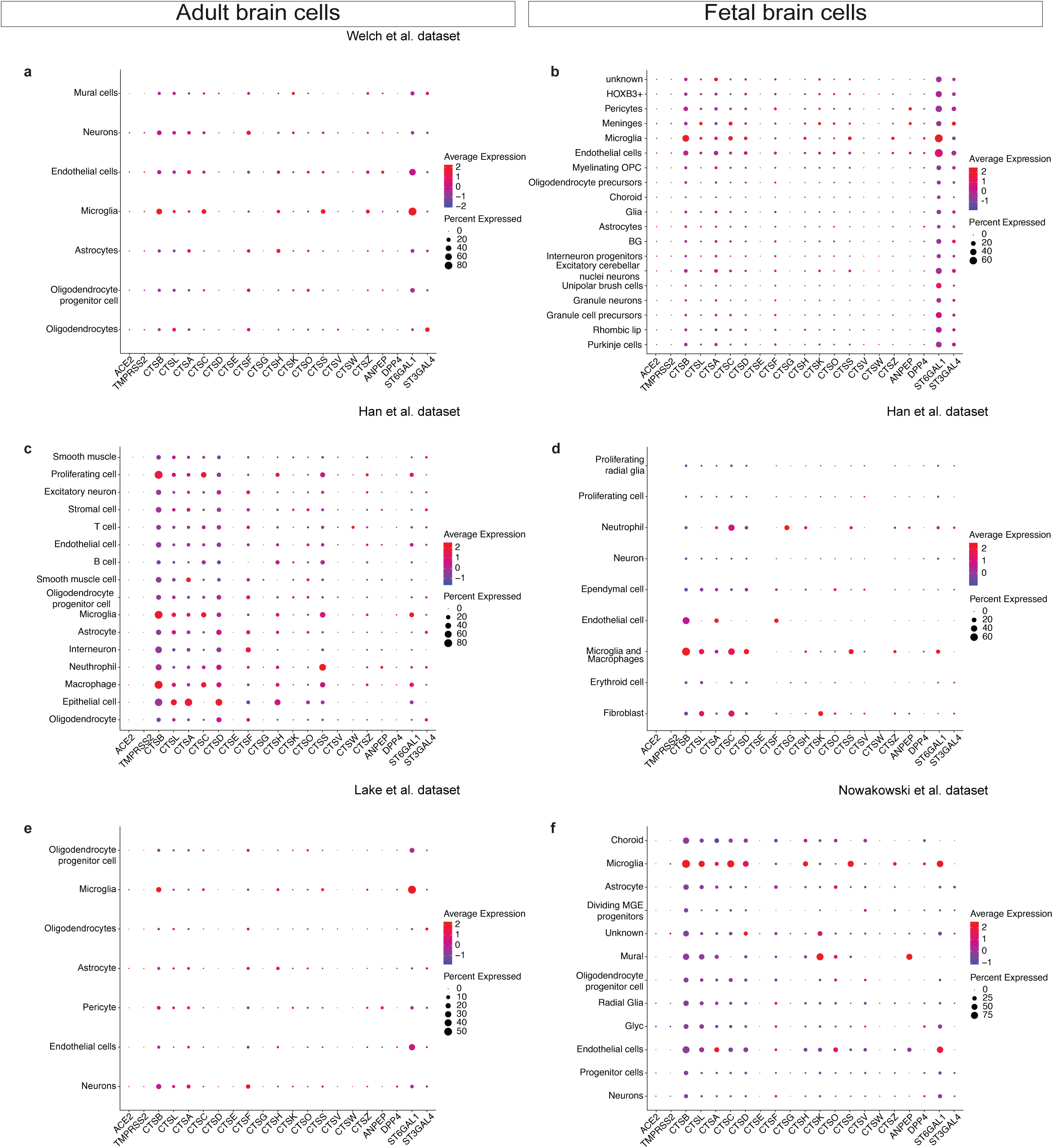
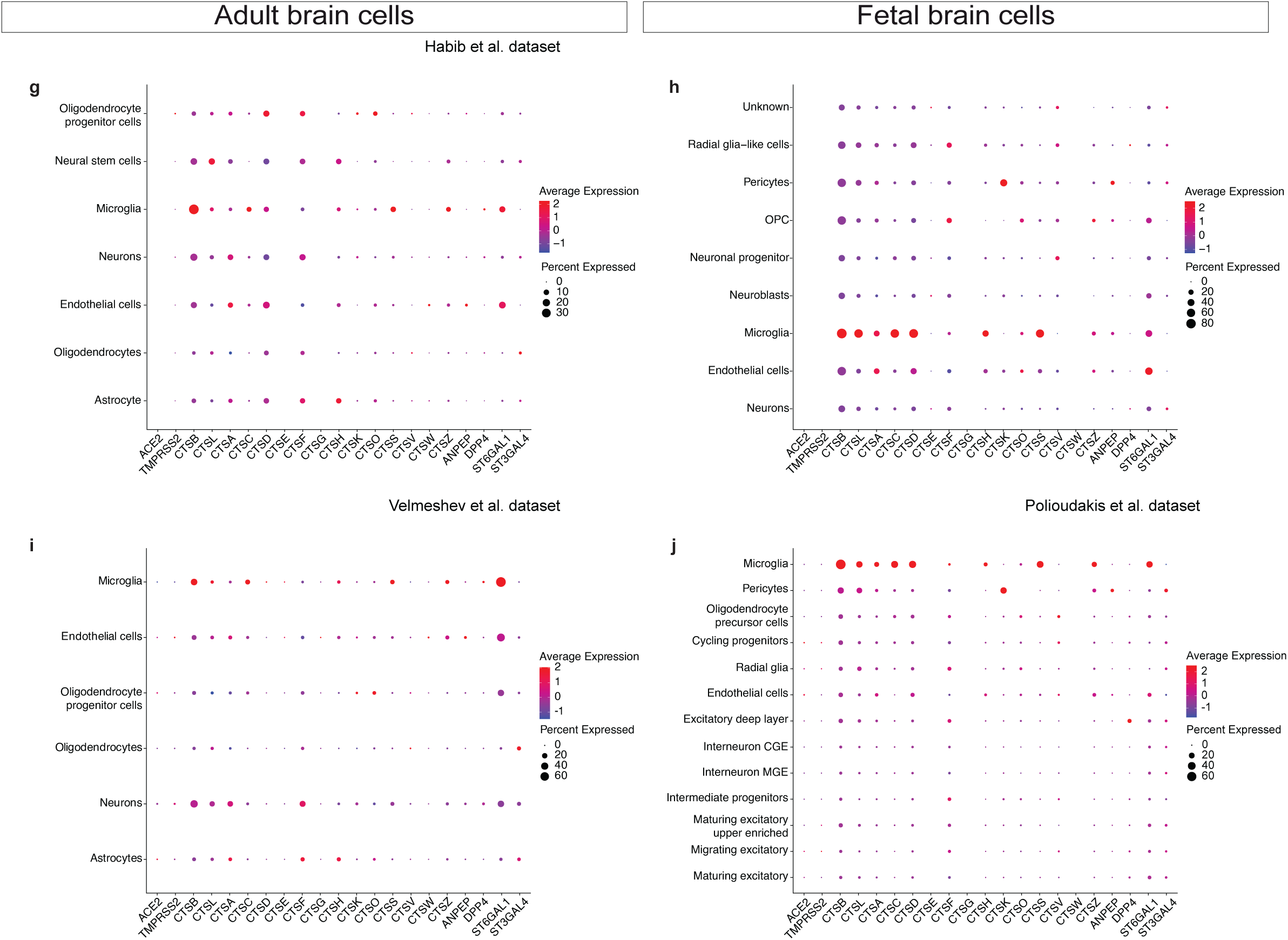
Expression of SARS-Cov-2 entry associated genes in human adult and fetal brain cells. **a-j,** RNA expression of SARS-CoV-2 entry receptor ACE2, entry-associated proteases, other cathepsins, and other viral receptors/receptor-associated enzymes in adult^63,71–74^ (**a,c,e,g,i**) and fetal^63,75,112–114^ (**b,d,f,h,j**) brain cells from various scRNA-seq datasets indicated.

**Extended Data Figure 6.**
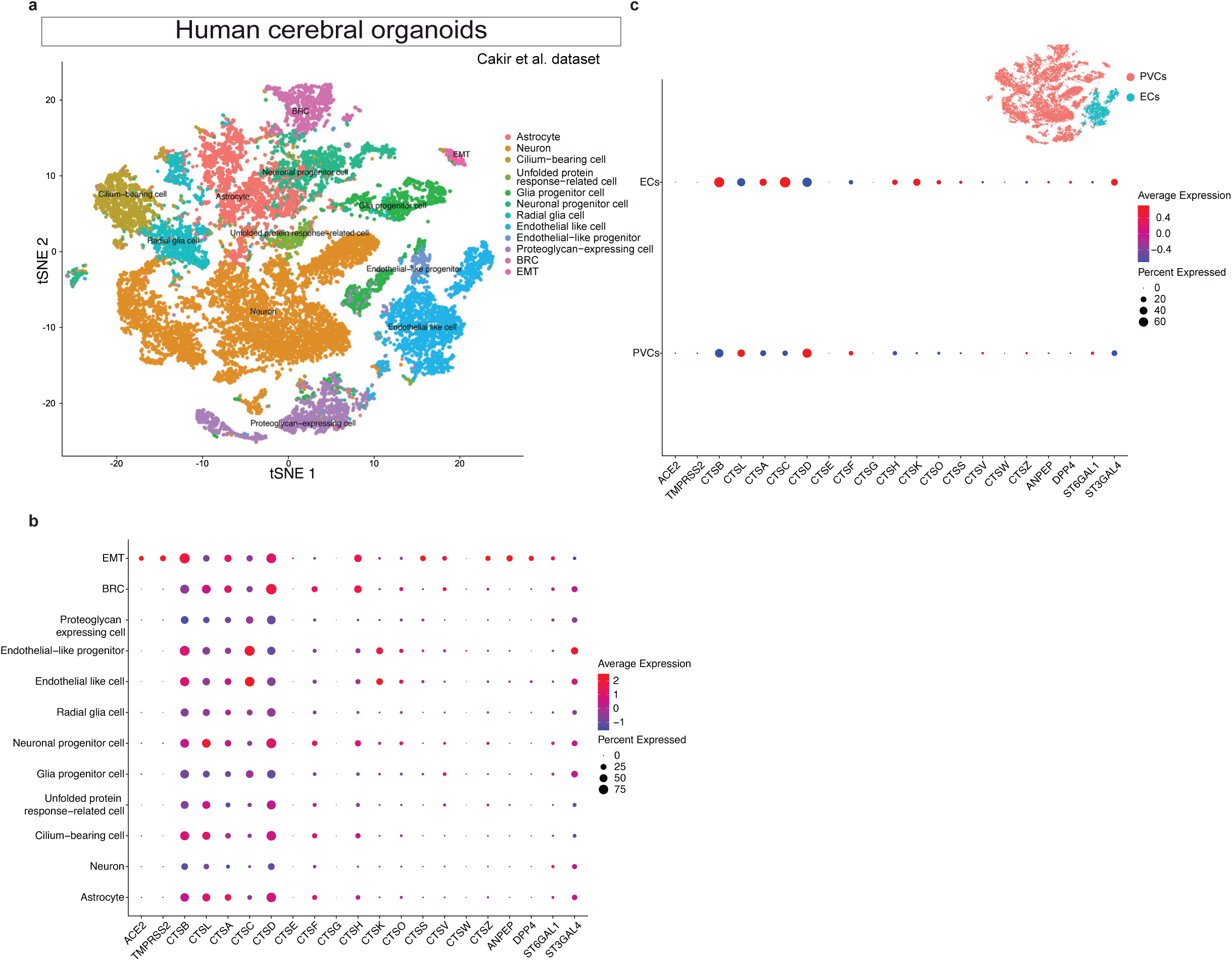
Expression of SARS-Cov-2 entry associated genes in human brain organoids. **a,** t-SNE plot of 20’026 single cells of cerebral organoids from the Cakir et al.^77^. dataset. In the *t*-SNE plot cell types are labelled by different colors and every cell type represents an individual cluster. **b,** RNA expression of the SARS-CoV-2 entry receptor ACE2, entry-associated proteases, other cathepsins, and other viral receptors/receptor-associated enzymes in cerebral organoids cells from Cakir et al. scRNA-seq dataset^77^. **c,** Dotplot comparing expression of SARS-CoV-2 entry receptor, entry-associated proteases, other cathepsins, and other viral receptors/receptor-associated enzymes in endothelial cells vs perivascular cells of the cerebral organoids.

**Extended Data Figure 7.**
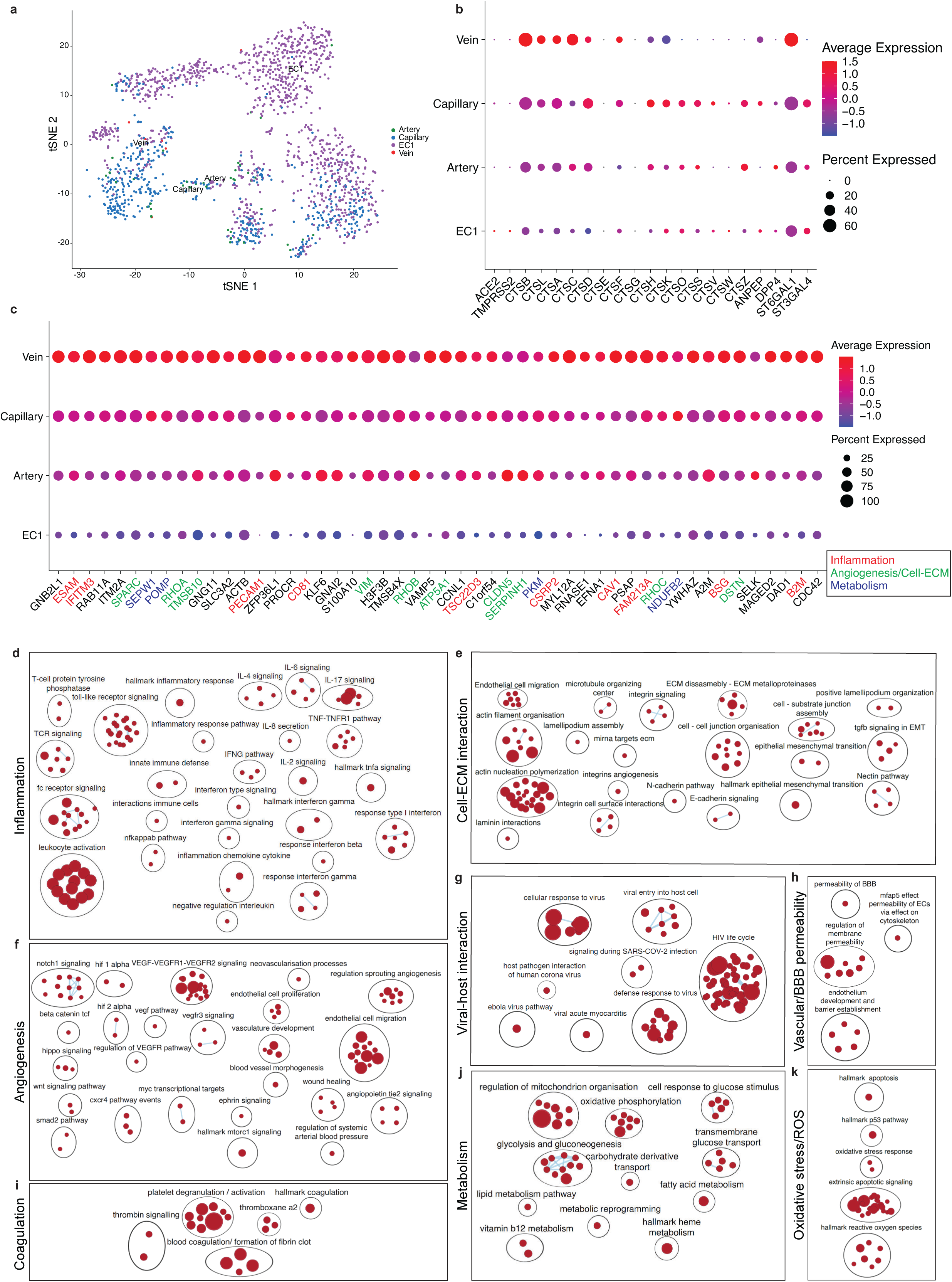
Expression of SARS-Cov-2 entry associated genes in human fetal brain endothelial cells. **a**, t-SNE plot of 1’672 endothelial cells of fetal brain tissue computationally extracted from the Bhaduri et al.^110^, Aldinger et al.^112^, Han et al.^63^, Nowakowski et al.^113^, La Manno et al.^75^, and Polioudakis et al.^114^ datasets. The endothelial cells from the different datasets were integrated using Harmony/Seurat-wrappers^115^. In the *t*-SNE plot, endothelial cell sub-types are labelled by different colors and every endothelial cell subtype represents an individual cluster. **b**, RNA expression of SARS-CoV-2 entry receptor ACE2, entry-associated proteases, other cathepsins and other viral receptors/receptor-associated enzymes in adult brain endothelial cell sub-types from the six studied datasets. **c**, Brain endothelial expression of the top 50 genes correlated with CTSB expression based on Spearman’s correlation analysis performed on endothelial cells computationally extracted from the six aforementioned datasets. The colored gene names represent genes that are immune-associated/inflammatory response related (red), angiogenesis/cell-ECM interaction related (green), and metabolism related (blue). For gene expression results in the dot plots, the dot size represents the proportion of cells within the respective cell type expressing the gene and the color represents the average gene expression level within the particular endothelial cell sub-type. (**d-k**), Enrichment map visualizing the GSEA pathway analysis results performed on the ranked list of the CTSB correlating genes, the top significantly enriched pathways included inflammation (**d**), angiogenesis (**f**), coagulation (**i**), cell-extracellular matrix interaction (**e**), viral-host interaction (**g**), vascular metabolism (**j**), blood-brain-barrier permeability (**h**), and reactive oxygen species (ROS) (**k**) in fetal brain endothelial cells.

**Extended Data Figure 8.**
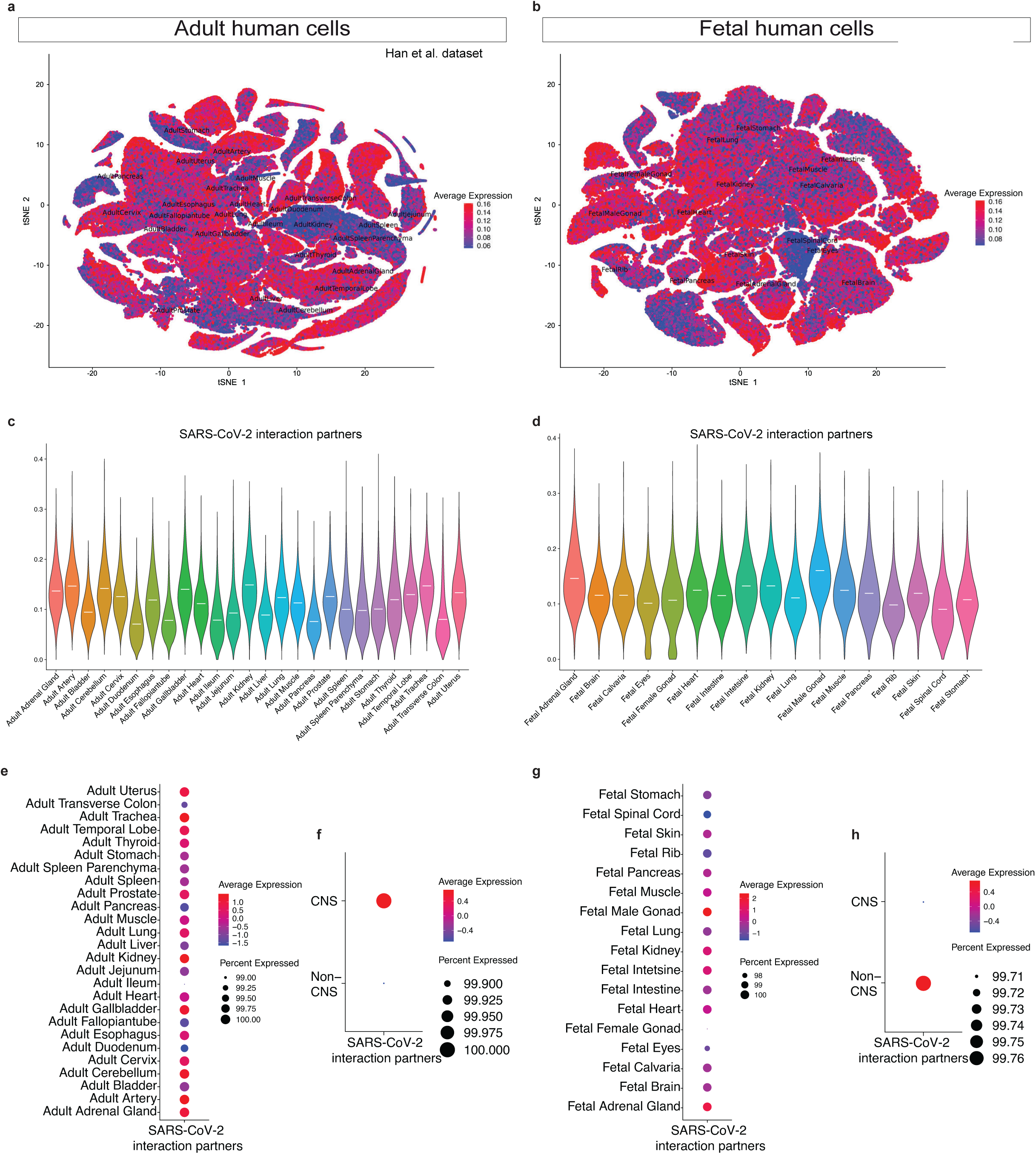
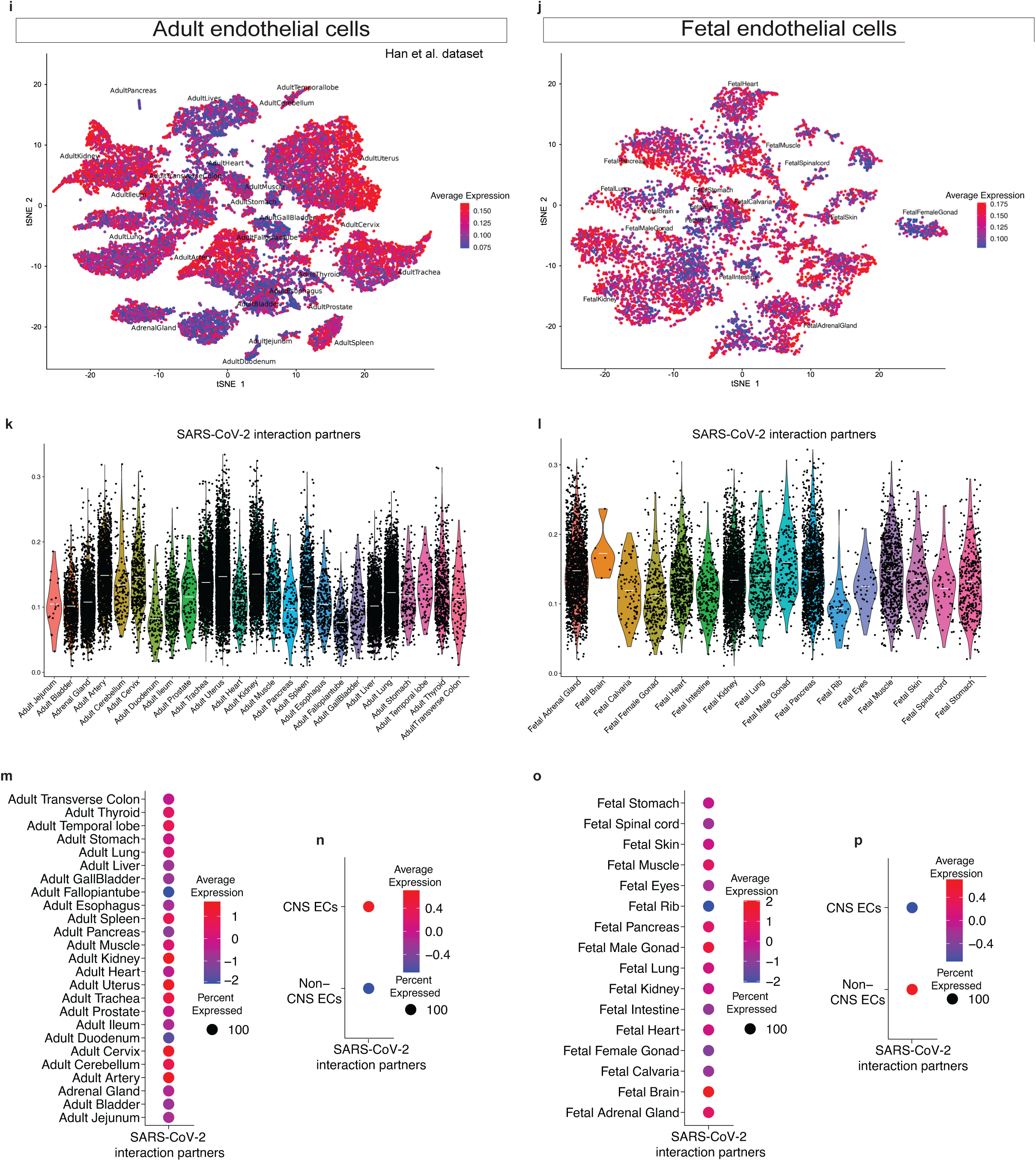
Expression of SARS-Cov-2 interaction partners in human adult and fetal CNS and non-CNS tissues. **a,b,** t-SNE plot showing the expression of SARS-CoV-2 interaction partners in all cells of adult (**a**) and fetal cells (**b**) tissues from the human cell landscape scRNA-seq dataset^63^. In the *t*-SNE plot, cells are colored according to their degree of expression of SARS-CoV-2 interaction partners (red is the highest whereas blue is the lowest expression). **c,d,** Violin plots showing the expression of SARS-CoV-2 interaction partners in 26 adult tissues and 16 fetal tissues respectively. Dots indicating individual data points (cells) are not shown for better visibility of the violin plot. **e,g,** Dotplots show RNA expression of SARS-CoV-2 interaction partners in individual adult (**e**) and fetal (**g**) tissues. **f,h,** Dotplots show comparison of RNA expression of SARS-CoV-2 interaction partners in CNS vs non-CNS cells in adult (**f**) and fetal (**h**) tissues, respectively. **i,j,** *t*-SNE plot showing the expression of SARS-CoV-2 interaction partners in endothelial cells of adult (**i**) and fetal (**j**) tissues from the human cell landscape scRNA-seq dataset^63^. In the *t*-SNE plot, cells are colored according to their degree of expression of SARS-CoV-2 interaction partners (red is the highest whereas blue is the lowest expression). **k,l,** Violin plots showing the expression of SARS-CoV-2 interaction partners in the endothelial cells of 26 adult and 16 fetal tissues, respectively; dots indicate individual data points (cells). **m,o,** Dotplots show RNA expression of SARS-CoV-2 interaction partners in the endothelial cells individual adult (**m**) and fetal (**o**) tissues. **n,p,** Dotplots show comparison of RNA expression of SARS-CoV-2 interaction partners in CNS vs non-CNS endothelial cells in adult (**n**) and fetal (**p**) tissues, respectively.

**Extended Data Figure 9.**
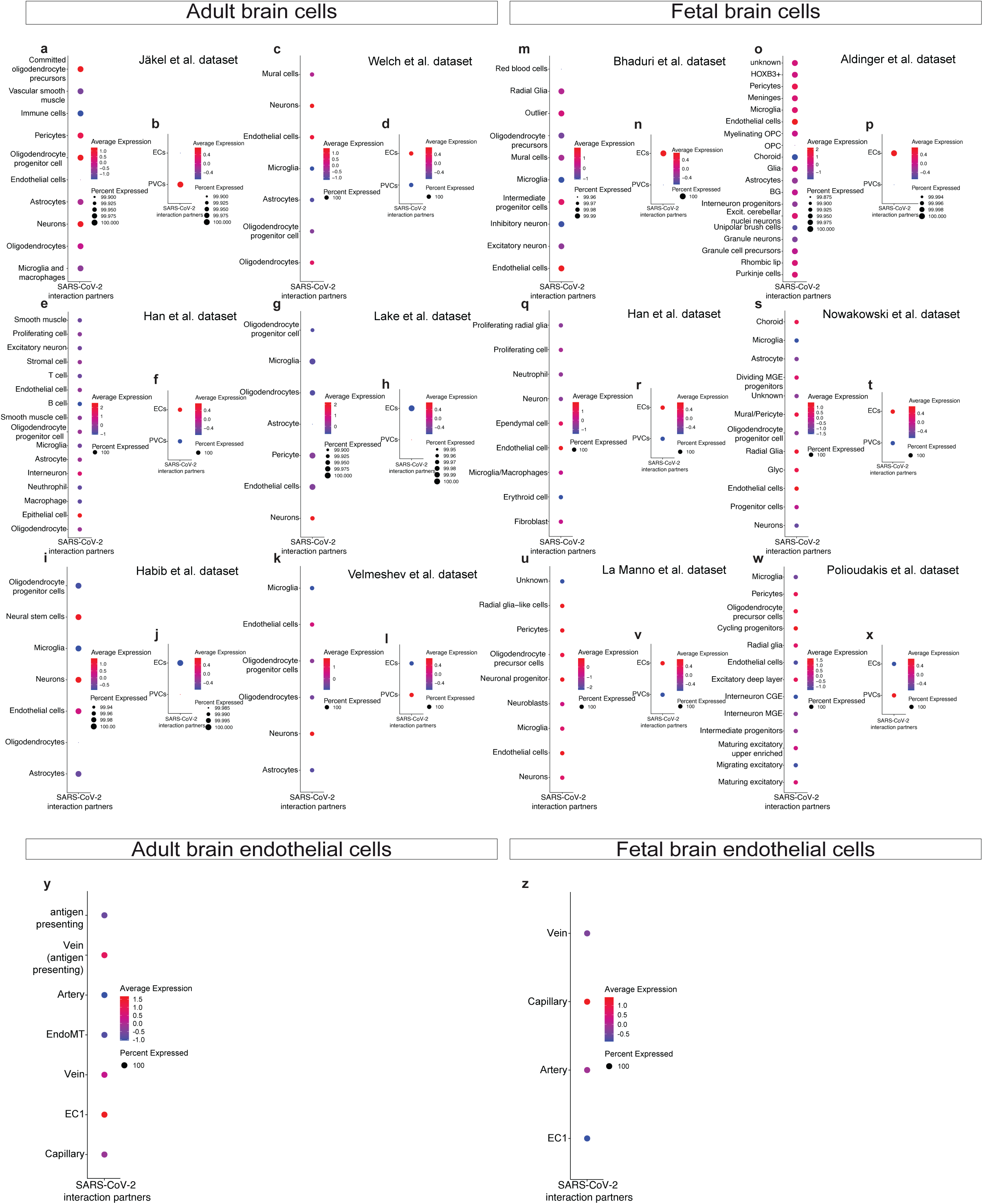
Expression of SARS-Cov-2 interaction partners in adult and fetal human brain cells. **a-l,** Dotplots show RNA expression of SARS-CoV-2 interaction partners in adult brain cells from multiple datasets^63,70–74^. **b,d,f,h,j,l,** Dotplots show comparison of RNA expression of SARS-CoV-2 interaction partners in endothelial cells vs perivascular cells in adult brain single cell RNA-seq datasets. **m-x,** Dotplots show RNA expression of SARS-CoV-2 interaction partners in fetal brain cells from multiple datasets^63,75,110,112–114^. **n,p,r,t,v,x,** Dotplots show comparison of RNA expression of SARS-CoV-2 interaction partners in endothelial cells vs perivascular cells in fetal brain single cell RNA-seq datasets. **y,z,** Dotplots show comparison of RNA expression of SARS-CoV-2 interaction partners in the different endothelial cell types in adult (**y**) and fetal (**z**) brain datasets.

**Extended Data Figure 10.**
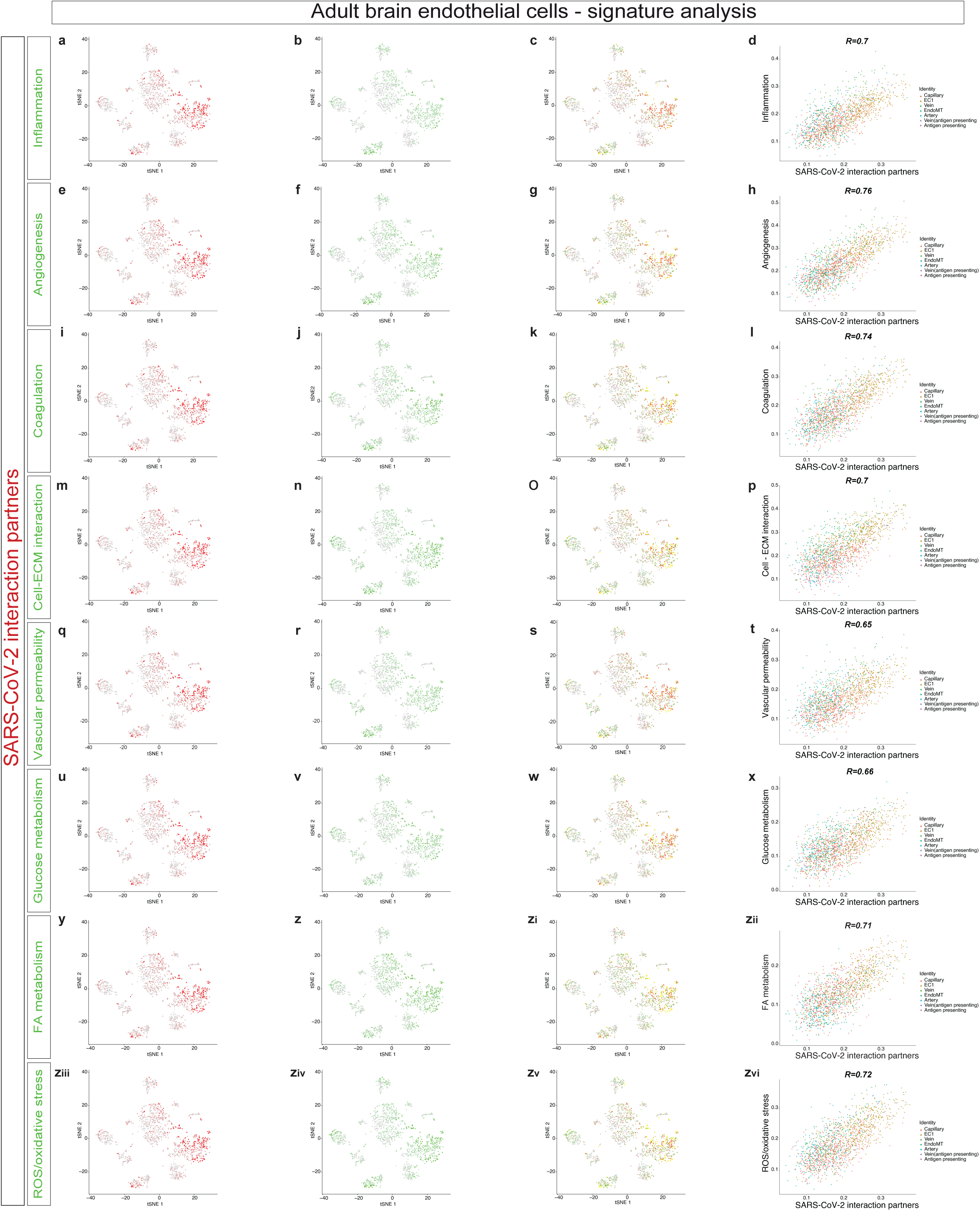
Correlation analysis of SARS-CoV-2 interaction partners with key COVID-19 implicated pathways in adult brain endothelial cells. >**a-zvi,** Co-expression and Pearson correlation analysis of SARS-CoV-2 interaction partners and selected gene signatures in human adult brain endothelial cells. **a-c,** Expression of SARS-CoV-2 interaction partners (red) is co-plotted with inflammation signature (green) in adult brain endothelial cells. **d,** Pearson correlation analysis between the SARS-CoV-2 interaction partners (on the x-axis) and inflammation signature (on the y-axis) done at the single cell level. Correlation coefficient is 0.7 as indicated on the graph. **e-g,** Expression of SARS-CoV-2 interaction partners (red) is co-plotted with angiogenesis signature (green) in adult brain endothelial cells. **h,** Pearson correlation analysis between the SARS-CoV-2 interaction partners (on the x-axis) and angiogenesis signature (on the y-axis) done at the single cell level. Correlation coefficient is 0.76 as indicated on the graph. **i-k,** Expression of SARS-CoV-2 interaction partners (red) is co-plotted with coagulation signature (green) in adult brain endothelial cells. **l,** Pearson correlation analysis between the SARS-CoV-2 interaction partners (on the x-axis) and coagulation signature (on the y-axis) done at the single cell level. Correlation coefficient is 0.74 as indicated on the graph. **m-o,** Expression of SARS-CoV-2 interaction partners (red) is co-plotted with cell-ECM interaction signature (green) in adult brain endothelial cells. **p,** Pearson correlation analysis between the SARS-CoV-2 interaction partners (on the x-axis) and cell-ECM interaction signature (on the y-axis) done at the single cell level. Correlation coefficient is 0.7 as indicated on the graph. **q-s,** Expression of SARS-CoV-2 interaction partners (red) is co-plotted with vascular permeability signature (green) in adult brain endothelial cells. **t,** Pearson correlation analysis between the SARS-CoV-2 interaction partners (on the x-axis) and vascular permeability signature (on the y-axis) done at the single cell level. Correlation coefficient is 0.65 as indicated on the graph. **u-w,** Expression of SARS-CoV2 interaction partners (red) is co-plotted with glucose metabolism signature (green) in adult brain endothelial cells. **x,** Pearson correlation analysis between the SARS-CoV2 interaction partners (on the x-axis) and glucose metabolism signature (on the y-axis) done at the single cell level. Correlation coefficient is 0.66 as indicated on the graph. **y-zi,** Expression of SARS-CoV2 interaction partners (red) is co-plotted with fatty acid metabolism signature (green) in adult brain endothelial cells. **zii,** Pearson correlation analysis between the SARS-CoV2 interaction partners (on the x-axis) and fatty acid metabolism signature (on the y-axis) done at the single cell level. Correlation coefficient is 0.71 as indicated on the graph. **ziii-zv,** Expression of SARS-CoV2 interaction partners (red) is co-plotted with ROS/oxidative stress signature (green) in adult brain endothelial cells. **zvi,** Pearson correlation analysis between the SARS-CoV2 interaction partners (on the x-axis) and ROS/oxidative stress signature (on the y-axis) done at the single cell level. Correlation coefficient is 0.72 as indicated on the graph.

**Extended Data Figure 11.**
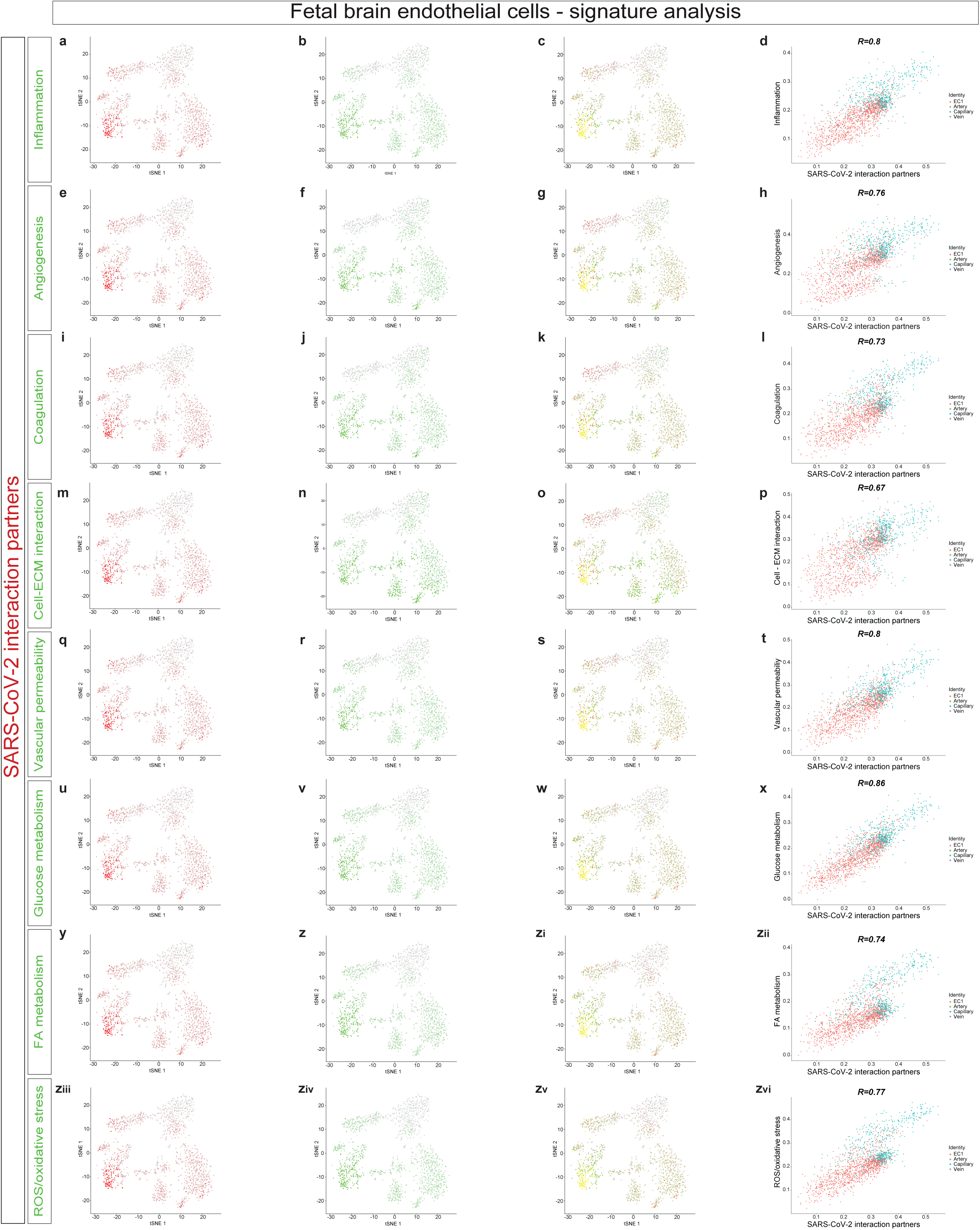
Correlation analysis of SARS-CoV-2 interaction partners with key COVID-19 implicated pathways in fetal brain endothelial cells. **a-zvi,** Co-expression and Pearson correlation analysis of SARS-CoV-2 interaction partners and selected gene signatures in human fetal brain endothelial cells. **a-c,** Expression of SARS-CoV-2 interaction partners (red) is co-plotted with inflammation signature (green) in fetal brain endothelial cells. **d,** Pearson correlation analysis between the SARS-CoV2 interaction partners (on the x-axis) and inflammation signature (on the y-axis) done at the single cell level. Correlation coefficient is 0.8 as indicated on the graph. **e-g,** Expression of SARS-CoV2 interaction partners (red) is co-plotted with angiogenesis signature (green) in fetal brain endothelial cells. **h,** Pearson correlation analysis between the SARS-CoV2 interaction partners (on the x-axis) and angiogenesis signature (on the y-axis) done at the single cell level. Correlation coefficient is 0.76 as indicated on the graph. **i-k,** Expression of SARS-CoV-2 interaction partners (red) is co-plotted with coagulation signature (green) in fetal brain endothelial cells. **l,** Pearson correlation analysis between the SARS-CoV-2 interaction partners (on the x-axis) and coagulation signature (on the y-axis) done at the single cell level. Correlation coefficient is 0.73 as indicated on the graph. **m-o,** Expression of SARS-CoV-2 interaction partners (red) is co-plotted with cell-ECM interaction signature (green) in fetal brain endothelial cells. **p,** Pearson correlation analysis between the SARS-CoV-2 interaction partners (on the x-axis) and cell-ECM interaction signature (on the y-axis) done at the single cell level. Correlation coefficient is 0.67 as indicated on the graph. **q-s,** Expression of SARS-CoV-2 interaction partners (red) is co-plotted with vascular permeability signature (green) in fetal brain endothelial cells. **t,** Pearson correlation analysis between the SARS-CoV-2 interaction partners (on the x-axis) and vascular permeability signature (on the y-axis) done at the single cell level. Correlation coefficient is 0.8 as indicated on the graph. **u-w,** Expression of SARS-CoV2 interaction partners (red) is co-plotted with glucose metabolism signature (green) in fetal brain endothelial cells. **x,** Pearson correlation analysis between the SARS-CoV2 interaction partners (on the x-axis) and glucose metabolism signature (on the y-axis) done at the single cell level. Correlation coefficient is 0.86 as indicated on the graph. **y-zi,** Expression of SARS-CoV2 interaction partners (red) is co-plotted with fatty acid metabolism signature (green) in fetal brain endothelial cells. **zii,** Pearson correlation analysis between the SARS-CoV2 interaction partners (on the x-axis) and fatty acid metabolism signature (on the y-axis) done at the single cell level. Correlation coefficient is 0.74 as indicated on the graph. **ziii-zv,** Expression of SARS-CoV2 interaction partners (red) is co-plotted with ROS/oxidative stress signature (green) in fetal brain endothelial cells. **zvi,** Pearson correlation analysis between the SARS-CoV2 interaction partners (on the x-axis) and ROS/oxidative stress signature (on the y-axis) done at the single cell level. Correlation coefficient is 0.77 as indicated on the graph.

**Extended Data Figure 12.**
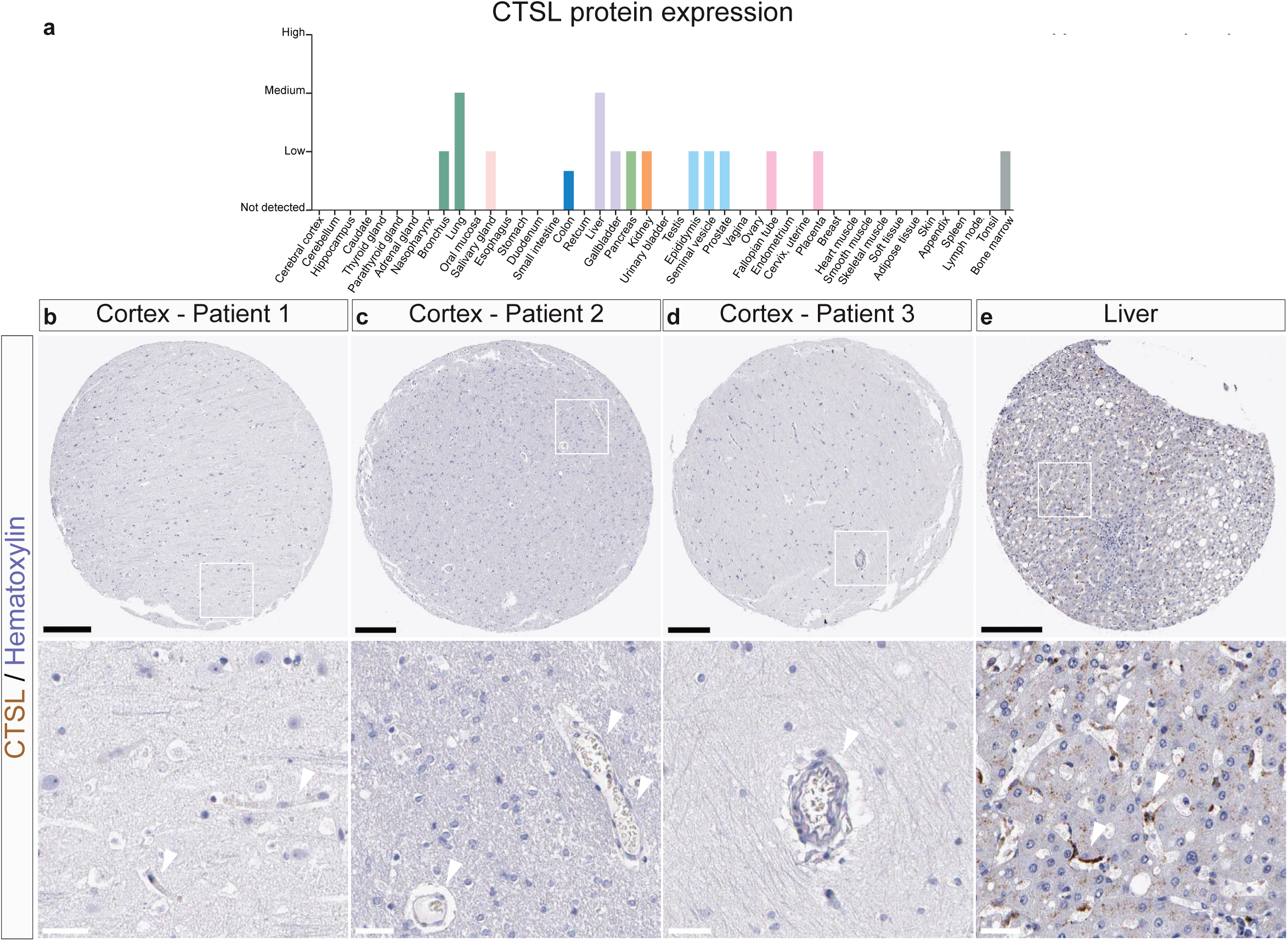
Protein expression of CTSL in situ in CNS and non-CNS tissues. **a,** Protein expression scores for CTSL in human tissues and organs in the human protein atlas revealing no expression in various brain regions and heterogeneous expression scores across various organs. **b-e,** IHC images for the protein expression of CTSL in the brain and in the liver indicating that CTSL is expressed in blood vessels in the liver (arrowheads) but not in the brain (arrowheads). Scale bars: 200 μm for core images, 40 μm for zoomed in images. Arrowheads indicate vascular structures, while arrows indicate positive IHC signals in non-vessel structures.

**Extended Data Figure 13.**
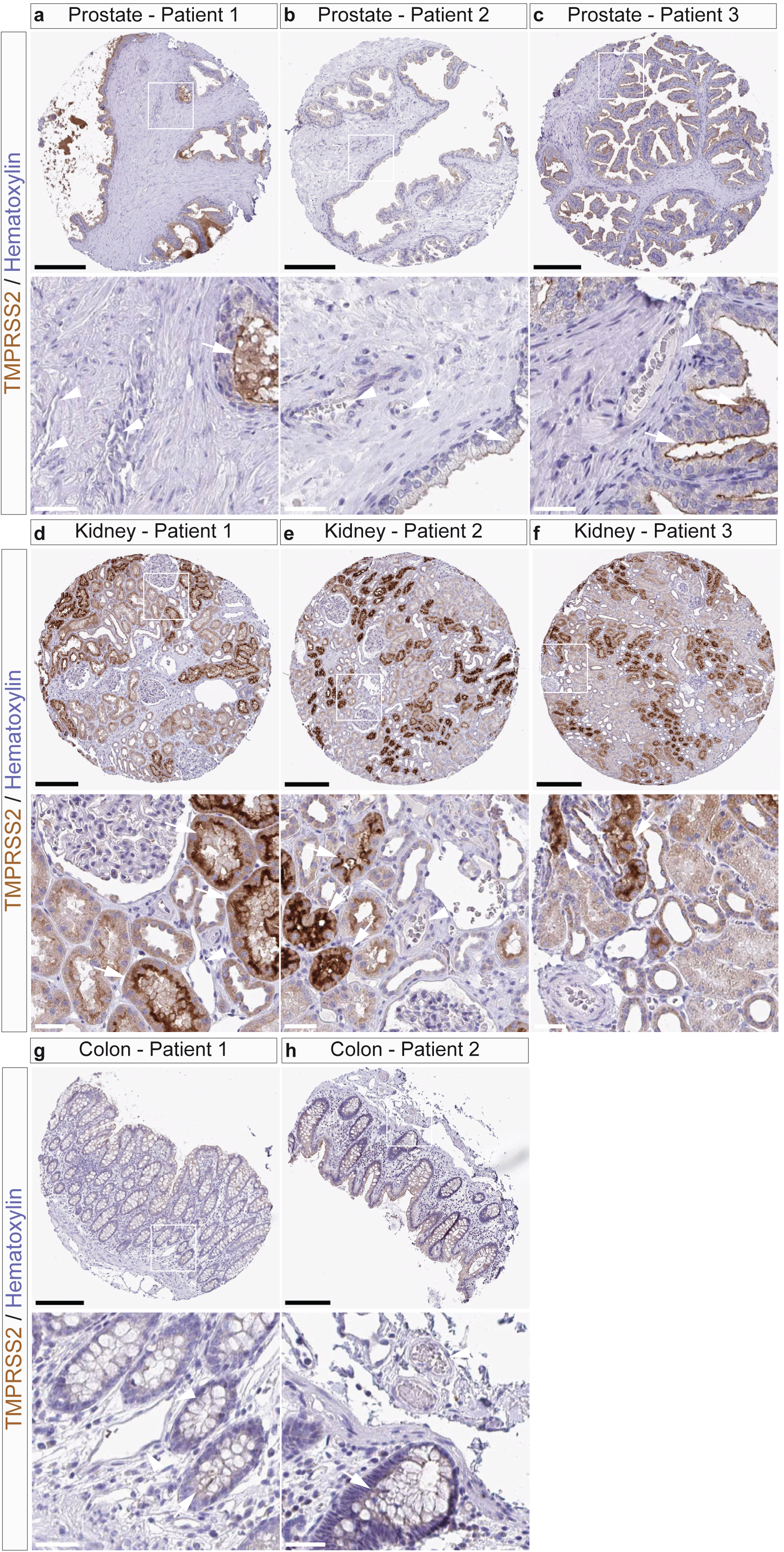
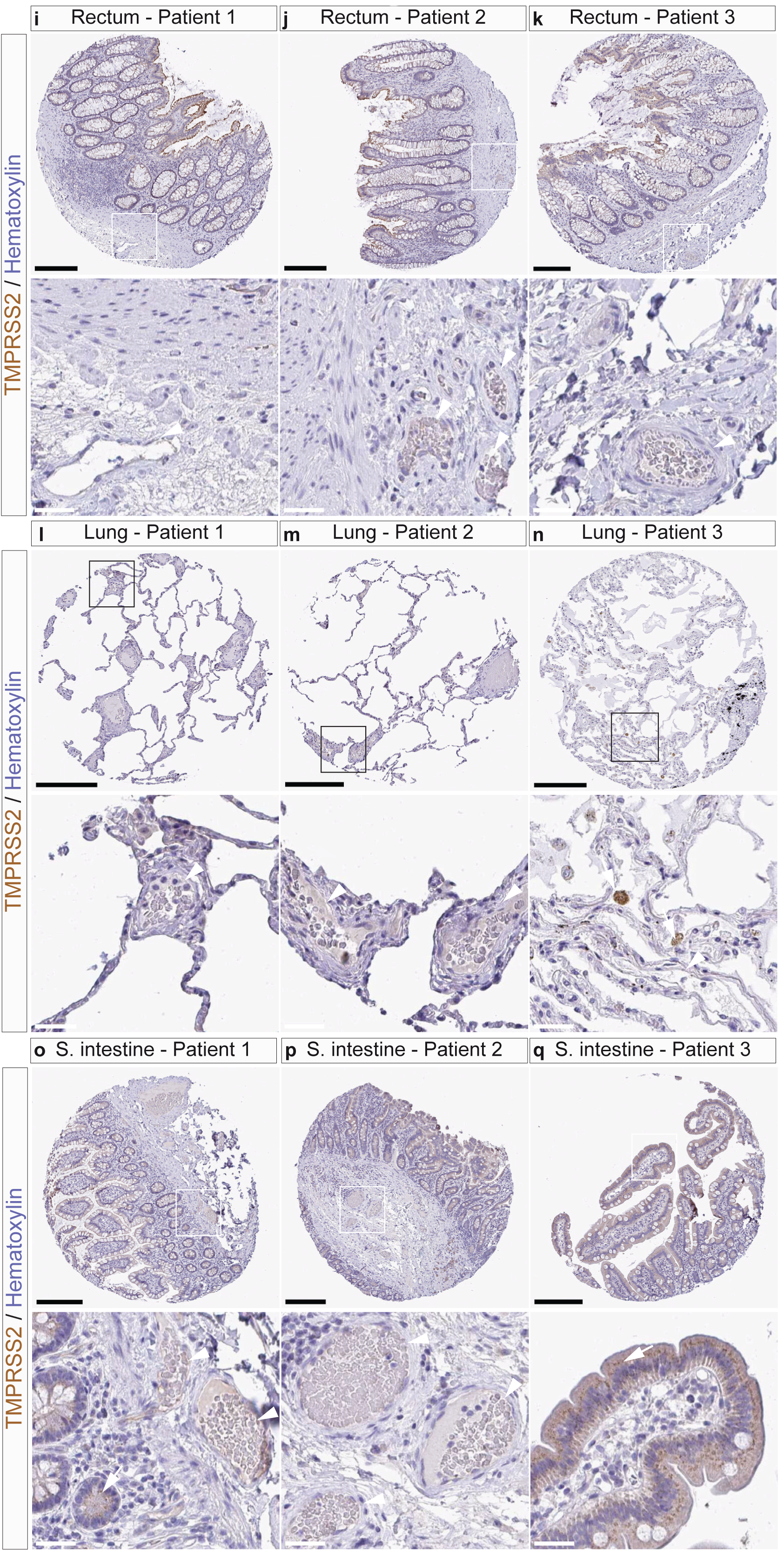
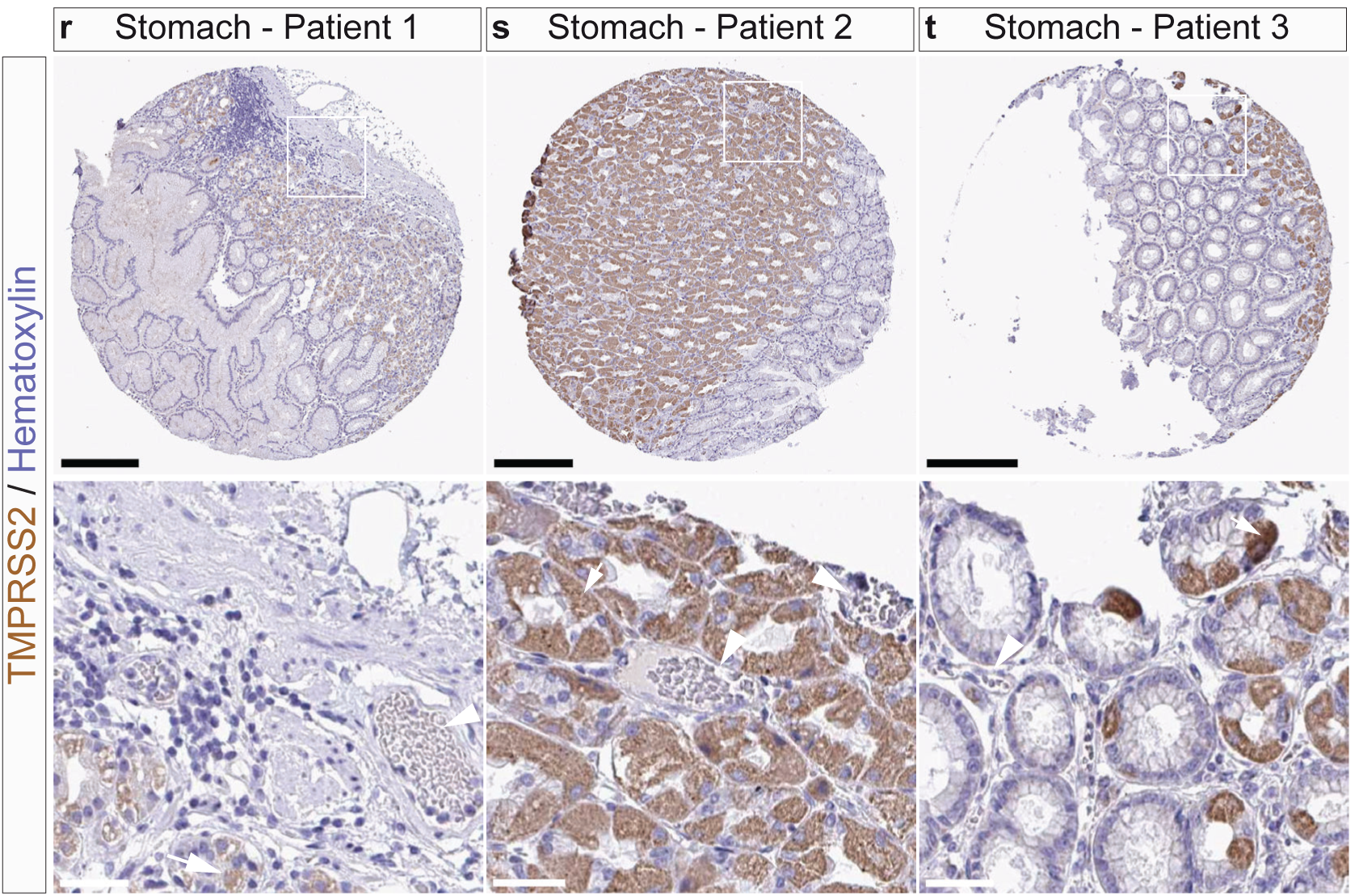
Protein expression of TMPRSS2 in situ in non-CNS tissues. **a-t**, Immunohistochemistry (IHC) images for the protein expression of TMPRSS2 in the various non-CNS tissues/organs indicated. **a-c**, In the prostate, TMPRSS2 is not expressed in blood vessels (arrowheads), but is expressed in the glandular cells (arrows). **d-f**, In the kidney, TMPRSS2 is not expressed in the glomeruli and blood vessels (arrowheads), but is highly expressed in the tubular cells (arrows). **g-h**, In the colon, TMPRSS2 is neither expressed in the blood vessels (arrowheads), nor in the glandular cells (arrows). **i-k**, In the rectum, TMPRSS2 is not expressed in the blood vessels (arrowheads), but is lowly expressed the glandular cells. **l-n**, In the lung, TMPRSS2 is not expressed in the blood vessels (arrowheads), but is expressed in the macrophages (arrows). **o-q**, In the small intestine, TMPRSS2 is not expressed in the blood vessels (arrowheads), but is lowly expressed the glandular cells (arrows). **r-t**, In the stomach, TMPRSS2 is not expressed in the blood vessels (arrowheads) but is expressed the glandular cells (arrows). Scale bars: 200 μm for core images, 40 μm for zoomed in images. Arrows heads indicate vascular structures, while arrows indicate positive IHC signals in non-vascular structures.

**Extended Data Figure 14.**
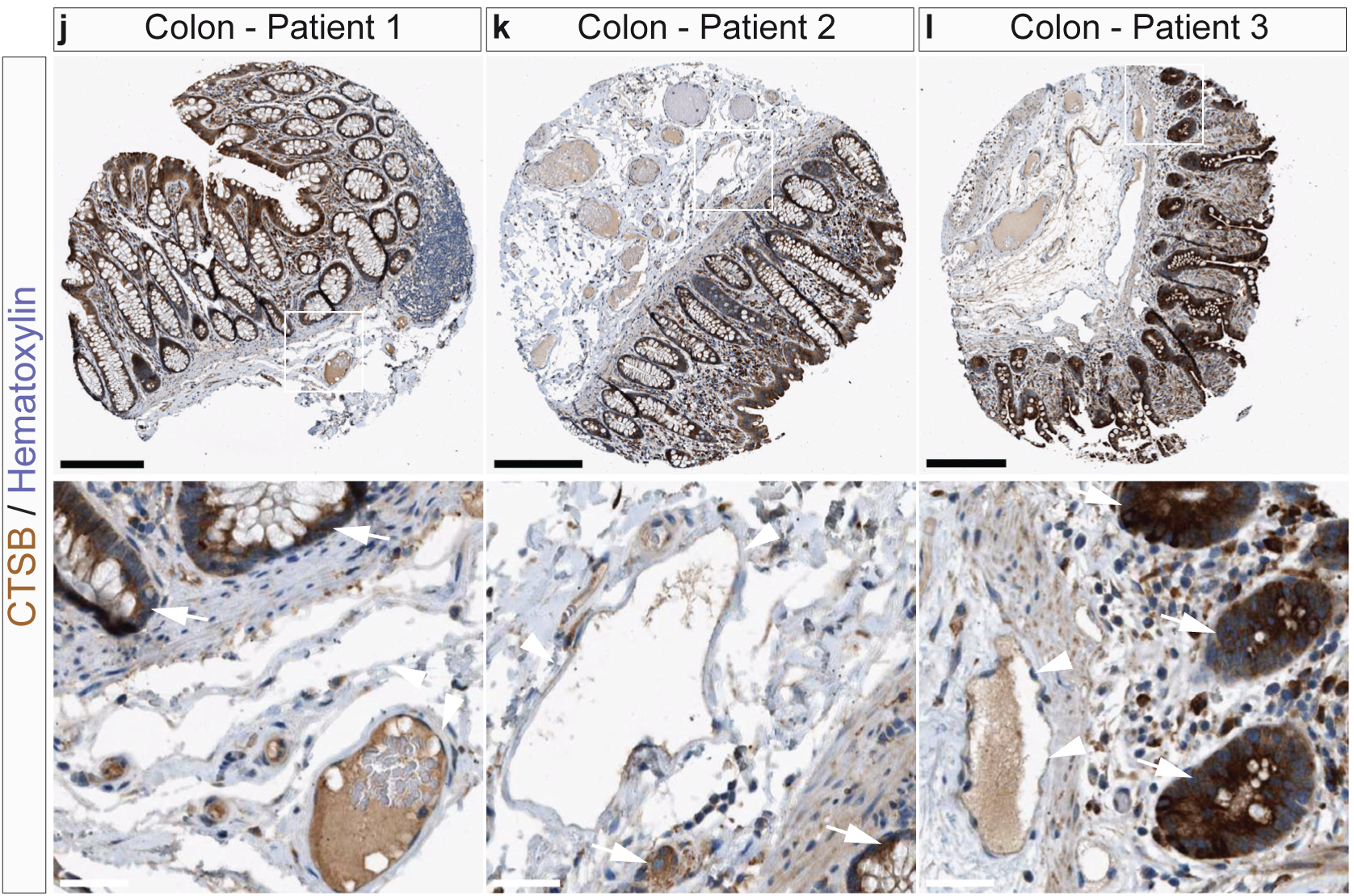
Protein expression of CTSB in situ in non-CNS tissues. **a-l**, Immunohistochemistry (IHC) images for the protein expression of CTSB in the various non-CNS tissues/organs indicated. **a-c**, In the lung, CTSB is slightly expressed in blood vessels (arrowheads) and is highly expressed in the macrophages (arrows). **d-f**, In the heart, CTSB is not expressed in the blood vessels (arrowheads) but is expressed in the myocytes (arrows). **g-i**, In the kidney, CTSB is neither expressed in the blood vessels (arrowheads), nor in the glandular cells (arrows). **j-l**, In the colon, CTSB is not expressed in the blood vessels (arrowheads) but is highly expressed in the glandular cells (arrows). Scale bars: 200 μm for core images, 40 μm for zoomed in images. Arrows heads indicate vascular structures, while arrows indicate positive IHC signals in non-vascular structures.

**Extended Data Figure 15.**
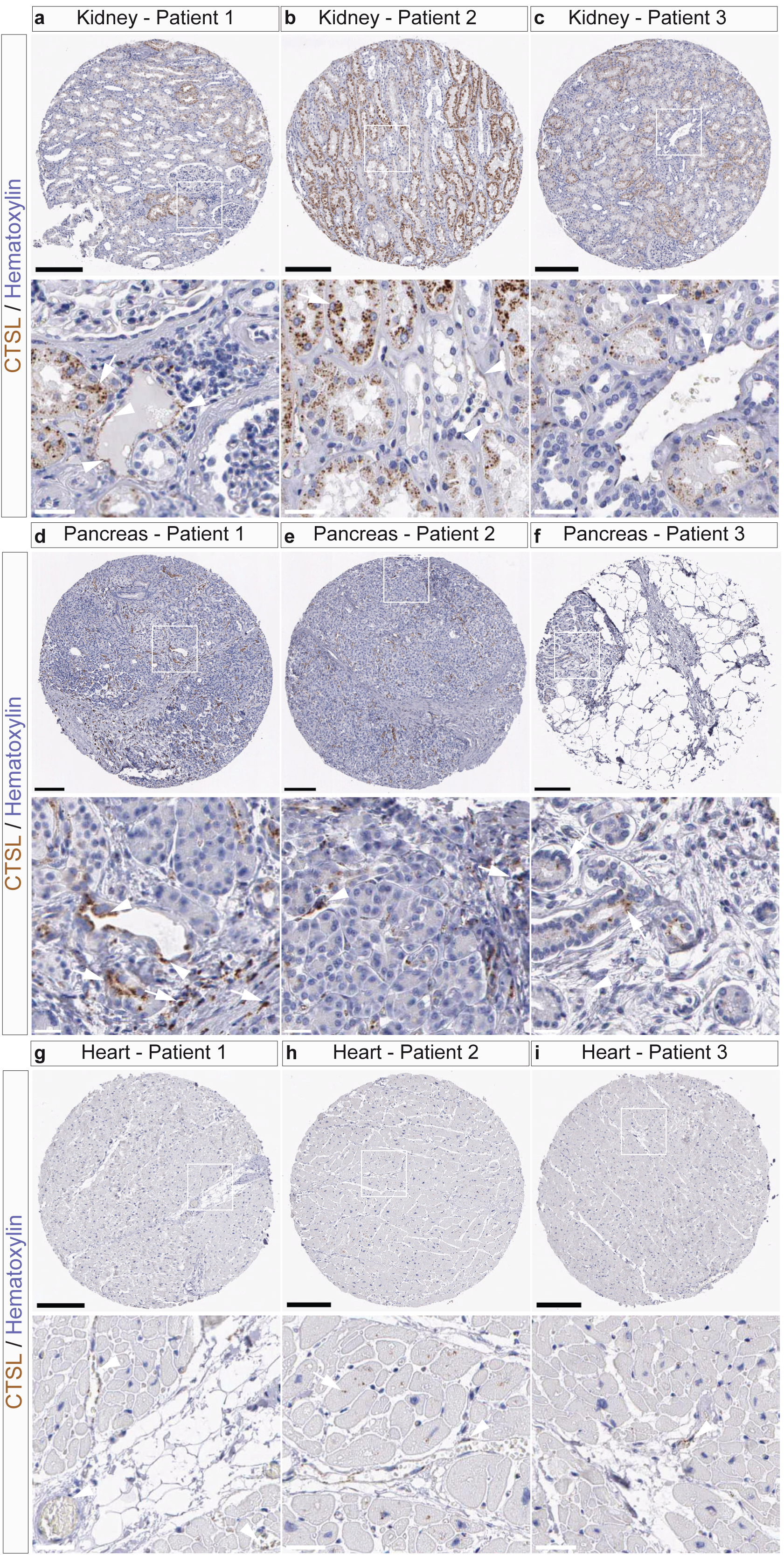
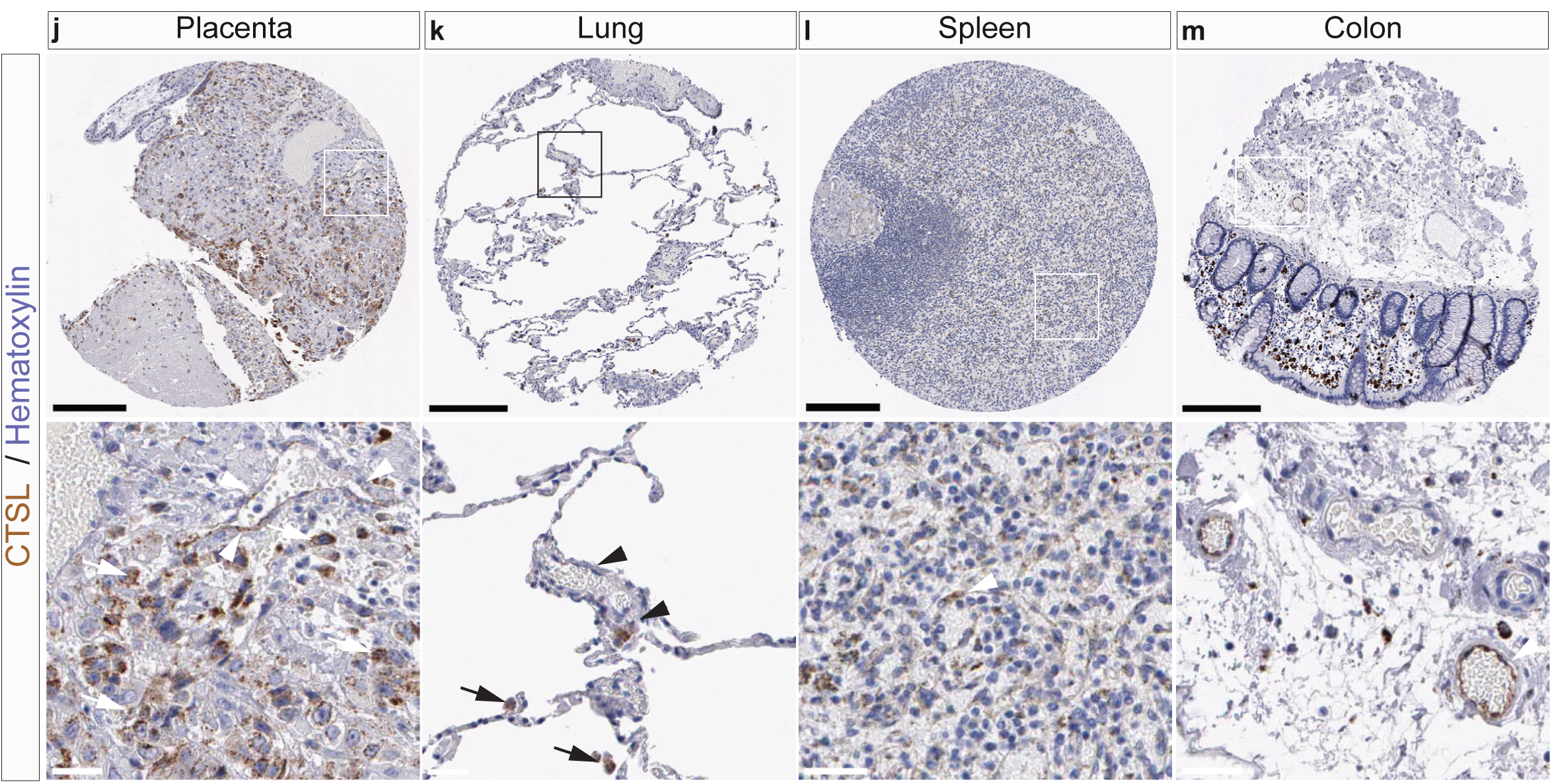
Protein expression of CTSB in situ in non-CNS tissues. **a-m**, Immunohistochemistry (IHC) images for the protein expression of CTSL in the various non-CNS tissues/organs indicated. **a-c**, In the kidney, CTSL is expressed in blood vessels (arrowheads) and is highly expressed in the tubular cells (arrows). **d-f**, In the pancreas, CTSL is expressed in the blood vessels (arrowheads) and in the exocrine glandular cells (arrows). **g-i**, In the heart, CTSL is slightly expressed in the blood vessels (arrowheads) and in the myocytes (arrows). **j**, In the placenta, CTSL is expressed in the blood vessels (arrowheads) and in the decidual/trophoblastic cells (arrows). **k**, In the lung, CTSL is slightly expressed in the blood vessels (arrowheads) and is expressed in macrophages (arrows). **l**, In the spleen, CTSL is expressed in the blood vessels (arrowheads). **m**, In the colon, CTSL is expressed in the blood vessels (arrowheads). Scale bars: 200 μm for core images, 40 μm for zoomed in images. Arrows heads indicate vascular structures, while arrows indicate positive IHC signals in non-vascular structures.

**Extended Data Figure 16.**
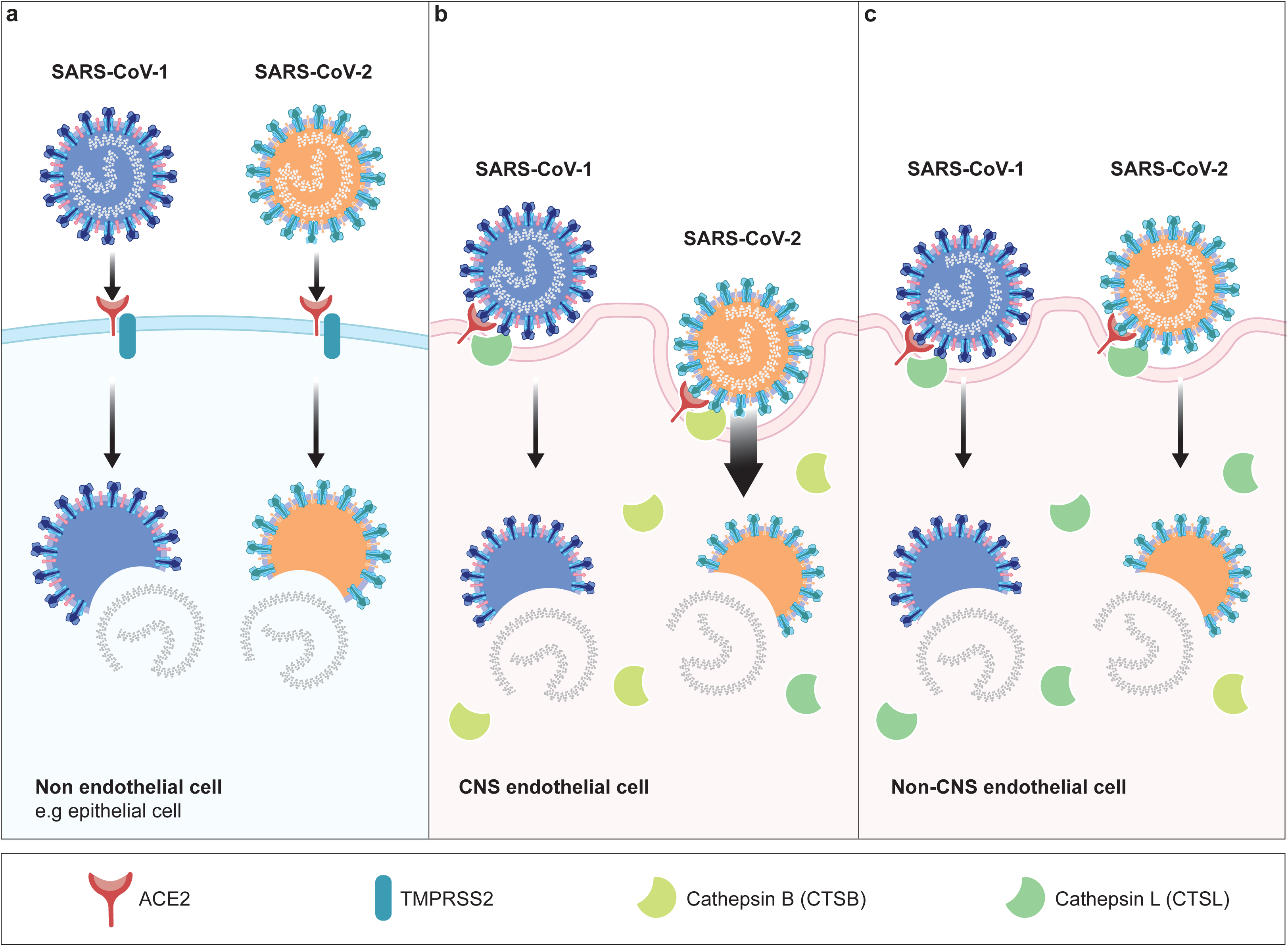
Working model for SARS-CoV-2 entry into CNS- and non-CNS endothelial cells. **a-c,** The schematic illustration of the SARS-CoV-2 entry depicts a non-endothelial (e.g. epithelial cell) which both SARS-CoV-1 and SARS-CoV-2 enter via ACE2 and its co-receptor TMPRSS2 (**a**), a CNS endothelial cell which SARS-CoV-1 enters via ACE2 and CTSL with low efficiency, while SARS-CoV-2 enters via ACE2 and CTSB with a higher efficiency (**b**); and a non-CNS endothelial cell which both SARS-CoV-1 and SARS-CoV-2 enter via ACE2 and CTSL with comparable efficiencies (**c**). In a non-endothelial cell (for instance an epithelial cell), SARS-CoV-1 and SARS-CoV-2 enter via ACE2 and TMPRSS2 (**a**). In the CNS endothelium, CTSB is highly expressed but TMPRSS2 is not, and SARS-CoV-2 cellular entry thus mainly occurs via ACE2 and CTSB, whereas SARS-CoV-1 enters the CNS endothelium via ACE2 and CTSL (**b**). In the non-CNS endothelium, CTSL is highly expressed but TMPRSS2 is not. SARS-CoV-1 and SARS-CoV-2 cellular entry thus mainly occurs via a ACE2 and CTSL (**c**).

## REFERENCES

1. Li, Y. C., Bai, W. Z. & Hashikawa, T. The neuroinvasive potential of SARS-CoV2 may play a role in the respiratory failure of COVID-19 patients. J Med Virol 92, 552–555, doi:10.1002/jmv.25728 (2020).

2. Chen, N. et al. Epidemiological and clinical characteristics of 99 cases of 2019 novel coronavirus pneumonia in Wuhan, China: a descriptive study. Lancet 395, 507–513, doi:10.1016/S0140-6736(20)30211-7 (2020).

3. Zhu, N. et al. A Novel Coronavirus from Patients with Pneumonia in China, 2019. N Engl J Med 382, 727–733, doi:10.1056/NEJMoa2001017 (2020).

4. Nicola, M. et al. The socio-economic implications of the coronavirus pandemic (COVID-19): A review. Int J Surg 78, 185–193, doi:10.1016/j.ijsu.2020.04.018 (2020).

5. Rossman, J. Coronavirus Patients Are Reporting Neurological Symptoms. Here’s What You Need to Know. (2020).

6. Zhou, L., Zhang, M., Wang, J. & Gao, J. Sars-Cov-2: Underestimated damage to nervous system. Travel Med Infect Dis, 101642, doi:10.1016/j.tmaid.2020.101642 (2020).

7. Frank, C. H. M. et al. Guillain-Barre Syndrome Associated with SARS-CoV-2 Infection in a Pediatric Patient. J Trop Pediatr, doi:10.1093/tropej/fmaa044 (2020).

8. Khalifa, M. et al. Guillain-Barre Syndrome Associated with SARS-CoV-2 Detection and a COVID-19 Infection in a Child. J Pediatric Infect Dis Soc, doi:10.1093/jpids/piaa086 (2020).

9. Su, X. W. et al. SARS-CoV-2-associated Guillain-Barre syndrome with dysautonomia. Muscle Nerve 62, E48–E49, doi:10.1002/mus.26988 (2020).

10. Toscano, G. et al. Guillain-Barre Syndrome Associated with SARS-CoV-2. N Engl J Med 382, 2574–2576, doi:10.1056/NEJMc2009191 (2020).

11. Velayos Galan, A., Del Saz Saucedo, P., Peinado Postigo, F. & Botia Paniagua, E. Guillain-Barre syndrome associated with SARS-CoV-2 infection. Neurologia 35, 268–269, doi:10.1016/j.nrl.2020.04.007 (2020).

12. Virani, A. et al. Guillain-Barre Syndrome associated with SARS-CoV-2 infection. IDCases, e00771, doi:10.1016/j.idcr.2020.e00771 (2020).

13. Zhao, H., Shen, D., Zhou, H., Liu, J. & Chen, S. Guillain-Barre syndrome associated with SARS-CoV-2 infection: causality or coincidence? Lancet Neurol 19, 383–384, doi:10.1016/S1474-4422(20)30109-5 (2020).

14. Li, Y. C., Bai, W. Z. & Hashikawa, T. The neuroinvasive potential of SARS-CoV2 may play a role in the respiratory failure of COVID-19 patients. J Med Virol 92, 552–555, doi:10.1002/jmv.25728 (2020).

15. Butowt, R. & Bilinska, K. SARS-CoV-2: Olfaction, Brain Infection, and the Urgent Need for Clinical Samples Allowing Earlier Virus Detection. ACS Chem Neurosci 11, 1200–1203, doi:10.1021/acschemneuro.0c00172 (2020).

16. Ghannam, M. et al. Neurological involvement of coronavirus disease 2019: a systematic review. J Neurol, doi:10.1007/s00415-020-09990-2 (2020).

17. Ellul, M. A. et al. Neurological associations of COVID-19. Lancet Neurol, doi:10.1016/S1474-4422(20)30221-0 (2020).

18. Varatharaj, A. et al. Neurological and neuropsychiatric complications of COVID-19 in 153 patients: a UK-wide surveillance study. Lancet Psychiatry, doi:10.1016/S2215-0366(20)30287-X (2020).

19. Helms, J. et al. Neurologic Features in Severe SARS-CoV-2 Infection. N Engl J Med 382, 2268–2270, doi:10.1056/NEJMc2008597 (2020).

20. Koralnik, I. J. & Tyler, K. L. COVID-19: A Global Threat to the Nervous System. Ann Neurol 88, 1–11, doi:10.1002/ana.25807 (2020).

21. Paterson, R. W. et al. The emerging spectrum of COVID-19 neurology: clinical, radiological and laboratory findings. Brain, doi:10.1093/brain/awaa240 (2020).

22. Tsivgoulis, G. et al. Neurological manifestations and implications of COVID-19 pandemic. Ther Adv Neurol Disord 13, 1756286420932036, doi:10.1177/1756286420932036 (2020).

23. Khateb, M., Bosak, N. & Muqary, M. Coronaviruses and Central Nervous System Manifestations. Front Neurol 11, 715, doi:10.3389/fneur.2020.00715 (2020).

24. Desforges, M., Le Coupanec, A., Brison, E., Meessen-Pinard, M. & Talbot, P. J. Neuroinvasive and neurotropic human respiratory coronaviruses: potential neurovirulent agents in humans. Adv Exp Med Biol 807, 75–96, doi:10.1007/978-81-322-1777-0_6 (2014).

25. Wood, H. New insights into the neurological effects of COVID-19. Nat Rev Neurol, doi:10.1038/s41582-020-0386-7 (2020).

26. Whittaker, A., Anson, M. & Harky, A. Neurological Manifestations of COVID-19: A systematic review and current update. Acta Neurol Scand 142, 14–22, doi:10.1111/ane.13266 (2020).

27. Niazkar, H. R., Zibaee, B., Nasimi, A. & Bahri, N. The neurological manifestations of COVID-19: a review article. Neurol Sci 41, 1667–1671, doi:10.1007/s10072-020-04486-3 (2020).

28. Montalvan, V., Lee, J., Bueso, T., De Toledo, J. & Rivas, K. Neurological manifestations of COVID-19 and other coronavirus infections: A systematic review. Clin Neurol Neurosur 194, doi:ARTN 105921 10.1016/j.clineuro.2020.105921 (2020).

29. Ahmad, I. & Rathore, F. A. Neurological manifestations and complications of COVID-19: A literature review. J Clin Neurosci 77, 8–12, doi:10.1016/j.jocn.2020.05.017 (2020).

30. Zubair, A. S. et al. Neuropathogenesis and Neurologic Manifestations of the Coronaviruses in the Age of Coronavirus Disease 2019: A Review. JAMA Neurol, doi:10.1001/jamaneurol.2020.2065 (2020).

31. Wadman, M., Couzin-Frankel, J., Kaiser, J. & Matacic, C. A Rampage through the Body. Science 368, 356–360 (2020).

32. Khosravani, H., Rajendram, P., Notario, L., Chapman, M. G. & Menon, B. K. Protected Code Stroke: Hyperacute Stroke Management During the Coronavirus Disease 2019 (COVID-19) Pandemic. Stroke 51, 1891–1895, doi:10.1161/STROKEAHA.120.029838 (2020).

33. Mao, L. et al. Neurologic Manifestations of Hospitalized Patients With Coronavirus Disease 2019 in Wuhan, China. Jama Neurol 77, 683–690, doi:10.1001/jamaneurol.2020.1127 (2020).

34. Oxley, T. J. et al. Large-Vessel Stroke as a Presenting Feature of Covid-19 in the Young. New Engl J Med 382, doi:10.1056/NEJMc2009787 (2020).

35. Poyiadji, N. et al. COVID-19-associated Acute Hemorrhagic Necrotizing Encephalopathy: Imaging Features. Radiology 296, E119–E120, doi:10.1148/radiol.2020201187 (2020).

36. Moriguchi, T. et al. A first case of meningitis/encephalitis associated with SARS-Coronavirus-2. Int J Infect Dis 94, 55–58, doi:10.1016/j.ijid.2020.03.062 (2020).

37. Huang, C. L. et al. Clinical features of patients infected with 2019 novel coronavirus in Wuhan, China. Lancet 395, 497–506, doi:10.1016/S0140-6736(20)30183-5 (2020).

38. Walchli, T. et al. Quantitative assessment of angiogenesis, perfused blood vessels and endothelial tip cells in the postnatal mouse brain. Nat Protoc 10, 53–74, doi:10.1038/nprot.2015.002 (2015).

39. Walchli, T. et al. Wiring the Vascular Network with Neural Cues: A CNS Perspective. Neuron 87, 271–296, doi:10.1016/j.neuron.2015.06.038 (2015).

40. Arvanitis, C. D., Ferraro, G. B. & Jain, R. K. The blood-brain barrier and blood-tumour barrier in brain tumours and metastases. Nat Rev Cancer 20, 26–41, doi:10.1038/s41568-019-0205-x (2020).

41. Langen, U. H., Ayloo, S. & Gu, C. H. Development and Cell Biology of the Blood-Brain Barrier. Annu Rev Cell Dev Bi 35, 591–613, doi:10.1146/annurev-cellbio-100617-062608 (2019).

42. Schwartz, D. A. et al. Confirming Vertical Fetal Infection with COVID-19: Neonatal and Pathology Criteria for Early Onset and Transplacental Transmission of SARS-CoV-2 from Infected Pregnant Mothers. Arch Pathol Lab Med, doi:10.5858/arpa.2020-0442-SA (2020).

43. Vivanti, A. J. et al. Transplacental transmission of SARS-CoV-2 infection. Nat Commun 11, 3572, doi:10.1038/s41467-020-17436-6 (2020).

44. Lorenz, N. et al. Neonatal Early-Onset Infection With SARS-CoV-2 in a Newborn Presenting With Encephalitic Symptoms. Pediatr Infect Dis J 39, e212, doi:10.1097/INF.0000000000002735 (2020).

45. Ouldali, N. et al. Emergence of Kawasaki disease related to SARS-CoV-2 infection in an epicentre of the French COVID-19 epidemic: a time-series analysis. Lancet Child Adolesc Health 4, 662–668, doi:10.1016/S2352-4642(20)30175-9 (2020).

46. Moreira, A. Kawasaki disease linked to COVID-19 in children. Nat Rev Immunol 20, 407, doi:10.1038/s41577-020-0350-1 (2020).

47. Ronconi, G. et al. SARS-CoV-2, which induces COVID-19, causes kawasaki-like disease in children: role of pro-inflammatory and anti-inflammatory cytokines. J Biol Regul Homeost Agents 34, 767–773, doi:10.23812/EDITORIAL-RONCONI-E-59 (2020).

48. Xu, S., Chen, M. & Weng, J. COVID-19 and Kawasaki disease in children. Pharmacol Res 159, 104951, doi:10.1016/j.phrs.2020.104951 (2020).

49. Varga, Z. et al. Endothelial cell infection and endotheliitis in COVID-19. Lancet 395, 1417–1418, doi:10.1016/S0140-6736(20)30937-5 (2020).

50. Monteil, V. et al. Inhibition of SARS-CoV-2 Infections in Engineered Human Tissues Using Clinical-Grade Soluble Human ACE2. Cell 181, 905-+, doi:10.1016/j.cell.2020.04.004 (2020).

51. Bao, L. et al. The pathogenicity of SARS-CoV-2 in hACE2 transgenic mice. Nature 583, 830–833, doi:10.1038/s41586-020-2312-y (2020).

52. Zhou, P. et al. A pneumonia outbreak associated with a new coronavirus of probable bat origin. Nature 579, 270-+, doi:10.1038/s41586-020-2012-7 (2020).

53. Hoffmann, M. et al. SARS-CoV-2 Cell Entry Depends on ACE2 and TMPRSS2 and Is Blocked by a Clinically Proven Protease Inhibitor. Cell 181, 271–280 e278, doi:10.1016/j.cell.2020.02.052 (2020).

54. Sungnak, W. et al. SARS-CoV-2 entry factors are highly expressed in nasal epithelial cells together with innate immune genes. Nat Med 26, 681–687, doi:10.1038/s41591-020-0868-6 (2020).

55. Simmons, G. et al. Inhibitors of cathepsin L prevent severe acute respiratory syndrome coronavirus entry. P Natl Acad Sci USA 102, 11876–11881, doi:10.1073/pnas.0505577102 (2005).

56. Glowacka, I. et al. Evidence that TMPRSS2 Activates the Severe Acute Respiratory Syndrome Coronavirus Spike Protein for Membrane Fusion and Reduces Viral Control by the Humoral Immune Response. J Virol 85, 4122–4134, doi:10.1128/Jvi.02232-10 (2011).

57. Matsuyama, S. et al. Efficient Activation of the Severe Acute Respiratory Syndrome Coronavirus Spike Protein by the Transmembrane Protease TMPRSS2. J Virol 84, 12658–12664, doi:10.1128/Jvi.01542-10 (2010).

58. Shulla, A. et al. A Transmembrane Serine Protease Is Linked to the Severe Acute Respiratory Syndrome Coronavirus Receptor and Activates Virus Entry. J Virol 85, 873–882, doi:10.1128/Jvi.02062-10 (2011).

59. Jaimes, J. A., Millet, J. K. & Whittaker, G. R. Proteolytic Cleavage of the SARS-CoV-2 Spike Protein and the Role of the Novel S1/S2 Site. iScience 23, 101212, doi:10.1016/j.isci.2020.101212 (2020).

60. Ferrario, C. M. et al. Effect of angiotensin-converting enzyme inhibition and angiotensin II receptor blockers on cardiac angiotensin-converting enzyme 2. Circulation 111, 2605–2610, doi:10.1161/CIRCULATIONAHA.104.510461 (2005).

61. Hamming, I. et al. Tissue distribution of ACE2 protein, the functional receptor for SARS coronavirus. A first step in understanding SARS pathogenesis. J Pathol 203, 631–637, doi:10.1002/path.1570 (2004).

62. Gordon, D. E. et al. A SARS-CoV-2 protein interaction map reveals targets for drug repurposing. Nature 583, 459–468, doi:10.1038/s41586-020-2286-9 (2020).

63. Han, X. P. et al. Construction of a human cell landscape at single-cell level. Nature 581, 303-+, doi:10.1038/s41586-020-2157-4 (2020).

64. Sungnak, W. et al. SARS-CoV-2 entry factors are highly expressed in nasal epithelial cells together with innate immune genes. Nature Medicine 26, 681-+, doi:10.1038/s41591-020-0868-6 (2020).

65. Zou, X. et al. Single-cell RNA-seq data analysis on the receptor ACE2 expression reveals the potential risk of different human organs vulnerable to 2019-nCoV infection. Front Med-Prc 14, 185–192, doi:10.1007/s11684-020-0754-0 (2020).

66. Qi, F. R., Qian, S., Zhang, S. Y. & Zhang, Z. Single cell RNA sequencing of 13 human tissues identify cell types and receptors of human coronaviruses. Biochem Bioph Res Co 526, 135–140, doi:10.1016/j.bbrc.2020.03.044 (2020).

67. Muoio, V., Persson, P. B. & Sendeski, M. M. The neurovascular unit - concept review. Acta Physiol 210, 790–798, doi:10.1111/apha.12250 (2014).

68. Hawkins, B. T. & Davis, T. P. The blood-brain barrier/neurovascular unit in health and disease. Pharmacol Rev 57, 173–185, doi:10.1124/pr.57.2.4 (2005).

69. Iadecola, C. The Neurovascular Unit Coming of Age: A Journey through Neurovascular Coupling in Health and Disease. Neuron 96, 17–42, doi:10.1016/j.neuron.2017.07.030 (2017).

70. Jakel, S. et al. Altered human oligodendrocyte heterogeneity in multiple sclerosis. Nature 566, 543–547, doi:10.1038/s41586-019-0903-2 (2019).

71. Welch, J. D. et al. Single-Cell Multi-omic Integration Compares and Contrasts Features of Brain Cell Identity. Cell 177, 1873–1887 e1817, doi:10.1016/j.cell.2019.05.006 (2019).

72. Lake, B. B. et al. Integrative single-cell analysis of transcriptional and epigenetic states in the human adult brain. Nat Biotechnol 36, 70–80, doi:10.1038/nbt.4038 (2018).

73. Habib, N. et al. Massively parallel single-nucleus RNA-seq with DroNc-seq. Nat Methods 14, 955–958, doi:10.1038/nmeth.4407 (2017).

74. Velmeshev, D. et al. Single-cell genomics identifies cell type-specific molecular changes in autism. Science 364, 685–689, doi:10.1126/science.aav8130 (2019).

75. La Manno, G. et al. Molecular Diversity of Midbrain Development in Mouse, Human, and Stem Cells. Cell 167, 566–580 e519, doi:10.1016/j.cell.2016.09.027 (2016).

76. Song, E. et al. Neuroinvasion of SARS-CoV-2 in human and mouse brain. bioRxiv, doi:10.1101/2020.06.25.169946 (2020).

77. Cakir, B. et al. Engineering of human brain organoids with a functional vascular-like system. Nat Methods 16, 1169–1175, doi:10.1038/s41592-019-0586-5 (2019).

78. Vanlandewijck, M. et al. A molecular atlas of cell types and zonation in the brain vasculature. Nature 554, 475–480, doi:10.1038/nature25739 (2018).

79. Kalucka, J. et al. Single-Cell Transcriptome Atlas of Murine Endothelial Cells. Cell 180, 764–779 e720, doi:10.1016/j.cell.2020.01.015 (2020).

80. Goveia, J. et al. An Integrated Gene Expression Landscape Profiling Approach to Identify Lung Tumor Endothelial Cell Heterogeneity and Angiogenic Candidates. Cancer Cell 37, 421, doi:10.1016/j.ccell.2020.03.002 (2020).

81. Ackermann, M., et al. Pulmonary Vascular Endothelialitis, Thrombosis, and Angiogenesis in Covid-19. N Engl J Med 383, 120–128, doi:10.1056/NEJMoa2015432 (2020).

82. Teuwen, L. A., Geldhof, V., Pasut, A. & Carmeliet, P. COVID-19: the vasculature unleashed. Nat Rev Immunol, doi:10.1038/s41577-020-0343-0 (2020).

83. Perisic, L. et al. Profiling of Atherosclerotic Lesions by Gene and Tissue Microarrays Reveals PCSK6 as a Novel Protease in Unstable Carotid Atherosclerosis. Arterioscl Throm Vas 33, 2432–2443, doi:10.1161/Atvbaha.113.301743 (2013).

84. Kryczka, J. et al. Matrix Metalloproteinase-2 Cleavage of the beta 1 Integrin Ectodomain Facilitates Colon Cancer Cell Motility. J Biol Chem 287, 36556–36566, doi:10.1074/jbc.M112.384909 (2012).

85. Wang, H., Chen, Y. L., Fernandez-Del Castillo, C., Yilmaz, O. & Deshpande, V. Heterogeneity in signaling pathways of gastroenteropancreatic neuroendocrine tumors: a critical look at notch signaling pathway. Modern Pathol 26, 139–147, doi:10.1038/modpathol.2012.143 (2013).

86. Zhao, X. L., Eyo, U. B., Murugan, M. & Wu, L. J. Microglial interactions with the neurovascular system in physiology and pathology. Dev Neurobiol 78, 604–617, doi:10.1002/dneu.22576 (2018).

87. Dudvarski Stankovic, N., Teodorczyk, M., Ploen, R., Zipp, F. & Schmidt, M. H. H. Microglia-blood vessel interactions: a double-edged sword in brain pathologies. Acta Neuropathol 131, 347–363, doi:10.1007/s00401-015-1524-y (2016).

88. Ding, Y. et al. The clinical pathology of severe acute respiratory syndrome (SARS): a report from China. J Pathol 200, 282–289, doi:10.1002/path.1440 (2003).

89. Stahl, K., Brasen, J. H., Hoeper, M. M. & David, S. Direct evidence of SARS-CoV-2 in gut endothelium. Intensive Care Med, doi:10.1007/s00134-020-06237-6 (2020).

90. Hoegen, T. et al. The NLRP3 Inflammasome Contributes to Brain Injury in Pneumococcal Meningitis and Is Activated through ATP-Dependent Lysosomal Cathepsin B Release. J Immunol 187, 5440–5451, doi:10.4049/jimmunol.1100790 (2011).

91. Moon, H. Y. et al. Running-Induced Systemic Cathepsin B Secretion Is Associated with Memory Function. Cell Metabolism 24, 332–340, doi:10.1016/j.cmet.2016.05.025 (2016).

92. vanderStappen, J. W. J., Williams, A. C., Maciewicz, R. A. & Paraskeva, C. Activation of cathepsin B, secreted by a colorectal cancer cell line requires low pH and is mediated by cathepsin D. Int J Cancer 67, 547–554, doi:Doi 10.1002/(Sici)1097-0215(19960807)67:4<547::Aid-Ijc14>3.0.Co;2-4 (1996).

93. Pavlova, A. & Bjork, I. Grafting of features of cystatins C or B into the N-terminal region or second binding loop of cystatin A (stefin A) substantially enhances inhibition of cysteine proteinases. Biochemistry-Us 42, 11326–11333, doi:10.1021/bi030119v (2003).

94. Estrada, S. et al. The role of Gly-4 of human cystatin A (Stefin A) in the binding of target proteinases. Characterization by kinetic and equilibrium methods of the interactions of cystatin a Gly-4 mutants with papain, cathepsin B, and cathepsin L. Biochemistry-Us 37, 7551–7560, doi:DOI 10.1021/bi980026r (1998).

95. Fais, S. Cannibalism: A way to feed on metastatic tumors. Cancer Lett 258, 155–164, doi:10.1016/j.canlet.2007.09.014 (2007).

96. Grundmann, U., Nerlich, C., Rein, T. & Zettlmeissl, G. Complete cDNA sequence encoding human beta-galactoside alpha-2,6-sialyltransferase. Nucleic Acids Res 18, 667, doi:10.1093/nar/18.3.667 (1990).

97. Bast, B. J. et al. The HB-6, CDw75, and CD76 differentiation antigens are unique cell-surface carbohydrate determinants generated by the beta-galactoside alpha 2,6- sialyltransferase. J Cell Biol 116, 423–435, doi:10.1083/jcb.116.2.423 (1992).

98. Lee, M. et al. Transcriptional programs of lymphoid tissue capillary and high endothelium reveal control mechanisms for lymphocyte homing. Nat Immunol 15, 982–U129, doi:10.1038/ni.2983 (2014).

99. Li, X., Sun, X. & Carmeliet, P. Hallmarks of Endothelial Cell Metabolism in Health and Disease. Cell Metab 30, 414–433, doi:10.1016/j.cmet.2019.08.011 (2019).

100. Gnirss, K. et al. Cathepsins B and L activate Ebola but not Marburg virus glycoproteins for efficient entry into cell lines and macrophages independent of TMPRSS2 expression. Virology 424, 3–10, doi:10.1016/j.virol.2011.11.031 (2012).

101. Felbor, U. et al. Neuronal loss and brain atrophy in mice lacking cathepsins B and L. Proc Natl Acad Sci U S A 99, 7883–7888, doi:10.1073/pnas.112632299 (2002).

102. Yoshida, M. et al. Primate neurons show different vulnerability to transient ischemia and response to cathepsin inhibition. Acta Neuropathologica 104, 267–272, doi:10.1007/s00401-002-0554-4 (2002).

103. Hook, G., Jacobsen, J. S., Grabstein, K., Kindy, M. & Hook, V. Cathepsin B is a New Drug Target for Traumatic Brain injury Therapeutics: evidence for E64d as a Promising Lead Drug Candidate. Frontiers in Neurology 6, doi:ARTN 17810.3389/fneur.2015.00178 (2015).

104. Nakanishi, H. Microglial cathepsin B as a key driver of inflammatory brain diseases and brain aging. Neural Regen Res 15, 25–29, doi:Pmid 3153563810.4103/1673-5374.264444 (2020).

105. Wu, Q. Q. et al. Cathepsin B deficiency attenuates cardiac remodeling in response to pressure overload via TNF-alpha/ASK1/JNK pathway. Am J Physiol-Heart C 308, H1143–H1154, doi:10.1152/ajpheart.00601.2014 (2015).

106. Cocchiaro, P. et al. The Multifaceted Role of the Lysosomal Protease Cathepsins in Kidney Disease. Front Cell Dev Biol 5, doi:UNSP 11410.3389/fcell.2017.00114 (2017).

107. Xu, Q., Mariman, E. C. M., Goossens, G. H., Blaak, E. E. & Jocken, J. W. E. Cathepsin gene expression in abdominal subcutaneous adipose tissue of obese/overweight humans. Adipocyte 9, 246–252, doi:10.1080/21623945.2020.1775035 (2020).

108. Potente, M. & Makinen, T. Vascular heterogeneity and specialization in development and disease. Nat Rev Mol Cell Bio 18, 477–494, doi:10.1038/nrm.2017.36 (2017).

109. Baig, A. M., Khaleeq, A., Ali, U. & Syeda, H. Evidence of the COVID-19 Virus Targeting the CNS: Tissue Distribution, Host-Virus Interaction, and Proposed Neurotropic Mechanisms. Acs Chemical Neuroscience 11, 995–998, doi:10.1021/acschemneuro.0c00122 (2020).

110. Bhaduri, A. et al. Cell stress in cortical organoids impairs molecular subtype specification. Nature 578, 142–148, doi:10.1038/s41586-020-1962-0 (2020).

111. Han, X. et al. Mapping the Mouse Cell Atlas by Microwell-Seq. Cell 173, 1307, doi:10.1016/j.cell.2018.05.012 (2018).

112. Aldinger. Spatial and single-cell transcriptional landscape of human cerebellar development. biorxiv (2020).

113. Nowakowski, T. J. et al. Spatiotemporal gene expression trajectories reveal developmental hierarchies of the human cortex. Science 358, 1318–1323, doi:10.1126/science.aap8809 (2017).

114. Polioudakis, D. et al. A Single-Cell Transcriptomic Atlas of Human Neocortical Development during Mid-gestation. Neuron 103, 785-+, doi:10.1016/j.neuron.2019.06.011 (2019).

115. Korsunsky, I. et al. Fast, sensitive and accurate integration of single-cell data with Harmony. Nature Methods 16, 1289-+, doi:10.1038/s41592-019-0619-0 (2019).

116. Tabula Muris, C. et al. Single-cell transcriptomics of 20 mouse organs creates a Tabula Muris. Nature 562, 367–372, doi:10.1038/s41586-018-0590-4 (2018).

